# Paralysis Efficiency (ED_50_) Scales Linearly with Lethality (LD_50_) in Spider Venoms

**DOI:** 10.64898/2026.03.06.710087

**Authors:** Keith Lyons, Dayle Leonard, Leona McSharry, Michael Martindale, Brandon L. Collier, Aiste Vitkauskaite, John P. Dunbar, Michel M. Dugon, Kevin Healy

**Author notes:** These authors contributed equally to this work (Keith Lyons, Dayle Leonard, Michel M. Dugon, Kevin Healy).

## Abstract

Historically, venom potencies have been assessed using measures of lethality, such as the median lethal dose (LD_50_). However, venoms may be selected primarily for their ability to rapidly incapacitate rather than cause mortality, meaning LD_50_ may not capture the efficacy of venoms in an ecological and evolutionary context. To capture this context, recent studies have adapted measures that assess venoms’ ability to rapidly incapacitate, such as the median effective dose (ED_50_). However, while ED_50_ values are expected to provide a more proximate assessment of ecological variation in venom potency, it is unknown whether historically available LD_50_ values are still useful proxies of ecologically relevant potency or whether they capture independent axes of venom variation. Here, we test the relationship between LD_50_ and ED_50_ in spider venoms by experimentally estimating LD_50_ and ED_50_ for 12 species and collating additional potency data for 40 species retrieved from the literature. We observed an isometric relationship between LD_50_ and ED_50_ in both analyses, showing these potency measures are both strongly coupled, with an increase in paralysis efficiency associated with a similar increase in lethality. Our results suggest that the functional aspects of venom potency, paralysis and lethality, are intrinsically linked, and due to this strong mechanistic coupling, historically available LD_50_ values may be used to compare general venom potencies in spiders, provided that they are based on the same prey model.

## 1. Introduction

Venom has evolved independently across the Animal Kingdom more than 100 times, in at least eight phyla and is involved in at least 14 distinct ecological roles (Schendel *et al*. 2019; Hayes *et al*. 2025). Predation is one of the more ubiquitous roles, functioning as the primary role of venoms in cephalopods, centipedes, cnidarians, cone snails, heteropterans, scorpions, snakes and spiders, among others (Arbuckle 2017; Schendel *et al*. 2019; Hayes *et al*. 2025; Herzig 2025; Jenner *et al*. 2025). Animals with predation-oriented venoms have evolved toxins that function both individually and synergistically to manipulate important physiological and signalling processes in their prey, resulting in non-lethal, detrimental effects or death while facilitating consumption (Arbuckle 2017; Kuhn-Nentwig *et al*. 2019; Schendel *et al*. 2019; Jenner *et al*. 2025). These toxins employ various mechanisms to subdue prey, including haemotoxins that enhance or disrupt blood clotting factors and lyse erythrocytes, necrotoxins (i.e., cytotoxins) that cause focal to locally extensive tissue necrosis, and neurotoxins that target the nervous system and induce both rapid and prolonged paralytic effects (Kuhn-Nentwig *et al*. 2019; Bolon *et al*. 2023). While these toxins can cause prey mortalities (Herrera *et al*. 2012; Brinkman *et al*. 2014), it is hypothesised that predation-oriented toxins are selected primarily for their ability to rapidly incapacitate prey (Arbuckle 2017; Michálek *et al*. 2024). This is because achieving prey mortality in an ecologically relevant timeframe may require more venom than is required to incapacitate, incurring higher metabolic costs, whereas rapid paralysis is often sufficient to facilitate mechanical dispatch and the consumption of prey (Wigger *et al*. 2002; Morgenstern and King 2013; Arbuckle 2017). However, classical potency measures have focused primarily on lethality, potentially missing ecologically relevant patterns associated with venom potency.

To better understand how venom potency has evolved, researchers use various measures which typically involve introducing a known amount of venom or specific venom fractions, serially diluted to multiple concentrations, into a prey model and recording the resulting effects (Finney 1985). Historically, the most common measure of venom potency is the median lethal dose (Finney 1985) or LD_50_, the amount of venom required to cause a 50% mortality rate in a prey model cohort (Bücherl 1946; Edwards 1961; McCrone 1964; Sutherland 1972; Norment and Foil 1980; Hillyard *et al*. 1989; Celerier *et al*. 1993; Nagai *et al*. 2000; Nunan *et al*. 2001; Wigger *et al*. 2002; Bhakuni and Rawat 2005; Gentz *et al*. 2009; de Roodt *et al*. 2017; Valenzuela-Rojas *et al*. 2019; Lyons *et al*. 2020; Lyons *et al*. 2025). Although many venoms can cause mortality in prey models, it typically takes considerably longer than the initial paralysis, suggesting the mortality rates recorded during LD_50_ assays may not be ecologically relevant (Michálek *et al*. 2024). Consequently, recent studies have adapted alternative measures such as the median effective dose (Pearce *et al*. 1995) or ED_50_ (McElroy *et al*. 2017; Michálek *et al*. 2019; Michálek *et al*. 2022; Lyons *et al*. 2023) which measures the amount of venom required to incapacitate 50% of a prey model cohort. While LD_50_ typically gauges the capacity of venoms to cause prey mortality within 24 hours (Michálek *et al*. 2024), ED_50_ usually gauges their ability to incapacitate prey within more ecologically relevant timeframes, which vary between studies—30 minutes (Rayner *et al*. 2022), one hour (Michálek *et al*. 2019), four hours (Lyons *et al*. 2023), five hours (Manzoli-Palma *et al*. 2003) and rarely more than 24 hours (Herzig and Hodgson 2008). Hence, ED_50_ is expected to be better suited for detecting novel, ecologically relevant aspects of venom potency. However, while ED_50_ values may capture more ecologically relevant variation, they are not as taxonomically widespread in the literature or as frequent as lethality measures, such as LD_50_, raising the question of whether such historically common measures should be used in comparative studies addressing ecological and evolutionary questions. The use of such historical lethality measures to better understand ecological patterns in venom potency may thus be dependent on whether the ability of venoms to incapacitate prey quickly is generally decoupled from subsequent lethality, or coupled, with selection for an increase in paralysis efficiency resulting in a similar increase in lethality.

In the context of evolutionary biology, functional coupling is typically associated with a physiological trait or set of traits that perform multiple, overlapping functions (Farina *et al*. 2019). For example, the elytra of beetles are modified forewings used in defence, flight and a plethora of other functions (Linz *et al*. 2016; Goczał and Beutel 2023), while the cranial components of teleost fish are functionally coordinated to perform the overlapping functions of suction feeding and gill ventilation (Farina *et al*. 2019). However, not all traits or functions in functional coupling are directly selected for. For example, species that change their body sizes due to selection are likely to display changes in the top speed at which they can move, not necessarily due to direct selection for speed but more likely due to the physiological constraints linked with size (Mezzini *et al*. 2025). Likewise, different functional aspects of venom potency, such as those measured by ED_50_ and LD_50_, may be tightly linked due to strong mechanistic coupling. For example, neurotoxins selected for their ability to rapidly incapacitate prey (Escoubas *et al*. 2000; Osipov and Utkin 2015; Gao *et al*. 2017) may demonstrate strong links with mortality as paralytic effects often involve the disruption of vital functions which can ultimately lead to organ failure (Langenegger *et al*. 2019; Bolon *et al*. 2023). However, such immediate effects of venoms may not necessarily be linked to the later onset of mortality, with many venom fractions inducing rapid but reversible paralysis in prey (Loret *et al*. 1992; Adams 2004; Michálek *et al*. 2022). Hence, determining whether there is a relationship between ED_50_ and LD_50_ values, thereby implying a link between the functions they measure, would not only help ascertain the potential value of historic LD_50_ values for inferring ecological patterns in venoms but would also improve our understanding of the general mechanisms underlying the functionality of predation-oriented venoms.

Through the combination of a smaller-scale experimental analysis and a larger-scale phylogenetic comparative analysis, we test the relationship between LD_50_ and ED_50_ for 50 spider species spanning 26 families. While we expect ED_50_ values to be far lower than their corresponding LD_50_ values, as less venom is typically required to achieve prey paralysis compared to prey mortality (Wigger *et al*. 2002; Morgenstern and King 2013), we predict that if LD_50_ and ED_50_ are functionally coupled, they will scale isometrically with a slope close to one. Alternatively, if LD_50_ and ED_50_ are functionally decoupled, they will instead have a sublinear relationship, with expected LD_50_ efficiencies decreasing faster than their respective ED_50_ efficiencies in the scenario where there are higher selective pressures for prey paralysis compared to lethality.

## 2. Materials and Methods

### 2.1 Animals and husbandry

We collected six of the twelve spider species studied, *Amaurobius similis* (Lace-Weaver), *Eratigena atrica* (Giant House Spider), *Larinioides sclopetarius* (Bridge Spider), *Meta menardi* (European Cave Spider), *Pholcus phalangioides* (Cellar Spider) and *Steatoda nobilis* (Noble False Widow) between August and November of 2021 and 2022, in sites between Co. Galway and Co. Sligo in Ireland. The other six species: *Heteropoda venatoria* (Giant Crab Spider), *Monocentropus balfouri* (Socotra Island Blue Baboon Tarantula), *Phormictopus cancerides* (Hispaniola Giant Tarantula), *Piloctenus haematostoma* (Guinean Wandering Spider), *Cupiennius coccineus* and *Cupiennius salei* (Bromeliad spiders) were either donated by Mark Stockmann, a captive breeder and owner of Buthidae.eu based in Hörstel, Germany or purchased from “The Pet Factory”, also based in Germany. All spiders were kept at constant room temperature (21°C), with the exception of *M. balfouri* and *P. haematostoma* (27°C) and *M. menardi*, which was stored at 10°C with moderate humidity (60-70%) as they are prone to desiccation (Howarth 1983). All spiders were not fed for two weeks before venom extraction to allow for adequate venom production, as per Boevé et al. (1995).

House crickets (*Acheta domesticus)* were purchased from retailers in Co. Galway while Common rough woodlice (*Porcellio scaber*) were collected from stone walls in Co. Galway. Specimens were selected based on their weight to fall within the range of 0.1 g – 0.5 g (median weight of 0.3 g per cohort) for *A. domesticus* and within the range of 0.06 g – 0.12 g (median weight of 0.09 g per cohort) for *P. scaber*. Prior to the bioassays, crickets and woodlice were housed in separate, large plastic boxes with egg cardboard hides (crickets) or bark covered with moss (woodlice) and provided with moistened pieces of peeled organic carrot for a minimum of 24 hours at 21^0^C.

### 2.2 Spider Venom Extractions

Venom was extracted from all spiders through electrical stimulation as described in Lyons et al. (2023), except for *P. phalangioides*, for which venom was obtained by excising the venom glands (Lyons *et al*. 2023). For all species, the venoms of both males and females were pooled. Venom was extracted from each spider only once to minimise effects of multiple extractions on the venom samples except for both *Cupiennius* spp. and *H. venatoria*, for which a second extraction was performed one month after the original extraction. Venom samples were flash frozen in liquid nitrogen, freeze-dried in a lyophiliser and stored at -80^0^C until required for bioassays (S1, S5: Table S1).

### 2.3 Spider Venom Bioassays

To determine the median lethal dose (LD_50_) (Finney 1985) and median effective (paralysis) dose (ED_50_) (Pearce *et al*. 1995) for each species venom, *in-vivo* bioassays were conducted in cricket and woodlice cohorts of 20 individuals each (n = 20). First, lyophilised venom samples from each spider species were weighed and rehydrated with 0.01 M Phosphate-buffered saline (PBS) to the original concentration of the crude venom initially extracted. Venom concentrations for each species were then determined based on the amount of venom available, starting as high as possible while allowing for serial dilution of the venom to at least two lower concentrations. For each venom concentration and prey model cohort, five microlitres (μl) of venom per specimen was injected using a 50 µl microsyringe (Hamilton Neurosyringe, model 1705RN, point style 3). Crickets were injected intrathoracically between the middle and hind leg, two millimetres (mm) deep while woodlice were injected ventrally, two mm deep, just posterior of the genital papilla on the first pleopod. Once injected, crickets and woodlice were maintained individually in petri dishes with damp paper towel and were checked for signs of paralysis or death at set intervals (1, 3, 5, 7, 10, 15, 30, 60, 120, 240 and 1440 minutes). Paralysis was recorded when a specimen was unable to move normally after a light touch with forceps (Michálek *et al*. 2019) and could not right itself within one minute after being flipped upside down (Khamtorn *et al*. 2020). Cricket and woodlice mortality was recorded if no response to gentle stimulus was observed from the moment the test was initiated until the end of the experiment (24 hours). After the initial three concentrations were tested in both prey models, for each species venom, additional venom concentrations were tested until there was insufficient venom to produce another viable concentration or the previous venom concentration tested proved ineffective (S1, S5: Table S2). Two control cohorts (n = 20) for each prey model were also tested using 0.01 M PBS instead of venom, with all crickets and woodlice surviving 24 hours post-injection and no signs of paralysis noted throughout the experiments.

### 2.4 LD_50_ and ED_50_ Calculations

To calculate each LD_50_ and ED_50_ value in milligrams of venom per kilogram of prey mass (mg/kg), we used Probit models from the Mass package (Venables and Ripley 2013) in R version 4.5.0 (Team 2022) (S1, S3). The LD_50_ value for each species venom, per prey model, was calculated using the number of deaths that occurred within 24 hours, per venom concentration tested, as per Lyons et al., (2023). The ED_50_ value for each species venom, per prey model, was calculated using the maximum number of paralysed individuals reached for each venom concentration within four hours, as four hours is one of the later times reported for the effects of the paralysis to conclude and for lethal effects to begin (Herzig and Hodgson 2008). In addition, supplementary ED_50_ values were calculated using the maximum number of paralysed individuals reached for each venom concentration within one hour, the most frequently used timepoint in the literature (Quistad *et al*. 1992; Dutertre *et al*. 2014; Michálek *et al*. 2019; Michálek *et al*. 2022), to determine if the difference between one hour and four hours as endpoints notably changes the resulting ED_50_ values and the relationship between LD_50_ and ED_50_ (S1, S3, S5: Table S3).

### 2.5 Data and phylogeny for larger scale comparative analysis

We collated additional LD_50_ and ED_50_ values from the literature to conduct a larger-scale analysis that could be compared with the smaller-scale analysis involving the 12 species potency values produced in the lab. Only potency values measured in microliters of venom per gram of prey mass (μl/g) were included as this was the most prevalent unit of measurement for spider ED_50_ values in the literature. Furthermore, potency values were only included if an ED_50_ value had a corresponding LD_50_ value for the same species venom and both values were tested in the same prey model species (S2).

To build a phylogeny to account for non-independence of data due to common ancestry in our larger-scale analysis, we used Wolff *et al*. (2022) as a backbone phylogeny, following the approach used in Lyons *et al*. (2025). For species not present in Wolff’s phylogeny, we inferred their position based on phylogenetic evidence from the literature (Maddison and Hedin 2003; Edwards 2004; Bolzern *et al*. 2013; Azevedo *et al*. 2018; Ortiz *et al*. 2018; Piacentini and Ramírez 2019; Guerrero-Fuentes *et al*. 2024) (S4). All steps performed to produce the final phylogeny were completed in R version 4.5.0 (Team 2022) (S2, S4, S5: Figure S3).

### 2.6 Statistical Analyses

To determine the relationship between LD_50_ and ED_50_ in the 12 spider species venoms tested in the lab, we fitted Generalised linear models (GLM) (Dey *et al*. 2000) using the ‘stats’ statistical package in R version 4.5.0 (Team 2022) (S1, S3). We included the prey model used to measure potency (*A. domesticus, P. scaber*) as a categorical factor. We also ran two supplementary GLMs. The first supplementary GLM tested the relationship between LD_50_ and ED_50_ using ED_50_ values calculated with the one hour endpoint while the second supplementary model tested the relationship between LD_50_ and ED_50_ for eleven species, with *P. phalangioides* excluded as its venom was extracted using a different method, which can potentially affect a spider’s ED_50_ value (Lyons *et al*. 2023) (S1, S3, S5).

For the larger-scale analysis testing the relationship between 40 spider species LD_50_ and ED_50_ values collated from the literature, we fitted a Bayesian phylogenetic mixed model, using the Markov chain Monte Carlo generalized linear mixed model (MCMCglmm) package (Hadfield 2010) in R version 4.5.0 (Team 2022), which allows for the inclusion of variance terms to account for multiple observations per species and the inclusion of a phylogenetic term (Hadfield 2010). We controlled for pseudoreplication due to shared ancestry between species through the ‘animal’ term, which uses a distance matrix of the phylogenetic distance between species to control for the expected similarity in factor values. We then calculated the relative variance attributable to the animal term as h^2^, which can be interpreted similarly to the phylogenetic lambda value (Hadfield and Nakagawa 2010). To include multiple LD_50_ and ED_50_ measures for each species in our analysis, we used a random term for species, similar to previous comparative models of venom variation (Healy *et al*. 2019; Lyons *et al*. 2020; Lyons *et al*. 2025). As the MCMCglmm is a Bayesian approach that requires specifying priors, we fit all models using standard, flat, non-informative priors, which assume no prior expectations for the estimated values across the model. We used a burn-in of 40 000 and a thinning of 100 over 2 400 000 iterations to ensure that the sample sizes exceeded 1000 for all parameter estimates. We tested for convergence using the Gelman–Rubin statistic over three separate chains (Hadfield 2010). Significance of an estimate is determined when the 95% credibility interval (CI) does not cross zero (Hadfield 2010). In all of our models, we included log_10_ of ED_50_ as the response variable and log_10_ of LD_50_ as the independent variable (S5).

## 3.0 Results

### 3.1 Relationship between LD_50_ and ED_50_ in our experimentally measured spider venoms (12 species analysis)

We measured the venoms of 12 spider species from nine different families (Amaurobiidae, Trechaleidae, Agelenidae, Sparassidae, Araneidae, Tetragnathidae, Theraphosidae, Pholcidae and Ctenidae) with a total of 48 venom potency measures (24 LD_50_ mg/kg (24 hr) and 24 ED_50_ mg/kg (4 hr)) performed on two prey models, crickets (*A. domesticus*) and woodlice (*P. scaber*) (Table 1, S1). LD_50_ values in crickets ranged from 0.72 mg/kg for *S. nobilis* venom to 58.85 mg/kg for *P. cancerides* venom while LD_50_ values in woodlice ranged from 7.47 mg/kg for *S. nobilis* venom to 300.54 mg/kg for *E. atrica* venom. ED_50_ (4 hr) values in crickets ranged from 0.036 mg/kg for *S. nobilis* venom to 16.7 mg/kg for *P. cancerides* venom while ED_50_ (4 hr) values in woodlice ranged from 0.87 for *S. nobilis* venom to 28.06 mg/kg for *P. cancerides* venom (Table 1, S1).

**Table 1:**
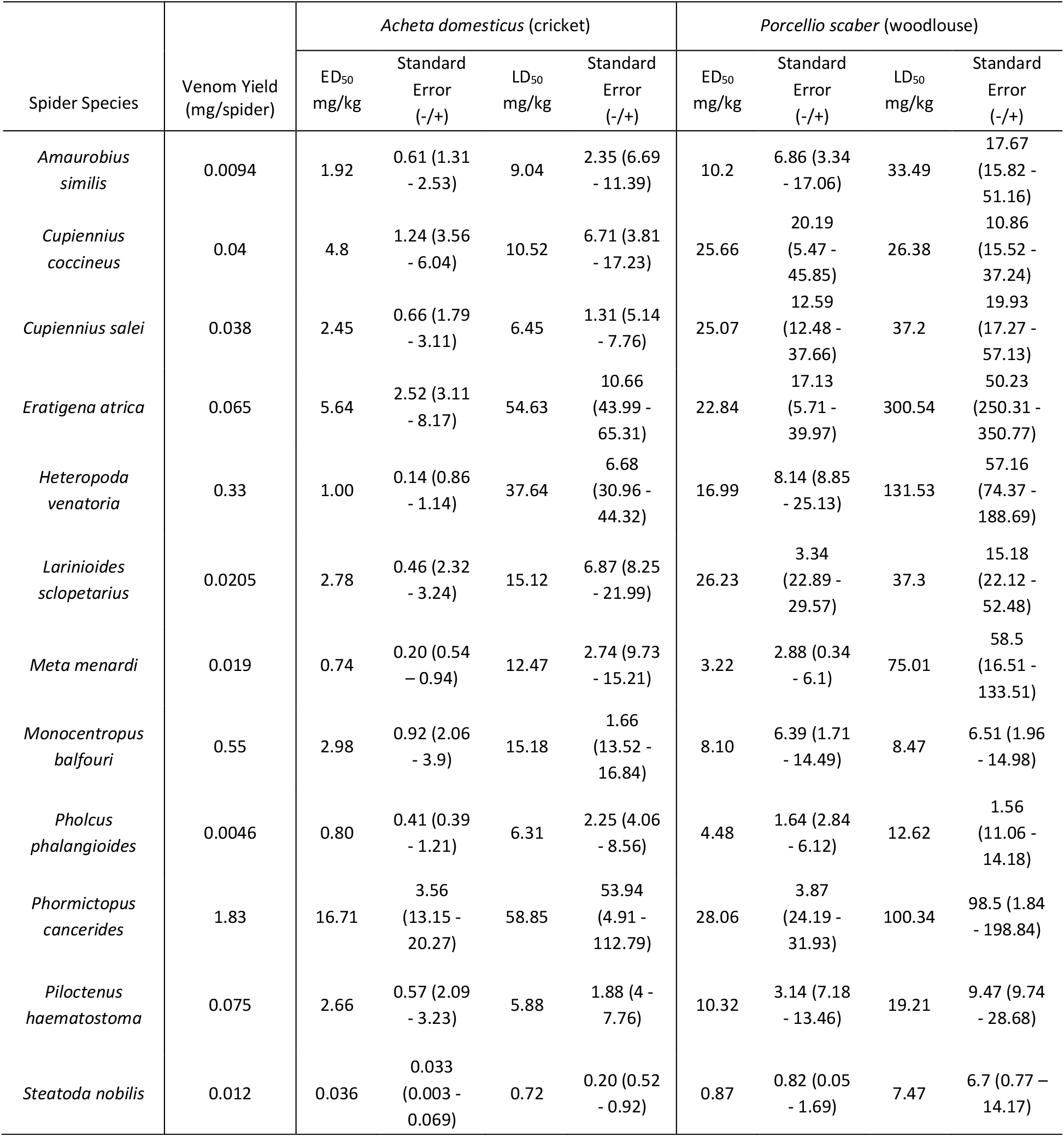
Calculated potency measures for 12 spider species venoms. tested on house cricket *A. domesticus* and woodlouse *P. scaber*. Presented here are the ED_50_ (mg/kg) (4 hr endpoint) and LD_50_ (mg/kg) (24 hr endpoint) values for each species venom, in both prey models. Standard errors were calculated using the lyophilised (dried) venom protein mass (mg) for each species venom sample and the median prey model mass for crickets (0.0003 kg) or woodlice (0.00009 kg). *A. domesticus* data for *Eratigena atrica* venom was produced as part of Lyons et al. (2023). Each species venom yield, expressed in milligram of dried venom extracted per specimen (mg/spider), is also presented.

| Spider Species | Venom Yield (mg/spider) | *Acheta domesticus* (cricket) |  |  |  | *Porcellio scaber* (woodlouse) |  |  |  |
| --- | --- | --- | --- | --- | --- | --- | --- | --- | --- |
|  |  | ED<sub>50</sub> mg/kg | Standard Error (-/+) | LD<sub>50</sub> mg/kg | Standard Error (-/+) | ED<sub>50</sub> mg/kg | Standard Error (-/+) | LD<sub>50</sub> mg/kg | Standard Error (-/+) |
| *Amaurobius similis* | 0.0094 | 1.92 | 0.61 (1.31 - 2.53) | 9.04 | 2.35 (6.69 - 11.39) | 10.2 | 6.86 (3.34 - 17.06) | 33.49 | 17.67 (15.82 - 51.16) |
| *Cupiennius coccineus* | 0.04 | 4.8 | 1.24 (3.56 - 6.04) | 10.52 | 6.71 (3.81 - 17.23) | 25.66 | 20.19 (5.47 - 45.85) | 26.38 | 10.86 (15.52 - 37.24) |
| *Cupiennius salei* | 0.038 | 2.45 | 0.66 (1.79 - 3.11) | 6.45 | 1.31 (5.14 - 7.76) | 25.07 | 12.59 (12.48 - 37.66) | 37.2 | 19.93 (17.27 - 57.13) |
| *Eratigena atrica* | 0.065 | 5.64 | 2.52 (3.11 - 8.17) | 54.63 | 10.66 (43.99 - 65.31) | 22.84 | 17.13 (5.71 - 39.97) | 300.54 | 50.23 (250.31 - 350.77) |
| *Heteropoda venatoria* | 0.33 | 1.00 | 0.14 (0.86 - 1.14) | 37.64 | 6.68 (30.96 - 44.32) | 16.99 | 8.14 (8.85 - 25.13) | 131.53 | 57.16 (74.37 - 188.69) |
| *Larinioides sclopetarius* | 0.0205 | 2.78 | 0.46 (2.32 - 3.24) | 15.12 | 6.87 (8.25 - 21.99) | 26.23 | 3.34 (22.89 - 29.57) | 37.3 | 15.18 (22.12 - 52.48) |
| *Meta menardi* | 0.019 | 0.74 | 0.20 (0.54 - 0.94) | 12.47 | 2.74 (9.73 - 15.21) | 3.22 | 2.88 (0.34 - 6.1) | 75.01 | 58.5 (16.51 - 133.51) |
| *Monocentropus balfouri* | 0.55 | 2.98 | 0.92 (2.06 - 3.9) | 15.18 | 1.66 (13.52 - 16.84) | 8.10 | 6.39 (1.71 - 14.49) | 8.47 | 6.51 (1.96 - 14.98) |
| *Pholcus phalangoides* | 0.0046 | 0.80 | 0.41 (0.39 - 1.21) | 6.31 | 2.25 (4.06 - 8.56) | 4.48 | 1.64 (2.84 - 6.12) | 12.62 | 1.56 (11.06 - 14.18) |
| *Phormictopus cancerides* | 1.83 | 16.71 | 3.56 (13.15 - 20.27) | 58.85 | 53.94 (4.91 - 112.79) | 28.06 | 3.87 (24.19 - 31.93) | 100.34 | 98.5 (1.84 - 198.84) |
| *Piloctenus haematostoma* | 0.075 | 2.66 | 0.57 (2.09 - 3.23) | 5.88 | 1.88 (4 - 7.76) | 10.32 | 3.14 (7.18 - 13.46) | 19.21 | 9.47 (9.74 - 28.68) |
| *Steatoda nobilis* | 0.012 | 0.036 | 0.033 (0.003 - 0.069) | 0.72 | 0.20 (0.52 - 0.92) | 0.87 | 0.82 (0.05 - 1.69) | 7.47 | 6.7 (0.77 - 14.17) |

We found support for an isometric relationship between LD_50_ mg/kg and ED_50_ mg/kg (4 hr) measures of spider venom potency (β = 1.01, SE = 0.24, p value < 0.05, n = 48 venom potency measures for 12 species; Figure 1, S5: Table M1). Hence, when LD_50_ increases, ED_50_ increases approximately by the same amount. While the intercepts differed between prey models in the main model, these differences were not significant (β = -0.48, SE = 0.35, p value = 0.18; Figure 1, S5: Table M1). The relationship between LD_50_ mg/kg and ED_50_ mg/kg was also isometric when ED_50_ mg/kg was calculated using the one-hour endpoint (β = 0.94, SE = 0.23, p value = 0.001, n = 46 venom potency measures for 11 species; S5: Figure S1, Table S3), while the supplementary model excluding *P. phalangioides* was also found to support an isometric relationship between LD_50_ mg/kg and ED_50_ mg/kg (4 hr) measures of spider venom potency (β = 1.00, SE = 0.25, p value = 0.001, n = 46 venom potency measures for 11 species; S5: Figure S2 Table S4).

**Figure 1:**
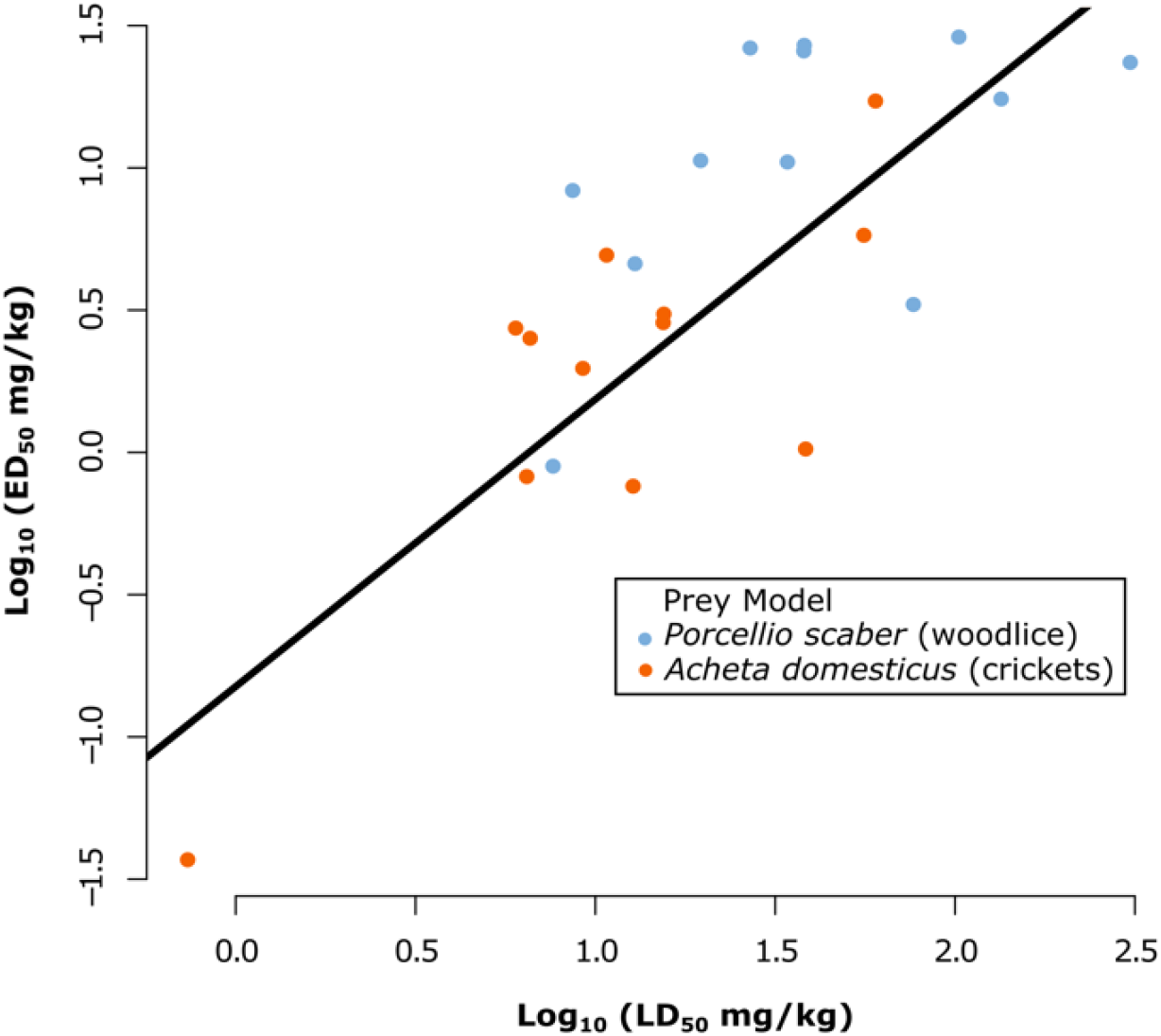
Relationship between log_10_ of LD_50_ (mg/kg) and log_10_ of ED_50_ (mg/kg) for 24 measures of LD_50_ and 24 measures of ED_50_ (4 hr endpoint) across 12 species venoms, tested in two prey models, *A. domesticus* (orange) and *P. scaber* (blue). The fitted line highlights the significant positive, isometric relationship between log_10_ LD_50_ (mg/kg) and log_10_ ED_50_ (mg/kg) (4 hr) (β = 1.01, SE = 0.24, p value = 0.0004; S5: Table M1).

### 3.2 Relationship between LD_50_ and ED_50_ in spider venoms (40 species analysis)

The comparative analysis based on data from the literature consisted of 40 spider species spanning 22 different families, with a total of 110 venom potency measures (55 LD_50_ μl/g and 55 ED_50_ μl/g (1 hr)) (S2). LD_50_ values ranged from 0.003 μl/g for *Haplodrassus sp*. venom tested on Tobacco hornworm *Manduca sexta* to 8.52 μl/g for *Stegodyphus lineatus* venom tested on *Pardosa* sp. spiders. ED_50_ values ranged from 0.00017 μl/g for *Latrodectus hesperus* venom tested on Oriental cockroach *Blatta orientalis* to 10.4 μl/g for *Scytodes sp*. venom tested on *M. sexta* (S2).

We also found support for a near isometric relationship between LD_50_ μl/g and ED_50_ μl/g measures of spider venom potency (S2) (β (slope) = 1.1, lower 95% CI = 0.81, upper 95% CI = 1.39; Figure 2, S5: Table M2). The model had an intercept of -0.52, which indicates that the venom volumes required to incapacitate prey are roughly between half to almost an order of magnitude lower than the volumes required for lethal effects. We find that the h^2^ value is 0.003, which indicates that there is a low phylogenetic signal.

**Figure 2:**
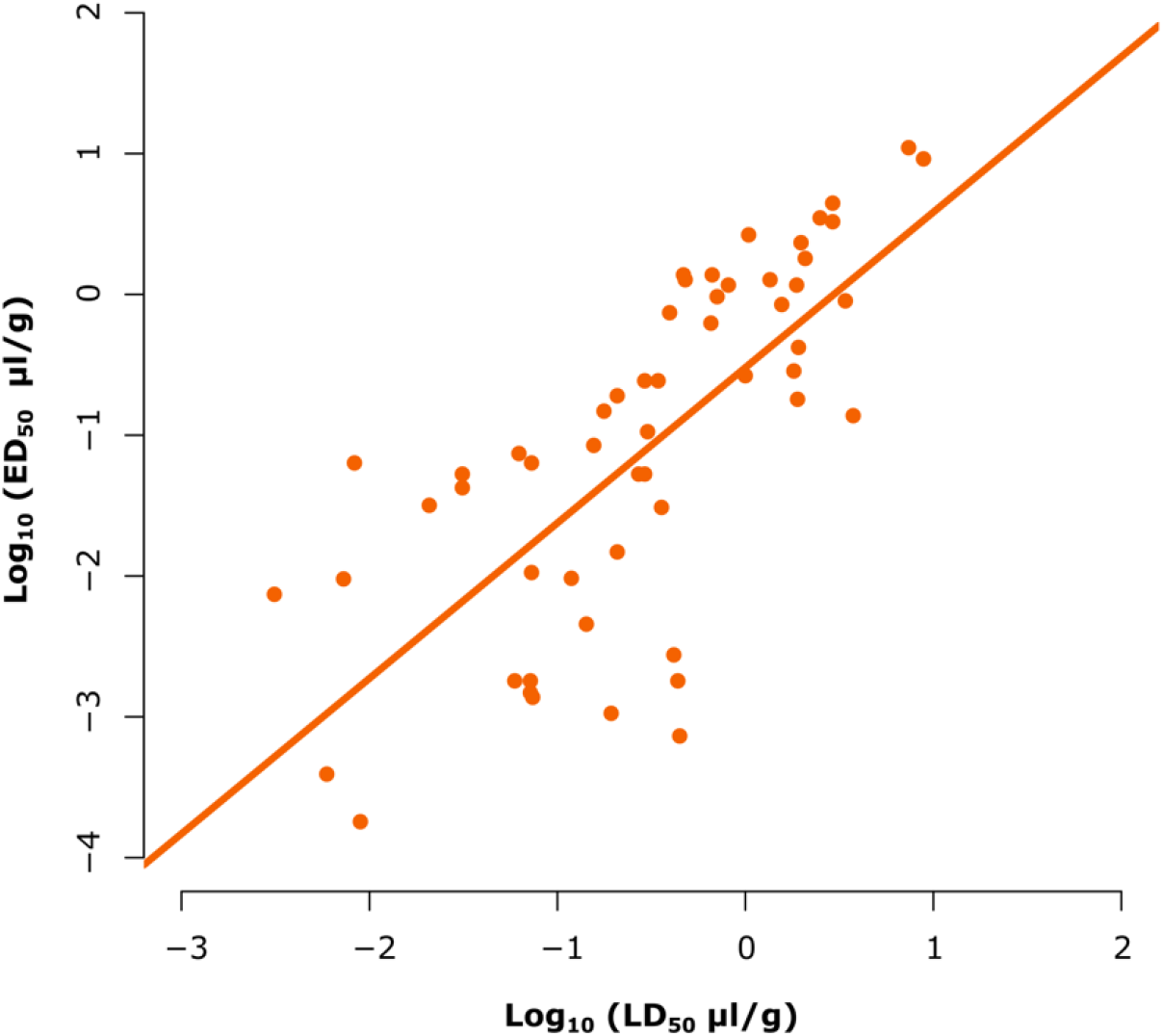
Relationship between log_10_ of LD_50_ (μl/g) and log_10_ of ED_50_ (μl/g) (1 hr) for 55 measures of LD_50_ and 55 measures of ED_50_ across 40 species venoms spanning 22 different families, performed on eight different prey models, including seven insect models and one arachnid model (S2). The fitted line highlights the significant positive, isometric relationship between log_10_ LD_50_ (μl/g) and log_10_ ED_50_ (μl/g) (β (slope) = 1.1, lower 95% CI = 0.81, upper 95% CI = 1.39; Figure 2, S5: Table M2).

## 4.0 Discussion

The results of both our experimental and comparative analyses demonstrate that spider venom potency measures, ED_50_ and LD_50_, have a near isometric, positive relationship. This suggests that spider venom ED_50_ and LD_50_ values are generally close proxies of one another, with an increase in paralysis efficiency resulting in a similar increase in lethality. Furthermore, we did not find support for the sublinear scaling that would indicate lethality is decoupled from the paralysing ability of venoms. Instead, our results indicate that either selection for lethality is comparable to the level of selection for paralysis efficiency, or that this correlation is a consequence of functionally coupled toxins, with selection for toxin paralysis efficiency positively impacting the lethality of the same toxins due to the overlapping nature of these functions.

While our analyses tested whole lyophilised spider venoms, with the modus operandi of each species venom requiring further investigation, it is likely that only specific toxins are functionally coupled between paralysis capacity and lethality. For example, several toxins are known to induce a gradual secondary paralysis after the initial rapid actions of other toxins, which can effectively lead to lethality. For example, the venoms of Australian funnel-web spiders (Atracidae) contain certain peptidic blockers of insect voltage-gated calcium channels that can induce a gradual, flaccid paralysis within 20 - 30 minutes (Chong *et al*. 2007; King and Hardy 2013). While such slow acting toxins may have been selected for their ability to prolong incapacitating effects in prey, they may also be tightly linked to later lethal effects, as their effects have been shown to be irreversible and typically target vital functions, which can ultimately lead to organ failure (King and Hardy 2013; Langenegger *et al*. 2019; Bolon *et al*. 2023).

Alternatively, the isometric relationship found here could indicate strong selection for slower acting toxins with lethal effects. For example, prolonged paralytic effects that result in prey mortality may be selected for in species that engage in food storage behaviours (Griffiths *et al*. 2003; Dippenaar-Schoeman and Harris 2005; Petráková *et al*. 2015), bite and run tactics (Pekár 2004; Pekár *et al*. 2014) or extensive feeding sessions on relatively large prey, or vertebrate prey, which can last from several hours to several days (Nyffeler and Gibbons 2022). However, the modus operandi for many spider venom toxins is still relatively unexplored, with such slower acting toxins only being observed in a few species (Adams 2004; Herzig and Hodgson 2008; Michálek *et al*. 2019). Hence, whether the isometric relationship found here is due to strong mechanistic coupling or direct selection is unclear and will require further data on the specific actions of toxins across a broader range of species.

Whether it is due to strong coupling or selection for prolonged effects in venoms, the isometric relationship between LD_50_ and ED_50_ observed here indicates that both measures are good proxies for one another, at least in spiders. While ED_50_ is regarded as the more ecologically relevant measure, available datasets remain sparse in the literature, for all venomous taxa. However, LD_50_ measures for spider venoms are far more prevalent (Pekár *et al*. 2021) and these could be used as tentative proxies to address both species-specific and large-scale questions related to venom potency. Our analyses support the results of previous studies that used LD_50_ values to test hypotheses related to spider venom potency (Wigger *et al*. 2002; Herzig *et al*. 2004; Eggs *et al*. 2015; Clémençon *et al*. 2020) and the idea that both LD_50_ and ED_50_ values can be used to identify ecological patterns in spider venoms, such as prey-specific potency (Michálek *et al*. 2019; Valenzuela-Rojas *et al*. 2019; Michálek *et al*. 2022; Lyons *et al*. 2025). However, whether our findings extend to other venomous groups is unclear. Specifically, future studies focusing on venomous groups that are unlikely to benefit from long-term or lethal effects are expected to offer further insight into whether lethality is generally functional coupled with paralysis efficiency or is more directly selected for in predation-oriented venoms.

Overall, our results indicate that the spider venom potency measures ED_50_ and LD_50_, as well as the functions they measure, have a positive, isometric relationship, meaning that selection for increased paralysis efficiency results in a similar increase for venom lethality. This relationship suggests: 1) that spider venom toxins responsible for incapacitating prey (especially prolonged paralytic effects) may be functionally coupled with lethality and 2) that ED_50_ and LD_50_ are good proxies for one another, meaning historically available LD_50_ values may be tentatively used to compare general spider venom potencies, provided they are based on the same prey model.

## CRediT authorship contribution statement

**Keith Lyons:** Conceptualization, Methodology, Software, Validation, Spider collection, Venom extraction and processing, Formal analysis, Investigation, Data curation, Writing – original draft, preparation, Visualization, Funding acquisition. **Dayle Leonard:** Methodology, Spider collection, Venom Spider collection, Venom extraction and processing, Formal analysis, Investigation, Writing – draft review. **Leona McSharry, Michael Martindale, Brandon L. Collier, Aiste Vitkauskaite, John P. Dunbar:** Spider collection, Venom extraction and processing, Formal analysis, Writing – draft review. **Michel M. Dugon:** Conceptualization, Methodology, Spider collection, Venom Spider collection, Venom extraction and processing, Formal analysis, Investigation, Resources, Writing – review & editing, Supervision, Funding acquisition. **Kevin Healy:** Conceptualization, Methodology, Software, Formal analysis, Investigation, Writing – review & editing, Supervision, Funding acquisition.

## Data availability

All data and code are included in the supplementary files S1 – S5 which can be obtained from the Dryad data repository link provided by bioRxiv post-submission or by contacting the corresponding author directly via email.

## Acknowledgments

We thank Dr Yuri Simone for putting us in contact with Mark Stockmann who kindly provided both *Cupiennius* spp. for our research. We thank Eoin MacLoughlin, Maeve Edwards, Dariusz Nowak and Aedin McAleer for their technical assistance. We thank Dr Sam Afoullous, Prof Olivier Thomas, Dr Clara Simon, Alisha Nelly (MSc), Dr Antoine Fort and Dr Ronan Sulpice for their assistance with the lyophilisation of the spider venom samples. K.L. would like to thank the following neighbours and family members for their assistance with spider sampling: Caitlin, Martin, Jack, Gerard, Aileen and Conor of the Warde family, Orla and Padraig Quirke, Bernie and Noreen Hansberry and Carrie and Oliver Craughwell.

