## Supplementary material for "Paralysis Efficiency (ED_50_) Scales Linearly with Lethality (LD_50_) in Spider Venoms": S3: S3_models_code_TX.Rmd

````
----------------
          Title: "S3_Relationship_between_ED50_and_LD50_2025_code"
         Author: "Keith Lyons"
Purpose of file: "to demonstrate how ED50 and LD50 was calculated for each spider and the relationship between them"
           Date: "27-08-2025"
         Output: html_document
----------------


```{r load packages}

library(ggplot2)
library(MASS)
library(MCMCglmm)
library(lme4)
library(invgamma)
library(mulTree)


### Installing the mulTree package if it doesn't work ###

# install.packages("devtools")
# library(devtools)
# install_github("TGuillerme/mulTree", ref = "release")
# library(mulTree)
```

Before running the code below, go to Session tab above -> Set Working Directory -> Choose Directory,
                                                                                   select location of data files

```{r load dataset}

bioassay_data_raw <- read.csv("S1_2025.csv",
                              sep = ",",
                              header = T)

```


##############################################################################################################################
############# Calculating the ED50 and LD50 Dose (%) per spider species AND prey species (crickets and woodlice) #############
##############################################################################################################################


```{r subsetting the ED50 data}


### The ED50 dataset contains the peak number of incapacitations per concentration, 
### for all concentrations tested,
### for each spider species venom and prey model tested

ED50_data <- bioassay_data_raw[,c("species",
                                  "taxa",
                                  "prey_species", 
                                  "time_minutes", 
                                  "venom_conc_percent" , 
                                  "venom_conc_mgul",
                                  "alive", 
                                  "incapacitated", 
                                  "dead", 
                                  "incapacitated_or_dead", 
                                  "paralysis_peak_time",
                                  "paralysis_peak_time_1hr",
                                  "ED50_dose_mgul", 
                                  "ED50_mgkg",
                                  "ED50_dose_mgul_1hr",
                                  "ED50_mgkg_1hr",
                                  "body_length_mean_mm", 
                                  "venom_yield_mg_spider", 
                                  "venom_yield_mgul_spider", 
                                  "comp_rep", 
                                  "yield_rep" 
                                  )]


### next exclude all non-peak ED50 data
### peak data refers to the timepoint when the highest or peak number of incapacitations was observed over 4 hours
### peak data 1hr refers to the timepoint when the highest or peak number of incapacitations was observed over 1 hr.

ED50_data_peaks <- droplevels(ED50_data[!ED50_data$paralysis_peak_time == 'non_peak',])

ED50_data_peaks_1hr <- droplevels(ED50_data[!ED50_data$paralysis_peak_time_1hr == 'non_peak',])
                                      

```


```{r subsetting the LD50 data}

### The LD50 dataset contains the number of deaths at endpoint 1440 minutes (24 hours) per concentration, 
### for all concentrations tested,
### for each spider species venom and prey model tested

LD50_data <- bioassay_data_raw[,c("species",
                                  "taxa",
                                  "prey_species", 
                                  "time_minutes", 
                                  "venom_conc_percent" , 
                                  "venom_conc_mgul",
                                  "alive", 
                                  "incapacitated", 
                                  "dead", 
                                  "incapacitated_or_dead", 
                                  "LD50_dose_mgul", 
                                  "LD50_mgkg", 
                                  "body_length_mean_mm", 
                                  "venom_yield_mg_spider", 
                                  "venom_yield_mgul_spider", 
                                  "comp_rep", 
                                  "yield_rep"
                                  )]


### next exclude all timepoints excluding endpoint 1440 minutes (24 hours) data

LD50_data24 <- droplevels(LD50_data[!LD50_data$time_minutes != '1440',])


```


Each of the following blocks of code contains the code necessary for 
calculating each spider species ED50 (4 hr), LD50 (24 hr) and ED50 (1hr)/

```{r Amaurobius similis ED50 and LD50}

#############################
### First, calculate ED50 ###
#############################

### this subsets dataset

A_similis_ED50_crickets <- ED50_data_peaks[ED50_data_peaks$species == "Amaurobius_similis" & ED50_data_peaks$prey_species == "Acheta_domesticus", ]                                                                      

model_A_similis_ED50_crickets <- glm(cbind(A_similis_ED50_crickets$incapacitated_or_dead, 
                                           A_similis_ED50_crickets$alive) ~ (A_similis_ED50_crickets$venom_conc_percent), 
                                           binomial(link = probit))

summary(model_A_similis_ED50_crickets)


### plotting the data

### this just generates a sequence of numbers
ypredicts <- seq(0, 1, 0.01)
xpredicts <- dose.p(model_A_similis_ED50_crickets,p=c(ypredicts))


plot(A_similis_ED50_crickets$incapacitated_or_dead/20 ~ (A_similis_ED50_crickets$venom_conc_percent),
     ylab = "Proportion_incap",
     xlab = "Dosage",
     pch = 16)

lines(unlist(xpredicts), ypredicts)


A_similis_ED50_crickets_result <- dose.p(model_A_similis_ED50_crickets, p=c(0.5))
A_similis_ED50_crickets_result
#                   Dose       SE
#    p = 0.5: 0.7281596 0.2259169


### Converting % dose to mg/kg ###

# The calculated dose is the % concentration dose (mg/ul) required to achieve an ED50 in crickets of median mass 0.0003 kg.
# To get ED50 mg/kg, you first figure out how much 100% of the sample equals (0.079080 mg/ul).
# 100% then divided by 100 to get 1% = 0.000791 mg/ul.
# 1% multiplied by % dose = 0.73 = 0.000577 mg/ul.
# 0.000577 mg/ul is then divided by cricket median mass to get mg/kg (0.000577/0.0003 = 1.92 mg/kg).
# Repeat with the standard error dose to get standard range of error in mg/kg (0.61 mg/kg).


### now same but for woodlice

### this subsets dataset (woodlice)

A_similis_ED50_woodlice <- ED50_data_peaks[ED50_data_peaks$species == "Amaurobius_similis" & ED50_data_peaks$prey_species == "Porcellio_scaber", ]                                                                      

model_A_similis_ED50_woodlice <- glm(cbind(A_similis_ED50_woodlice$incapacitated_or_dead, 
                                           A_similis_ED50_woodlice$alive) ~ (A_similis_ED50_woodlice$venom_conc_percent), 
                                           binomial(link = probit))                                           

summary(model_A_similis_ED50_woodlice)


### plotting the data

### this just generates a sequence of numbers
ypredicts <- seq(0, 1, 0.01)
xpredicts <- dose.p(model_A_similis_ED50_woodlice,p=c(ypredicts))


plot(A_similis_ED50_woodlice$incapacitated_or_dead/20 ~ (A_similis_ED50_woodlice$venom_conc_percent),
     ylab = "Proportion_incap",
     xlab = "Dosage",
     pch = 16)

lines(unlist(xpredicts), ypredicts)


A_similis_ED50_woodlice_result <- dose.p(model_A_similis_ED50_woodlice, p=c(0.5))
A_similis_ED50_woodlice_result
#                  Dose        SE
#     p = 0.5: 1.164152 0.7762379


### Converting % dose to mg/kg ###

# The calculated dose is the % concentration dose (mg/ul) required to achieve an ED50 in woodlice of median mass 0.00009 kg.
# To get ED50 mg/kg, you first figure out how much 100% of the sample equals (0.079080 mg/ul).
# 100% then divided by 100 to get 1% = 0.000791 mg/ul.
# 1% multiplied by % dose = 1.16 = 0.000918 mg/ul.
# 0.000918 mg/ul is then divided by woodlice median mass to get mg/kg (0.000918/0.00009 = 10.2 mg/kg).
# Repeat with the standard error dose to get standard range of error in mg/kg (6.86 mg/kg).


##########################
### Now calculate LD50 ###
##########################


### this subsets dataset

A_similis_LD50_crickets <- LD50_data24[LD50_data24$species == "Amaurobius_similis" & LD50_data24$prey_species == "Acheta_domesticus", ]

model_A_similis_LD50_crickets <- glm(cbind(A_similis_LD50_crickets$dead, 
                                           A_similis_LD50_crickets$alive) ~ (A_similis_LD50_crickets$venom_conc_percent), 
                                           binomial(link = probit))
                                       

summary(model_A_similis_LD50_crickets)


### plotting the data

### this just generates a sequence of numbers
ypredicts <- seq(0, 1, 0.01)
xpredicts <- dose.p(model_A_similis_LD50_crickets,p=c(ypredicts))


plot(A_similis_LD50_crickets$dead/20 ~ (A_similis_LD50_crickets$venom_conc_percent),
     ylab = "Proportion_dead",
     xlab = "Dosage",
     pch = 16)

lines(unlist(xpredicts), ypredicts)

### LD50 ###
A_similis_LD50_crickets_result <- dose.p(model_A_similis_LD50_crickets, p=c(0.50))
A_similis_LD50_crickets_result
#          Dose     SE
# p = 0.5: 3.427666 0.8892782


### Converting % dose to mg/kg ###

# The calculated dose is the % concentration dose (mg/ul) required to achieve an LD50 in crickets of median mass 0.0003 kg.
# To get LD50 mg/kg, you first figure out how much 100% of the sample equals (0.079080 mg/ul).
# 100% then divided by 100 to get 1% = 0.000791 mg/ul.
# 1% multiplied by % dose = 3.43 = 0.002713 mg/ul.
# 0.002713 mg/ul is then divided by cricket median mass to get mg/kg (0.002713/0.0003 = 9.04 mg/kg).
# Repeat with the standard error dose to get standard range of error in mg/kg (2.35 mg/kg).


### this subsets dataset (woodlice)

A_similis_LD50_woodlice <- LD50_data24[LD50_data24$species == "Amaurobius_similis" & LD50_data24$prey_species == "Porcellio_scaber", ]


model_A_similis_LD50_woodlice <- glm(cbind(A_similis_LD50_woodlice$dead, 
                                           A_similis_LD50_woodlice$alive) ~ (A_similis_LD50_woodlice$venom_conc_percent),
                                           binomial(link = probit))
                                       
summary(model_A_similis_LD50_woodlice)


### plotting the data

### this just generates a sequence of numbers
ypredicts <- seq(0, 1, 0.01)
xpredicts <- dose.p(model_A_similis_LD50_woodlice,p=c(ypredicts))


plot(A_similis_LD50_woodlice$dead/20 ~ (A_similis_LD50_woodlice$venom_conc_percent),
     ylab = "Proportion_dead",
     xlab = "Dosage",
     pch = 16)

lines(unlist(xpredicts), ypredicts)

### LD50 ###
A_similis_LD50_woodlice_result <- dose.p(model_A_similis_LD50_woodlice, p=c(0.50))
A_similis_LD50_woodlice_result
#          Dose     SE
# p = 0.5: 3.805169 2.012056


### Converting % dose to mg/kg ###

# The calculated dose is the % concentration dose (mg/ul) required to achieve an LD50 in woodlice of median mass 0.00009 kg.
# To get LD50 mg/kg, you first figure out how much 100% of the sample equals (0.079080 mg/ul).
# 100% then divided by 100 to get 1% = 0.000791 mg/ul.
# 1% multiplied by % dose = 3.81 = 0.003014 mg/ul.
# 0.003014 mg/ul is then divided by woodlice median mass to get mg/kg (0.003014/0.00009 = 33.49 mg/kg).
# Repeat with the standard error dose to get standard range of error in mg/kg (17.67 mg/kg).


########################################
##### Let's try with 1hr peak data #####
########################################


#################################
### First, calculate ED50 1hr ###
#################################

### this subsets dataset

A_similis_ED50_crickets_1hr <- ED50_data_peaks_1hr[ED50_data_peaks_1hr$species == "Amaurobius_similis" & ED50_data_peaks_1hr$prey_species == "Acheta_domesticus", ]                                                                      

model_A_similis_ED50_crickets_1hr <- glm(cbind(A_similis_ED50_crickets_1hr$incapacitated_or_dead, 
                                           A_similis_ED50_crickets_1hr$alive) ~ (A_similis_ED50_crickets_1hr$venom_conc_percent), 
                                           binomial(link = probit))

summary(model_A_similis_ED50_crickets_1hr)


### plotting the data

### this just generates a sequence of numbers
ypredicts <- seq(0, 1, 0.01)
xpredicts <- dose.p(model_A_similis_ED50_crickets_1hr,p=c(ypredicts))


plot(A_similis_ED50_crickets_1hr$incapacitated_or_dead/20 ~ (A_similis_ED50_crickets_1hr$venom_conc_percent),
     ylab = "Proportion_incap",
     xlab = "Dosage",
     pch = 16)

lines(unlist(xpredicts), ypredicts)


A_similis_ED50_crickets_result_1hr <- dose.p(model_A_similis_ED50_crickets_1hr, p=c(0.5))
A_similis_ED50_crickets_result_1hr
#                   Dose       SE
#    p = 0.5: 0.7281596 0.2259169


### Converting % dose to mg/kg ###

# The calculated dose is the % concentration dose (mg/ul) required to achieve an ED50 in crickets of median mass 0.0003 kg.
# To get ED50 mg/kg, you first figure out how much 100% of the sample equals (0.079080 mg/ul).
# 100% then divided by 100 to get 1% = 0.000791 mg/ul.
# 1% multiplied by % dose = 0.73 = 0.000577 mg/ul.
# 0.000577 mg/ul is then divided by cricket median mass to get mg/kg (0.000577/0.0003 = 1.92 mg/kg).
# Repeat with the standard error dose to get standard range of error in mg/kg (0.61 mg/kg).


### now same but for woodlice

### this subsets dataset (woodlice)

A_similis_ED50_woodlice_1hr <- ED50_data_peaks_1hr[ED50_data_peaks_1hr$species == "Amaurobius_similis" & ED50_data_peaks_1hr$prey_species == "Porcellio_scaber", ]                                                                      

model_A_similis_ED50_woodlice_1hr <- glm(cbind(A_similis_ED50_woodlice_1hr$incapacitated_or_dead, 
                                           A_similis_ED50_woodlice_1hr$alive) ~ (A_similis_ED50_woodlice_1hr$venom_conc_percent), 
                                           binomial(link = probit))                                           

summary(model_A_similis_ED50_woodlice_1hr)


### plotting the data

### this just generates a sequence of numbers
ypredicts <- seq(0, 1, 0.01)
xpredicts <- dose.p(model_A_similis_ED50_woodlice_1hr,p=c(ypredicts))


plot(A_similis_ED50_woodlice_1hr$incapacitated_or_dead/20 ~ (A_similis_ED50_woodlice_1hr$venom_conc_percent),
     ylab = "Proportion_incap",
     xlab = "Dosage",
     pch = 16)

lines(unlist(xpredicts), ypredicts)


A_similis_ED50_woodlice_result_1hr <- dose.p(model_A_similis_ED50_woodlice_1hr, p=c(0.5))
A_similis_ED50_woodlice_result_1hr
#                  Dose        SE
#     p = 0.5: 1.654906 0.7950264


### Converting % dose to mg/kg ###

# The calculated dose is the % concentration dose (mg/ul) required to achieve an ED50 in woodlice of median mass 0.00009 kg.
# To get ED50 mg/kg, you first figure out how much 100% of the sample equals (0.079080 mg/ul).
# 100% then divided by 100 to get 1% = 0.000791 mg/ul.
# 1% multiplied by % dose = 1.65 = 0.00130515 mg/ul.
# 0.00130515 mg/ul is then divided by woodlice median mass to get mg/kg (0.000918/0.00009 = 14.5 mg/kg).
# Repeat with the standard error dose to get standard range of error in mg/kg (6.99 mg/kg).


```


```{r Cupiennius coccineus ED50 and LD50}

#############################
### First, calculate ED50 ###
#############################

### this subsets dataset

C_coccineus_ED50_crickets <- ED50_data_peaks[ED50_data_peaks$species == "Cupiennius_coccineus" & ED50_data_peaks$prey_species == "Acheta_domesticus", ] 

model_C_coccineus_ED50_crickets <- glm(cbind(C_coccineus_ED50_crickets$incapacitated_or_dead, 
                                             C_coccineus_ED50_crickets$alive) ~ (C_coccineus_ED50_crickets$venom_conc_percent), 
                                             binomial(link = probit))
                                         
summary(model_C_coccineus_ED50_crickets)


### plotting the data

### this just generates a sequence of numbers
ypredicts <- seq(0, 1, 0.01)
xpredicts <- dose.p(model_C_coccineus_ED50_crickets,p=c(ypredicts))


plot(C_coccineus_ED50_crickets$incapacitated_or_dead/20 ~ (C_coccineus_ED50_crickets$venom_conc_percent),
     ylab = "Proportion_incap",
     xlab = "Dosage",
     pch = 16)

lines(unlist(xpredicts), ypredicts)


C_coccineus_ED50_crickets_result <- dose.p(model_C_coccineus_ED50_crickets, p=c(0.5))
C_coccineus_ED50_crickets_result
#                 Dose        SE
#    p = 0.5: 2.430895 0.6310878


### Converting % dose to mg/kg ###

# The calculated dose is the % concentration dose (mg/ul) required to achieve an ED50 in crickets of median mass 0.0003 kg.
# To get ED50 mg/kg, you first figure out how much 100% of the sample equals (0.0591966173361522 mg/ul).
# 100% then divided by 100 to get 1% = 0.000592 mg/ul.
# 1% multiplied by % dose = 2.43 = 0.001439 mg/ul.
# 0.001439 mg/ul is then divided by cricket median mass to get mg/kg (0.001439/0.0003 = 4.8 mg/kg).
# Repeat with the standard error dose to get standard range of error in mg/kg (1.24 mg/kg).


### this subsets dataset (woodlice)

C_coccineus_ED50_woodlice <- ED50_data_peaks[ED50_data_peaks$species == "Cupiennius_coccineus" & ED50_data_peaks$prey_species == "Porcellio_scaber", ]


model_C_coccineus_ED50_woodlice <- glm(cbind(C_coccineus_ED50_woodlice$incapacitated_or_dead, 
                                             C_coccineus_ED50_woodlice$alive) ~ (C_coccineus_ED50_woodlice$venom_conc_percent),
                                             binomial(link = probit))
                                      
summary(model_C_coccineus_ED50_woodlice)


### plotting the data

### this just generates a sequence of numbers
ypredicts <- seq(0, 1, 0.01)
xpredicts <- dose.p(model_C_coccineus_ED50_woodlice,p=c(ypredicts))


plot(C_coccineus_ED50_woodlice$incapacitated_or_dead/20 ~ (C_coccineus_ED50_woodlice$venom_conc_percent),
     ylab = "Proportion_incap",
     xlab = "Dosage",
     pch = 16)

lines(unlist(xpredicts), ypredicts)


C_coccineus_ED50_woodlice_result <- dose.p(model_C_coccineus_ED50_woodlice, p=c(0.5))
C_coccineus_ED50_woodlice_result
#          Dose     SE
# p = 0.5: 3.895541 3.07113


### Converting % dose to mg/kg ###

# The calculated dose is the % concentration dose (mg/ul) required to achieve an ED50 in woodlice of median mass 0.00009 kg.
# To get ED50 mg/kg, you first figure out how much 100% of the sample equals (0.0591966173361522 mg/ul).
# 100% then divided by 100 to get 1% = 0.000592 mg/ul.
# 1% multiplied by % dose = 3.9 = 0.002309 mg/ul.
# 0.002309 mg/ul is then divided by woodlice median mass to get mg/kg (0.002309/0.00009 = 25.66 mg/kg).
# Repeat with the standard error dose to get standard range of error in mg/kg (20.19 mg/kg).


##########################
### Now calculate LD50 ###
##########################

### this subsets dataset

C_coccineus_LD50_crickets <- LD50_data24[LD50_data24$species == "Cupiennius_coccineus" & LD50_data24$prey_species == "Acheta_domesticus", ]

model_C_coccineus_LD50_crickets <- glm(cbind(C_coccineus_LD50_crickets$dead, 
                                             C_coccineus_LD50_crickets$alive) ~ (C_coccineus_LD50_crickets$venom_conc_percent), 
                                             binomial(link = probit))
                                         
summary(model_C_coccineus_LD50_crickets)


### plotting the data

### this just generates a sequence of numbers
ypredicts <- seq(0, 1, 0.01)
xpredicts <- dose.p(model_C_coccineus_LD50_crickets,p=c(ypredicts))


plot(C_coccineus_LD50_crickets$dead/20 ~ (C_coccineus_LD50_crickets$venom_conc_percent),
     ylab = "Proportion_dead",
     xlab = "Dosage",
     pch = 16)

lines(unlist(xpredicts), ypredicts)

### LD50 ###
C_coccineus_LD50_crickets_result <- dose.p(model_C_coccineus_LD50_crickets, p=c(0.5))
C_coccineus_LD50_crickets_result
#          Dose     SE
# p = 0.5: 5.329767 3.402913


### Converting % dose to mg/kg ###

# The calculated dose is the % concentration dose (mg/ul) required to achieve an LD50 in crickets of median mass 0.0003 kg.
# To get LD50 mg/kg, you first figure out how much 100% of the sample equals (0.0591966173361522 mg/ul).
# 100% then divided by 100 to get 1% = 0.000592 mg/ul.
# 1% multiplied by % dose = 5.33 = 0.003155 mg/ul.
# 0.003155 mg/ul is then divided by cricket median mass to get mg/kg (0.003155/0.0003 = 10.52 mg/kg).
# Repeat with the standard error dose to get standard range of error in mg/kg (6.71 mg/kg).


### this subsets dataset (woodlice)

C_coccineus_LD50_woodlice <- LD50_data24[LD50_data24$species == "Cupiennius_coccineus" & LD50_data24$prey_species == "Porcellio_scaber", ]


model_C_coccineus_LD50_woodlice <- glm(cbind(C_coccineus_LD50_woodlice$dead, 
                                             C_coccineus_LD50_woodlice$alive) ~ (C_coccineus_LD50_woodlice$venom_conc_percent),
                                             binomial(link = probit))
                                         
summary(model_C_coccineus_LD50_woodlice)


### plotting the data

### this just generates a sequence of numbers
ypredicts <- seq(0, 1, 0.01)
xpredicts <- dose.p(model_C_coccineus_LD50_woodlice,p=c(ypredicts))


plot(C_coccineus_LD50_woodlice$dead/20 ~ (C_coccineus_LD50_woodlice$venom_conc_percent),
     ylab = "Proportion_dead",
     xlab = "Dosage",
     pch = 16)

lines(unlist(xpredicts), ypredicts)

### LD50 ###
C_coccineus_LD50_woodlice_result <- dose.p(model_C_coccineus_LD50_woodlice, p=c(0.5))
C_coccineus_LD50_woodlice_result
#          Dose     SE
# p = 0.5: 4.012199 1.651826


### Converting % dose to mg/kg ###

# The calculated dose is the % concentration dose (mg/ul) required to achieve an LD50 in woodlice of median mass 0.00009 kg.
# To get LD50 mg/kg, you first figure out how much 100% of the sample equals (0.0591966173361522 mg/ul).
# 100% then divided by 100 to get 1% = 0.000592 mg/ul.
# 1% multiplied by % dose = 4.01 = 0.002374 mg/ul.
# 0.002374 mg/ul is then divided by woodlice median mass to get mg/kg (0.002374/0.00009 = 26.38 mg/kg).
# Repeat with the standard error dose to get standard range of error in mg/kg (10.86 mg/kg).


########################################
##### Let's try with 1hr peak data #####
########################################


#################################
### First, calculate ED50 1hr ###
#################################

### this subsets dataset

C_coccineus_ED50_crickets_1hr <- ED50_data_peaks_1hr[ED50_data_peaks_1hr$species == "Cupiennius_coccineus" & ED50_data_peaks_1hr$prey_species == "Acheta_domesticus", ] 

model_C_coccineus_ED50_crickets_1hr <- glm(cbind(C_coccineus_ED50_crickets_1hr$incapacitated_or_dead, 
                                             C_coccineus_ED50_crickets_1hr$alive) ~ (C_coccineus_ED50_crickets_1hr$venom_conc_percent), 
                                             binomial(link = probit))
                                         
summary(model_C_coccineus_ED50_crickets_1hr)


### plotting the data

### this just generates a sequence of numbers
ypredicts <- seq(0, 1, 0.01)
xpredicts <- dose.p(model_C_coccineus_ED50_crickets_1hr,p=c(ypredicts))


plot(C_coccineus_ED50_crickets_1hr$incapacitated_or_dead/20 ~ (C_coccineus_ED50_crickets_1hr$venom_conc_percent),
     ylab = "Proportion_incap",
     xlab = "Dosage",
     pch = 16)

lines(unlist(xpredicts), ypredicts)


C_coccineus_ED50_crickets_result_1hr <- dose.p(model_C_coccineus_ED50_crickets_1hr, p=c(0.5))
C_coccineus_ED50_crickets_result_1hr
#                 Dose        SE
#    p = 0.5: 2.430895 0.6310878


### Converting % dose to mg/kg ###

# The calculated dose is the % concentration dose (mg/ul) required to achieve an ED50 in crickets of median mass 0.0003 kg.
# To get ED50 mg/kg, you first figure out how much 100% of the sample equals (0.0591966173361522 mg/ul).
# 100% then divided by 100 to get 1% = 0.000592 mg/ul.
# 1% multiplied by % dose = 2.43 = 0.001439 mg/ul.
# 0.001439 mg/ul is then divided by cricket median mass to get mg/kg (0.001439/0.0003 = 4.8 mg/kg).
# Repeat with the standard error dose to get standard range of error in mg/kg (1.24 mg/kg).


### this subsets dataset (woodlice)

C_coccineus_ED50_woodlice_1hr <- ED50_data_peaks_1hr[ED50_data_peaks_1hr$species == "Cupiennius_coccineus" & ED50_data_peaks_1hr$prey_species == "Porcellio_scaber", ]


model_C_coccineus_ED50_woodlice_1hr <- glm(cbind(C_coccineus_ED50_woodlice_1hr$incapacitated_or_dead, 
                                             C_coccineus_ED50_woodlice_1hr$alive) ~ (C_coccineus_ED50_woodlice_1hr$venom_conc_percent),
                                             binomial(link = probit))
                                      
summary(model_C_coccineus_ED50_woodlice_1hr)


### plotting the data

### this just generates a sequence of numbers
ypredicts <- seq(0, 1, 0.01)
xpredicts <- dose.p(model_C_coccineus_ED50_woodlice_1hr,p=c(ypredicts))


plot(C_coccineus_ED50_woodlice_1hr$incapacitated_or_dead/20 ~ (C_coccineus_ED50_woodlice_1hr$venom_conc_percent),
     ylab = "Proportion_incap",
     xlab = "Dosage",
     pch = 16)

lines(unlist(xpredicts), ypredicts)


C_coccineus_ED50_woodlice_result_1hr <- dose.p(model_C_coccineus_ED50_woodlice_1hr, p=c(0.5))
C_coccineus_ED50_woodlice_result_1hr
#           Dose         SE
# p = 0.5: -3.258487e+15 2.602993e+30


### Converting % dose to mg/kg ###

# - dose result means this result is unusable


```


```{r Cupiennius salei ED50 and LD50}

#############################
### First, calculate ED50 ###
#############################


### this subsets dataset

C_salei_ED50_crickets <- ED50_data_peaks[ED50_data_peaks$species == "Cupiennius_salei" & ED50_data_peaks$prey_species == "Acheta_domesticus", ]

model_C_salei_ED50_crickets <- glm(cbind(C_salei_ED50_crickets$incapacitated_or_dead, 
                                         C_salei_ED50_crickets$alive) ~ (C_salei_ED50_crickets$venom_conc_percent), 
                                         binomial(link = probit))
                                     
summary(model_C_salei_ED50_crickets)


### plotting the data

### this just generates a sequence of numbers
ypredicts <- seq(0, 1, 0.01)
xpredicts <- dose.p(model_C_salei_ED50_crickets,p=c(ypredicts))


plot(C_salei_ED50_crickets$incapacitated_or_dead/20 ~ (C_salei_ED50_crickets$venom_conc_percent),
     ylab = "Proportion_incap",
     xlab = "Dosage",
     pch = 16)

lines(unlist(xpredicts), ypredicts)


C_salei_ED50_crickets_result <- dose.p(model_C_salei_ED50_crickets, p=c(0.5))
C_salei_ED50_crickets_result
#                 Dose       SE
#    p = 0.5: 1.557906 0.4155914


### Converting % dose to mg/kg ###

# The calculated dose is the % concentration dose (mg/ul) required to achieve an ED50 in crickets of median mass 0.0003 kg.
# To get ED50 mg/kg, you first figure out how much 100% of the sample equals (0.047205 mg/ul).
# 100% then divided by 100 to get 1% = 0.000472 mg/ul.
# 1% multiplied by % dose = 1.56 = 0.000736 mg/ul.
# 0.000736 mg/ul is then divided by cricket median mass to get mg/kg (0.000736/0.0003 = 2.45 mg/kg).
# Repeat with the standard error dose to get standard range of error in mg/kg (0.66 mg/kg).


### this subsets dataset (woodlice)

C_salei_ED50_woodlice <- ED50_data_peaks[ED50_data_peaks$species == "Cupiennius_salei" & ED50_data_peaks$prey_species == "Porcellio_scaber", ]


model_C_salei_ED50_woodlice <- glm(cbind(C_salei_ED50_woodlice$incapacitated_or_dead, 
                                         C_salei_ED50_woodlice$alive) ~ (C_salei_ED50_woodlice$venom_conc_percent),
                                         binomial(link = probit))
                                     
summary(model_C_salei_ED50_woodlice)


### plotting the data

### this just generates a sequence of numbers
ypredicts <- seq(0, 1, 0.01)
xpredicts <- dose.p(model_C_salei_ED50_woodlice,p=c(ypredicts))


plot(C_salei_ED50_woodlice$incapacitated_or_dead/20 ~ (C_salei_ED50_woodlice$venom_conc_percent),
     ylab = "Proportion_incap",
     xlab = "Dosage",
     pch = 16)

lines(unlist(xpredicts), ypredicts)


C_salei_ED50_woodlice_result <- dose.p(model_C_salei_ED50_woodlice, p=c(0.5))
C_salei_ED50_woodlice_result
#          Dose     SE
# p = 0.5: 4.782947 2.402801


### Converting % dose to mg/kg ###

# The calculated dose is the % concentration dose (mg/ul) required to achieve an ED50 in woodlice of median mass 0.00009 kg.
# To get ED50 mg/kg, you first figure out how much 100% of the sample equals (0.047205 mg/ul).
# 100% then divided by 100 to get 1% = 0.000472 mg/ul.
# 1% multiplied by % dose = 4.78 = 0.002256 mg/ul.
# 0.002256 mg/ul is then divided by woodlice median mass to get mg/kg (0.002256/0.00009 = 25.07 mg/kg).
# Repeat with the standard error dose to get standard range of error in mg/kg (12.59 mg/kg).


##########################
### Now calculate LD50 ###
##########################

### this subsets dataset

C_salei_LD50_crickets <- LD50_data24[LD50_data24$species == "Cupiennius_salei" & LD50_data24$prey_species == "Acheta_domesticus", ]

model_C_salei_LD50_crickets <- glm(cbind(C_salei_LD50_crickets$dead, 
                                         C_salei_LD50_crickets$alive) ~ (C_salei_LD50_crickets$venom_conc_percent), 
                                         binomial(link = probit))
                                     
summary(model_C_salei_LD50_crickets)


### plotting the data

### this just generates a sequence of numbers
ypredicts <- seq(0, 1, 0.01)
xpredicts <- dose.p(model_C_salei_LD50_crickets,p=c(ypredicts))


plot(C_salei_LD50_crickets$dead/20 ~ (C_salei_LD50_crickets$venom_conc_percent),
     ylab = "Proportion_dead",
     xlab = "Dosage",
     pch = 16)

lines(unlist(xpredicts), ypredicts)


### LD50 ###
C_salei_LD50_crickets_result <- dose.p(model_C_salei_LD50_crickets, p=c(0.5))
C_salei_LD50_crickets_result
#          Dose     SE
# p = 0.5: 4.097421 0.8256657


### Converting % dose to mg/kg ###

# The calculated dose is the % concentration dose (mg/ul) required to achieve an LD50 in crickets of median mass 0.0003 kg.
# To get LD50 mg/kg, you first figure out how much 100% of the sample equals (0.047205 mg/ul).
# 100% then divided by 100 to get 1% = 0.000472 mg/ul.
# 1% multiplied by % dose = 4.10 = 0.001935 mg/ul.
# 0.001935 mg/ul is then divided by cricket median mass to get mg/kg (0.001935/0.0003 = 6.45 mg/kg).
# Repeat with the standard error dose to get standard range of error in mg/kg (1.31 mg/kg).


### this subsets dataset (woodlice)

C_salei_LD50_woodlice <- LD50_data24[LD50_data24$species == "Cupiennius_salei" & LD50_data24$prey_species == "Porcellio_scaber", ]


model_C_salei_LD50_woodlice <- glm(cbind(C_salei_LD50_woodlice$dead, 
                                         C_salei_LD50_woodlice$alive) ~ (C_salei_LD50_woodlice$venom_conc_percent),
                                         binomial(link = probit))
                                     
summary(model_C_salei_LD50_woodlice)


### plotting the data

### this just generates a sequence of numbers
ypredicts <- seq(0, 1, 0.01)
xpredicts <- dose.p(model_C_salei_LD50_woodlice,p=c(ypredicts))


plot(C_salei_LD50_woodlice$dead/20 ~ (C_salei_LD50_woodlice$venom_conc_percent),
     ylab = "Proportion_dead",
     xlab = "Dosage",
     pch = 16)

lines(unlist(xpredicts), ypredicts)


### LD50 ###
C_salei_LD50_woodlice_result <- dose.p(model_C_salei_LD50_woodlice, p=c(0.5))
C_salei_LD50_woodlice_result
#          Dose     SE
# p = 0.5: 7.111184 3.8117


### Converting % dose to mg/kg ###

# The calculated dose is the % concentration dose (mg/ul) required to achieve an LD50 in woodlice of median mass 0.00009 kg.
# To get LD50 mg/kg, you first figure out how much 100% of the sample equals (0.047205 mg/ul).
# 100% then divided by 100 to get 1% = 0.000472 mg/ul.
# 1% multiplied by % dose = 7.11 = 0.003356 mg/ul.
# 0.003356 mg/ul is then divided by woodlice median mass to get mg/kg (0.003356/0.00009 = 37.29 mg/kg).
# Repeat with the standard error dose to get standard range of error in mg/kg (19.93 mg/kg).


########################################
##### Let's try with 1hr peak data #####
########################################


#################################
### First, calculate ED50 1hr ###
#################################


### this subsets dataset

C_salei_ED50_crickets_1hr <- ED50_data_peaks_1hr[ED50_data_peaks_1hr$species == "Cupiennius_salei" & ED50_data_peaks_1hr$prey_species == "Acheta_domesticus", ]

model_C_salei_ED50_crickets_1hr <- glm(cbind(C_salei_ED50_crickets_1hr$incapacitated_or_dead, 
                                         C_salei_ED50_crickets_1hr$alive) ~ (C_salei_ED50_crickets_1hr$venom_conc_percent), 
                                         binomial(link = probit))
                                     
summary(model_C_salei_ED50_crickets_1hr)


### plotting the data

### this just generates a sequence of numbers
ypredicts <- seq(0, 1, 0.01)
xpredicts <- dose.p(model_C_salei_ED50_crickets_1hr,p=c(ypredicts))


plot(C_salei_ED50_crickets_1hr$incapacitated_or_dead/20 ~ (C_salei_ED50_crickets_1hr$venom_conc_percent),
     ylab = "Proportion_incap",
     xlab = "Dosage",
     pch = 16)

lines(unlist(xpredicts), ypredicts)


C_salei_ED50_crickets_result_1hr <- dose.p(model_C_salei_ED50_crickets_1hr, p=c(0.5))
C_salei_ED50_crickets_result_1hr
#                 Dose       SE
#    p = 0.5: 1.557906 0.4155914


### Converting % dose to mg/kg ###

# The calculated dose is the % concentration dose (mg/ul) required to achieve an ED50 in crickets of median mass 0.0003 kg.
# To get ED50 mg/kg, you first figure out how much 100% of the sample equals (0.047205 mg/ul).
# 100% then divided by 100 to get 1% = 0.000472 mg/ul.
# 1% multiplied by % dose = 1.56 = 0.000736 mg/ul.
# 0.000736 mg/ul is then divided by cricket median mass to get mg/kg (0.000736/0.0003 = 2.45 mg/kg).
# Repeat with the standard error dose to get standard range of error in mg/kg (0.66 mg/kg).


### this subsets dataset (woodlice)

C_salei_ED50_woodlice_1hr <- ED50_data_peaks_1hr[ED50_data_peaks_1hr$species == "Cupiennius_salei" & ED50_data_peaks_1hr$prey_species == "Porcellio_scaber", ]


model_C_salei_ED50_woodlice_1hr <- glm(cbind(C_salei_ED50_woodlice_1hr$incapacitated_or_dead, 
                                         C_salei_ED50_woodlice_1hr$alive) ~ (C_salei_ED50_woodlice_1hr$venom_conc_percent),
                                         binomial(link = probit))
                                     
summary(model_C_salei_ED50_woodlice_1hr)


### plotting the data

### this just generates a sequence of numbers
ypredicts <- seq(0, 1, 0.01)
xpredicts <- dose.p(model_C_salei_ED50_woodlice_1hr,p=c(ypredicts))


plot(C_salei_ED50_woodlice_1hr$incapacitated_or_dead/20 ~ (C_salei_ED50_woodlice_1hr$venom_conc_percent),
     ylab = "Proportion_incap",
     xlab = "Dosage",
     pch = 16)

lines(unlist(xpredicts), ypredicts)


C_salei_ED50_woodlice_result_1hr <- dose.p(model_C_salei_ED50_woodlice_1hr, p=c(0.5))
C_salei_ED50_woodlice_result_1hr
#          Dose     SE
# p = 0.5: 4.321536 1.574921


### Converting % dose to mg/kg ###

# The calculated dose is the % concentration dose (mg/ul) required to achieve an ED50 in woodlice of median mass 0.00009 kg.
# To get ED50 mg/kg, you first figure out how much 100% of the sample equals (0.047205 mg/ul).
# 100% then divided by 100 to get 1% = 0.000472 mg/ul.
# 1% multiplied by % dose = 4.32 = 0.00204 mg/ul.
# 0.00204 mg/ul is then divided by woodlice median mass to get mg/kg (0.00204/0.00009 = 22.66 mg/kg).
# Repeat with the standard error dose to get standard range of error in mg/kg (8.26 mg/kg).

```


```{r Eratigena atrica ED50 and LD50}

#############################
### First, calculate ED50 ###
#############################

### this subsets the data

E_atrica_ED50_cricket <- ED50_data_peaks[ED50_data_peaks$species == "Eratigena_atrica_EL" & ED50_data_peaks$prey_species == "Acheta_domesticus",]

model_E_atrica_ED50_cricket <- glm(cbind(E_atrica_ED50_cricket$incapacitated_or_dead, 
                                         E_atrica_ED50_cricket$alive) ~ (E_atrica_ED50_cricket$venom_conc_percent), 
                                         binomial(link = probit))
                                     
summary(model_E_atrica_ED50_cricket)

### plotting the data

ypredicts <- seq(0, 1, 0.01)
xpredicts <- dose.p(model_E_atrica_ED50_cricket,p=c(ypredicts))

plot(ED50_data_peaks$incapacitated_or_dead/20 ~ (ED50_data_peaks$venom_conc_percent),
     ylab = "Proportion Incapacitated",
     xlab = "Percent Dosage",
     pch = 16)

lines(unlist(xpredicts), ypredicts)


E_atrica_ED50_crickets_result <- dose.p(model_E_atrica_ED50_cricket, p=c(0.5))
E_atrica_ED50_crickets_result
#                 Dose        SE
#    p = 0.5: 1.119667 0.5028537


### Converting % dose to mg/kg ###

# The calculated dose is the % concentration dose (mg/ul) required to achieve an ED50 in crickets of median mass 0.0003 kg.
# To get ED50 mg/kg, you first figure out how much 100% of the sample equals (0.151197 mg/ul).
# 100% then divided by 100 to get 1% = 0.001512 mg/ul.
# 1% multiplied by % dose = 1.12 = 0.001693 mg/ul.
# 0.001693 mg/ul is then divided by cricket median mass to get mg/kg (0.001693/0.0003 = 5.64 mg/kg).
# Repeat with the standard error dose to get standard range of error in mg/kg (2.52 mg/kg).


### this subsets dataset (woodlice)

E_atrica_EL_ED50_woodlice <- ED50_data_peaks[ED50_data_peaks$species == "Eratigena_atrica_EL" & ED50_data_peaks$prey_species == "Porcellio_scaber", ]


model_E_atrica_EL_ED50_woodlice <- glm(cbind(E_atrica_EL_ED50_woodlice$incapacitated_or_dead, 
                                             E_atrica_EL_ED50_woodlice$alive) ~ (E_atrica_EL_ED50_woodlice$venom_conc_percent),
                                             binomial(link = probit))
                                         
summary(model_E_atrica_EL_ED50_woodlice)


### plotting the data

### this just generates a sequence of numbers
ypredicts <- seq(0, 1, 0.01)
xpredicts <- dose.p(model_E_atrica_EL_ED50_woodlice,p=c(ypredicts))


plot(E_atrica_EL_ED50_woodlice$incapacitated_or_dead/20 ~ (E_atrica_EL_ED50_woodlice$venom_conc_percent),
     ylab = "Proportion_incap",
     xlab = "Dosage",
     pch = 16)

lines(unlist(xpredicts), ypredicts)


E_atrica_EL_ED50_woodlice_result <- dose.p(model_E_atrica_EL_ED50_woodlice, p=c(0.5))
E_atrica_EL_ED50_woodlice_result
#                    Dose       SE
#     p = 0.5:   1.363679  1.02039


### Converting % dose to mg/kg ###

# The calculated dose is the % concentration dose (mg/ul) required to achieve an ED50 in woodlice of median mass 0.00009 kg.
# To get ED50 mg/kg, you first figure out how much 100% of the sample equals (0.151197 mg/ul).
# 100% then divided by 100 to get 1% = 0.001512 mg/ul.
# 1% multiplied by % dose = 1.36 = 0.002056 mg/ul.
# 0.002056 mg/ul is then divided by woodlice median mass to get mg/kg (0.002056/0.00009 = 22.84 mg/kg).
# Repeat with the standard error dose to get standard range of error in mg/kg (17.13 mg/kg).


##########################
### Now calculate LD50 ###
##########################

### this subsets the data

E_atrica_LD50_cricket <- LD50_data24[LD50_data24$species == "Eratigena_atrica_EL" & LD50_data24$prey_species == "Acheta_domesticus",]

model_E_atrica_LD50_cricket <- glm(cbind(E_atrica_LD50_cricket$dead, 
                                         E_atrica_LD50_cricket$alive) ~ (E_atrica_LD50_cricket$venom_conc_percent), 
                                         binomial(link = probit))
                                     
summary(model_E_atrica_LD50_cricket)

### plotting the data

ypredicts <- seq(0, 1, 0.01)
xpredicts <- dose.p(model_E_atrica_LD50_cricket,p=c(ypredicts))

plot(LD50_data$dead/20 ~ (LD50_data$venom_conc_percent),
     ylab = "Proportion dead",
     xlab = "Percent Dosage",
     pch = 16)

lines(unlist(xpredicts), ypredicts)


### LD50 ###
E_atrica_LD50_crickets_result <- dose.p(model_E_atrica_LD50_cricket, p=c(0.5))
E_atrica_LD50_crickets_result
#          Dose     SE
# p = 0.5: 10.84371 2.115504


### Converting % dose to mg/kg ###

# The calculated dose is the % concentration dose (mg/ul) required to achieve an LD50 in crickets of median mass 0.0003 kg.
# To get LD50 mg/kg, you first figure out how much 100% of the sample equals, for El this was 0.1511967578 mg/ul.
# 100% then divided by 100 to get 1% = 0.001511967578 mg/ul.
# 1% multiplied by % dose = 10.84 = 0.01638972854552 mg/ul.
# 0.01638972854552 mg/ul is then divided by cricket median mass to get mg/kg (0.01638972854552/0.0003 = 54.63 mg/kg)
# Repeat with the standard error dose to get standard range of error in mg/kg (10.66 mg/kg).


### this subsets dataset (woodlice)

E_atrica_EL_LD50_woodlice <- LD50_data24[LD50_data24$species == "Eratigena_atrica_EL" & LD50_data24$prey_species == "Porcellio_scaber", ]


model_E_atrica_EL_LD50_woodlice <- glm(cbind(E_atrica_EL_LD50_woodlice$dead, 
                                             E_atrica_EL_LD50_woodlice$alive) ~ (E_atrica_EL_LD50_woodlice$venom_conc_percent),
                                             binomial(link = probit))
                                         
summary(model_E_atrica_EL_LD50_woodlice)


### plotting the data

### this just generates a sequence of numbers
ypredicts <- seq(0, 1, 0.01)
xpredicts <- dose.p(model_E_atrica_EL_LD50_woodlice,p=c(ypredicts))


plot(E_atrica_EL_LD50_woodlice$dead/20 ~ (E_atrica_EL_LD50_woodlice$venom_conc_percent),
     ylab = "Proportion_dead",
     xlab = "Dosage",
     pch = 16)

lines(unlist(xpredicts), ypredicts)

### LD50 ###
E_atrica_EL_LD50_woodlice_result <- dose.p(model_E_atrica_EL_LD50_woodlice, p=c(0.5))
E_atrica_EL_LD50_woodlice_result
#          Dose     SE
# p = 0.5: 17.88788 2.991044


### Converting % dose to mg/kg ###

# The calculated dose is the % concentration dose (mg/ul) required to achieve an LD50 in woodlice of median mass 0.00009 kg.
# To get LD50 mg/kg, you first figure out how much 100% of the sample equals, for Ea this was 0.1511967578 mg/ul.
# 100% then divided by 100 to get 1% = 0.001511967578 mg/ul.
# 1% multiplied by % dose = 17.89 = 0.027049 mg/ul.
# 0.027049 mg/ul is then divided by woodlice median mass to get mg/kg (0.027049/0.00009 = 300.54 mg/kg)
# Repeat with the standard error dose to get standard range of error in mg/kg (50.23 mg/kg).


########################################
##### Let's try with 1hr peak data #####
########################################


#################################
### First, calculate ED50 1hr ###
#################################

### this subsets the data

E_atrica_ED50_cricket_1hr <- ED50_data_peaks_1hr[ED50_data_peaks_1hr$species == "Eratigena_atrica_EL" & ED50_data_peaks_1hr$prey_species == "Acheta_domesticus",]

model_E_atrica_ED50_cricket_1hr <- glm(cbind(E_atrica_ED50_cricket_1hr$incapacitated_or_dead, 
                                         E_atrica_ED50_cricket_1hr$alive) ~ (E_atrica_ED50_cricket_1hr$venom_conc_percent), 
                                         binomial(link = probit))
                                     
summary(model_E_atrica_ED50_cricket_1hr)

### plotting the data

ypredicts <- seq(0, 1, 0.01)
xpredicts <- dose.p(model_E_atrica_ED50_cricket_1hr,p=c(ypredicts))

plot(ED50_data_peaks_1hr$incapacitated_or_dead/20 ~ (ED50_data_peaks_1hr$venom_conc_percent),
     ylab = "Proportion Incapacitated",
     xlab = "Percent Dosage",
     pch = 16)

lines(unlist(xpredicts), ypredicts)


E_atrica_ED50_crickets_result_1hr <- dose.p(model_E_atrica_ED50_cricket_1hr, p=c(0.5))
E_atrica_ED50_crickets_result_1hr
#             Dose     SE
#    p = 0.5: 1.518181 0.560439


### Converting % dose to mg/kg ###

# The calculated dose is the % concentration dose (mg/ul) required to achieve an ED50 in crickets of median mass 0.0003 kg.
# To get ED50 mg/kg, you first figure out how much 100% of the sample equals (0.151197 mg/ul).
# 100% then divided by 100 to get 1% = 0.001512 mg/ul.
# 1% multiplied by % dose = 1.52 = 0.002298 mg/ul.
# 0.002298 mg/ul is then divided by cricket median mass to get mg/kg (0.002298/0.0003 = 7.66 mg/kg).
# Repeat with the standard error dose to get standard range of error in mg/kg (2.53 mg/kg).


### this subsets dataset (woodlice)

E_atrica_EL_ED50_woodlice_1hr <- ED50_data_peaks_1hr[ED50_data_peaks_1hr$species == "Eratigena_atrica_EL" & ED50_data_peaks_1hr$prey_species == "Porcellio_scaber", ]


model_E_atrica_EL_ED50_woodlice_1hr <- glm(cbind(E_atrica_EL_ED50_woodlice_1hr$incapacitated_or_dead, 
                                             E_atrica_EL_ED50_woodlice_1hr$alive) ~ (E_atrica_EL_ED50_woodlice_1hr$venom_conc_percent),
                                             binomial(link = probit))
                                         
summary(model_E_atrica_EL_ED50_woodlice_1hr)


### plotting the data

### this just generates a sequence of numbers
ypredicts <- seq(0, 1, 0.01)
xpredicts <- dose.p(model_E_atrica_EL_ED50_woodlice_1hr,p=c(ypredicts))


plot(E_atrica_EL_ED50_woodlice_1hr$incapacitated_or_dead/20 ~ (E_atrica_EL_ED50_woodlice_1hr$venom_conc_percent),
     ylab = "Proportion_incap",
     xlab = "Dosage",
     pch = 16)

lines(unlist(xpredicts), ypredicts)


E_atrica_EL_ED50_woodlice_result_1hr <- dose.p(model_E_atrica_EL_ED50_woodlice_1hr, p=c(0.5))
E_atrica_EL_ED50_woodlice_result_1hr
#                    Dose       SE
#     p = 0.5:   1.283907 1.540256


### Converting % dose to mg/kg ###

# The calculated dose is the % concentration dose (mg/ul) required to achieve an ED50 in woodlice of median mass 0.00009 kg.
# To get ED50 mg/kg, you first figure out how much 100% of the sample equals (0.151197 mg/ul).
# 100% then divided by 100 to get 1% = 0.001512 mg/ul.
# 1% multiplied by % dose = 1.28 = 0.001935 mg/ul.
# 0.001935 mg/ul is then divided by woodlice median mass to get mg/kg (0.001935/0.00009 = 21.5 mg/kg).
# Repeat with the standard error dose to get standard range of error in mg/kg (25.87 mg/kg).

```


```{r Heteropoda venatoria ED50 and LD50}

#############################
### First, calculate ED50 ###
#############################

### this subsets dataset

H_venatoria_ED50_crickets <- ED50_data_peaks[ED50_data_peaks$species == "Heteropoda_venatoria" & ED50_data_peaks$prey_species == "Acheta_domesticus", ]

model_H_venatoria_ED50_crickets <- glm(cbind(H_venatoria_ED50_crickets$incapacitated_or_dead, 
                                             H_venatoria_ED50_crickets$alive) ~ (H_venatoria_ED50_crickets$venom_conc_percent),
                                             binomial(link = probit))
 
summary(model_H_venatoria_ED50_crickets)


### plotting the data

### this just generates a sequence of numbers
ypredicts <- seq(0, 1, 0.01)
xpredicts <- dose.p(model_H_venatoria_ED50_crickets,p=c(ypredicts))


plot(H_venatoria_ED50_crickets$incapacitated_or_dead/20 ~ (H_venatoria_ED50_crickets$venom_conc_percent),
     ylab = "Proportion_incap",
     xlab = "Dosage",
     pch = 16)

lines(unlist(xpredicts), ypredicts)


H_venatoria_ED50_crickets_result <- dose.p(model_H_venatoria_ED50_crickets, p=c(0.5))
H_venatoria_ED50_crickets_result
#                  Dose       SE
#  p = 0.5: 0.2906368 0.04188438


### Converting % dose to mg/kg ###

# The calculated dose is the % concentration dose (mg/ul) required to achieve an ED50 in crickets of median mass 0.0003 kg.
# To get ED50 mg/kg, you first figure out how much 100% of the sample equals (0.1032547699214366 mg/ul).
# 100% then divided by 100 to get 1% = 0.001033 mg/ul.
# 1% multiplied by % dose = 0.29 = 0.000300 mg/ul.
# 0.000300 mg/ul is then divided by cricket median mass to get mg/kg (0.000300/0.0003 = 1.00 mg/kg).
# Repeat with the standard error dose to get standard range of error in mg/kg (0.14 mg/kg).


### this subsets dataset (woodlice)

H_venatoria_ED50_woodlice <- ED50_data_peaks[ED50_data_peaks$species == "Heteropoda_venatoria" & ED50_data_peaks$prey_species == "Porcellio_scaber", ]


model_H_venatoria_ED50_woodlice <- glm(cbind(H_venatoria_ED50_woodlice$incapacitated_or_dead, 
                                             H_venatoria_ED50_woodlice$alive) ~ (H_venatoria_ED50_woodlice$venom_conc_percent),
                                             binomial(link = probit))
                                         
summary(model_H_venatoria_ED50_woodlice)


### plotting the data

### this just generates a sequence of numbers
ypredicts <- seq(0, 1, 0.01)
xpredicts <- dose.p(model_H_venatoria_ED50_woodlice,p=c(ypredicts))


plot(H_venatoria_ED50_woodlice$incapacitated_or_dead/20 ~ (H_venatoria_ED50_woodlice$venom_conc_percent),
     ylab = "Proportion_incap",
     xlab = "Dosage",
     pch = 16)

lines(unlist(xpredicts), ypredicts)


H_venatoria_ED50_woodlice_result <- dose.p(model_H_venatoria_ED50_woodlice, p=c(0.5))
H_venatoria_ED50_woodlice_result
#                     Dose       SE
#         p = 0.5: 1.48095 0.7053408


### Converting % dose to mg/kg ###

# The calculated dose is the % concentration dose (mg/ul) required to achieve an ED50 in woodlice of median mass 0.00009 kg.
# To get ED50 mg/kg, you first figure out how much 100% of the sample equals (0.1032547699214366 mg/ul).
# 100% then divided by 100 to get 1% = 0.001033 mg/ul.
# 1% multiplied by % dose = 1.48 = 0.001529 mg/ul.
# 0.001529 mg/ul is then divided by woodlice median mass to get mg/kg (0.001529/0.00009 = 16.99 mg/kg).
# Repeat with the standard error dose to get standard range of error in mg/kg (8.14 mg/kg).


##########################
### Now calculate LD50 ###
##########################


### this subsets dataset

H_venatoria_LD50_crickets <- LD50_data24[LD50_data24$species == "Heteropoda_venatoria" & LD50_data24$prey_species == "Acheta_domesticus", ]

model_H_venatoria_LD50_crickets <- glm(cbind(H_venatoria_LD50_crickets$dead, 
                                             H_venatoria_LD50_crickets$alive) ~ (H_venatoria_LD50_crickets$venom_conc_percent), 
                                             binomial(link = probit))
                                         
summary(model_H_venatoria_LD50_crickets)


### plotting the data

### this just generates a sequence of numbers
ypredicts <- seq(0, 1, 0.01)
xpredicts <- dose.p(model_H_venatoria_LD50_crickets,p=c(ypredicts))


plot(H_venatoria_LD50_crickets$dead/20 ~ (H_venatoria_LD50_crickets$venom_conc_percent),
     ylab = "Proportion_dead",
     xlab = "Dosage",
     pch = 16)

lines(unlist(xpredicts), ypredicts)

### LD50 ###
H_venatoria_LD50_crickets_result <- dose.p(model_H_venatoria_LD50_crickets, p=c(0.5))
H_venatoria_LD50_crickets_result
#          Dose     SE
# p = 0.5: 10.93444 1.93649


### Converting % dose to mg/kg ###

# The calculated dose is the % concentration dose (mg/ul) required to achieve an LD50 in crickets of median mass 0.0003 kg.
# To get LD50 mg/kg, you first figure out how much 100% of the sample equals (0.1032547699214366 mg/ul).
# 100% then divided by 100 to get 1% = 0.001033 mg/ul.
# 1% multiplied by % dose = 10.93 = 0.011291 mg/ul.
# 0.011291 mg/ul is then divided by crickets median mass to get mg/kg (0.011291/0.0003 = 37.64 mg/kg).
# Repeat with the standard error dose to get standard range of error in mg/kg (6.68 mg/kg).


### this subsets dataset (woodlice)

H_venatoria_LD50_woodlice <- LD50_data24[LD50_data24$species == "Heteropoda_venatoria" & LD50_data24$prey_species == "Porcellio_scaber", ]


model_H_venatoria_LD50_woodlice <- glm(cbind(H_venatoria_LD50_woodlice$dead, 
                                             H_venatoria_LD50_woodlice$alive) ~ (H_venatoria_LD50_woodlice$venom_conc_percent),
                                             binomial(link = probit))
                                         
summary(model_H_venatoria_LD50_woodlice)


### plotting the data

### this just generates a sequence of numbers
ypredicts <- seq(0, 1, 0.01)
xpredicts <- dose.p(model_H_venatoria_LD50_woodlice,p=c(ypredicts))


plot(H_venatoria_LD50_woodlice$dead/20 ~ (H_venatoria_LD50_woodlice$venom_conc_percent),
     ylab = "Proportion_dead",
     xlab = "Dosage",
     pch = 16)

lines(unlist(xpredicts), ypredicts)

### LD50 ###
H_venatoria_LD50_woodlice_result <- dose.p(model_H_venatoria_LD50_woodlice, p=c(0.5))
H_venatoria_LD50_woodlice_result
#          Dose     SE
# p = 0.5: 11.45773 4.983277


### Converting % dose to mg/kg ###

# The calculated dose is the % concentration dose (mg/ul) required to achieve an LD50 in woodlice of median mass 0.00009 kg.
# To get LD50 mg/kg, you first figure out how much 100% of the sample equals (0.1032547699214366 mg/ul).
# 100% then divided by 100 to get 1% = 0.001033 mg/ul.
# 1% multiplied by % dose = 11.46 = 0.011838 mg/ul.
# 0.011838 mg/ul is then divided by woodlice median mass to get mg/kg (0.011838/0.00009 = 131.53 mg/kg)
# Repeat with the standard error dose to get standard range of error in mg/kg (57.16 mg/kg).


########################################
##### Let's try with 1hr peak data #####
########################################


#################################
### First, calculate ED50 1hr ###
#################################

### this subsets dataset

H_venatoria_ED50_crickets_1hr <- ED50_data_peaks_1hr[ED50_data_peaks_1hr$species == "Heteropoda_venatoria" & ED50_data_peaks_1hr$prey_species == "Acheta_domesticus", ]

model_H_venatoria_ED50_crickets_1hr <- glm(cbind(H_venatoria_ED50_crickets_1hr$incapacitated_or_dead, 
                                             H_venatoria_ED50_crickets_1hr$alive) ~ (H_venatoria_ED50_crickets_1hr$venom_conc_percent),
                                             binomial(link = probit))
 
summary(model_H_venatoria_ED50_crickets_1hr)


### plotting the data

### this just generates a sequence of numbers
ypredicts <- seq(0, 1, 0.01)
xpredicts <- dose.p(model_H_venatoria_ED50_crickets_1hr,p=c(ypredicts))


plot(H_venatoria_ED50_crickets_1hr$incapacitated_or_dead/20 ~ (H_venatoria_ED50_crickets_1hr$venom_conc_percent),
     ylab = "Proportion_incap",
     xlab = "Dosage",
     pch = 16)

lines(unlist(xpredicts), ypredicts)


H_venatoria_ED50_crickets_result_1hr <- dose.p(model_H_venatoria_ED50_crickets_1hr, p=c(0.5))
H_venatoria_ED50_crickets_result_1hr
#                  Dose       SE
#  p = 0.5: 0.2906368 0.04188438


### Converting % dose to mg/kg ###

# The calculated dose is the % concentration dose (mg/ul) required to achieve an ED50 in crickets of median mass 0.0003 kg.
# To get ED50 mg/kg, you first figure out how much 100% of the sample equals (0.1032547699214366 mg/ul).
# 100% then divided by 100 to get 1% = 0.001033 mg/ul.
# 1% multiplied by % dose = 0.29 = 0.000300 mg/ul.
# 0.000300 mg/ul is then divided by cricket median mass to get mg/kg (0.000300/0.0003 = 1.00 mg/kg).
# Repeat with the standard error dose to get standard range of error in mg/kg (0.14 mg/kg).


### this subsets dataset (woodlice)

H_venatoria_ED50_woodlice_1hr <- ED50_data_peaks_1hr[ED50_data_peaks_1hr$species == "Heteropoda_venatoria" & ED50_data_peaks_1hr$prey_species == "Porcellio_scaber", ]


model_H_venatoria_ED50_woodlice_1hr <- glm(cbind(H_venatoria_ED50_woodlice_1hr$incapacitated_or_dead, 
                                             H_venatoria_ED50_woodlice_1hr$alive) ~ (H_venatoria_ED50_woodlice_1hr$venom_conc_percent),
                                             binomial(link = probit))
                                         
summary(model_H_venatoria_ED50_woodlice_1hr)


### plotting the data

### this just generates a sequence of numbers
ypredicts <- seq(0, 1, 0.01)
xpredicts <- dose.p(model_H_venatoria_ED50_woodlice_1hr,p=c(ypredicts))


plot(H_venatoria_ED50_woodlice_1hr$incapacitated_or_dead/20 ~ (H_venatoria_ED50_woodlice_1hr$venom_conc_percent),
     ylab = "Proportion_incap",
     xlab = "Dosage",
     pch = 16)

lines(unlist(xpredicts), ypredicts)


H_venatoria_ED50_woodlice_result_1hr <- dose.p(model_H_venatoria_ED50_woodlice_1hr, p=c(0.5))
H_venatoria_ED50_woodlice_result_1hr
#                  Dose     SE
#         p = 0.5: 1.881523 0.7252144


### Converting % dose to mg/kg ###

# The calculated dose is the % concentration dose (mg/ul) required to achieve an ED50 in woodlice of median mass 0.00009 kg.
# To get ED50 mg/kg, you first figure out how much 100% of the sample equals (0.1032547699214366 mg/ul).
# 100% then divided by 100 to get 1% = 0.001033 mg/ul.
# 1% multiplied by % dose = 1.88 = 0.001942 mg/ul.
# 0.001942 mg/ul is then divided by woodlice median mass to get mg/kg (0.001942/0.00009 = 21.58 mg/kg).
# Repeat with the standard error dose to get standard range of error in mg/kg (8.32 mg/kg).

```


```{r Larinioides sclopetarius ED50 and LD50}

#############################
### First, calculate ED50 ###
#############################

### this subsets dataset

L_sclopetarius_ED50_crickets <- ED50_data_peaks[ED50_data_peaks$species == "Larinioides_sclopetarius" & ED50_data_peaks$prey_species == "Acheta_domesticus", ] 

model_L_sclopetarius_ED50_crickets <- glm(cbind(L_sclopetarius_ED50_crickets$incapacitated_or_dead, 
                                                L_sclopetarius_ED50_crickets$alive) ~ (L_sclopetarius_ED50_crickets$venom_conc_percent), 
                                                binomial(link = probit))
                                            
summary(model_L_sclopetarius_ED50_crickets)


### plotting the data

### this just generates a sequence of numbers
ypredicts <- seq(0, 1, 0.01)
xpredicts <- dose.p(model_L_sclopetarius_ED50_crickets,p=c(ypredicts))


plot(L_sclopetarius_ED50_crickets$incapacitated_or_dead/20 ~ (L_sclopetarius_ED50_crickets$venom_conc_percent),
     ylab = "Proportion_incap",
     xlab = "Dosage",
     pch = 16)

lines(unlist(xpredicts), ypredicts)


L_sclopetarius_ED50_crickets_result <- dose.p(model_L_sclopetarius_ED50_crickets, p=c(0.5))
L_sclopetarius_ED50_crickets_result
#                 Dose       SE
# p = 0.5: 0.3592372 0.05942168


### Converting % dose to mg/kg ###

# The calculated dose is the % concentration dose (mg/ul) required to achieve an ED50 in crickets of median mass 0.0003 kg.
# To get ED50 mg/kg, you first figure out how much 100% of the sample equals (0.2315202231520223 mg/ul).
# 100% then divided by 100 to get 1% = 0.002315 mg/ul.
# 1% multiplied by % dose = 0.36 = 0.000833 mg/ul.
# 0.000833 mg/ul is then divided by cricket median mass to get mg/kg (0.000833/0.0003 = 2.78 mg/kg).
# Repeat with the standard error dose to get standard range of error in mg/kg (0.46 mg/kg).


### this subsets dataset (woodlice)

L_sclopetarius_ED50_woodlice <- ED50_data_peaks[ED50_data_peaks$species == "Larinioides_sclopetarius" & ED50_data_peaks$prey_species == "Porcellio_scaber", ]


model_L_sclopetarius_ED50_woodlice <- glm(cbind(L_sclopetarius_ED50_woodlice$incapacitated_or_dead, 
                                                L_sclopetarius_ED50_woodlice$alive) ~ (L_sclopetarius_ED50_woodlice$venom_conc_percent),
                                                binomial(link = probit))
                                            
summary(model_L_sclopetarius_ED50_woodlice)


### plotting the data

### this just generates a sequence of numbers
ypredicts <- seq(0, 1, 0.01)
xpredicts <- dose.p(model_L_sclopetarius_ED50_woodlice,p=c(ypredicts))


plot(L_sclopetarius_ED50_woodlice$incapacitated_or_dead/20 ~ (L_sclopetarius_ED50_woodlice$venom_conc_percent),
     ylab = "Proportion_incap",
     xlab = "Dosage",
     pch = 16)

lines(unlist(xpredicts), ypredicts)


L_sclopetarius_ED50_woodlice_result <- dose.p(model_L_sclopetarius_ED50_woodlice, p=c(0.5))
L_sclopetarius_ED50_woodlice_result
#               Dose        SE
# p = 0.5:  1.021453 0.1340273


### Converting % dose to mg/kg ###

# The calculated dose is the % concentration dose (mg/ul) required to achieve an ED50 in woodlice of median mass 0.00009 kg.
# To get ED50 mg/kg, you first figure out how much 100% of the sample equals (0.2315202231520223 mg/ul).
# 100% then divided by 100 to get 1% = 0.002315 mg/ul.
# 1% multiplied by % dose = 1.02 = 0.002361 mg/ul.
# 0.002361 mg/ul is then divided by woodlice median mass to get mg/kg (0.002361/0.00009 = 26.23 mg/kg).
# Repeat with the standard error dose to get standard range of error in mg/kg (3.34 mg/kg).


##########################
### Now calculate LD50 ###
##########################

### this subsets dataset

L_sclopetarius_LD50_crickets <- LD50_data24[LD50_data24$species == "Larinioides_sclopetarius" & LD50_data24$prey_species == "Acheta_domesticus", ]

model_L_sclopetarius_LD50_crickets <- glm(cbind(L_sclopetarius_LD50_crickets$dead, 
                                                L_sclopetarius_LD50_crickets$alive) ~ (L_sclopetarius_LD50_crickets$venom_conc_percent), 
                                                binomial(link = probit))
                                            
summary(model_L_sclopetarius_LD50_crickets)


### plotting the data

### this just generates a sequence of numbers
ypredicts <- seq(0, 1, 0.01)
xpredicts <- dose.p(model_L_sclopetarius_LD50_crickets,p=c(ypredicts))


plot(L_sclopetarius_LD50_crickets$dead/20 ~ (L_sclopetarius_LD50_crickets$venom_conc_percent),
     ylab = "Proportion_dead",
     xlab = "Dosage",
     pch = 16)

lines(unlist(xpredicts), ypredicts)

### LD50 ###
L_sclopetarius_LD50_crickets_result <- dose.p(model_L_sclopetarius_LD50_crickets, p=c(0.5))
L_sclopetarius_LD50_crickets_result
#          Dose     SE
# p = 0.5: 1.963493 0.8892865


### Converting % dose to mg/kg ###

# The calculated dose is the % concentration dose (mg/ul) required to achieve an LD50 in crickets of median mass 0.0003 kg.
# To get LD50 mg/kg, you first figure out how much 100% of the sample equals (0.2315202231520223 mg/ul).
# 100% then divided by 100 to get 1% = 0.002315 mg/ul.
# 1% multiplied by % dose = 1.96 = 0.004537 mg/ul.
# 0.004537 mg/ul is then divided by crickets median mass to get mg/kg (0.004537/0.0003 = 15.12 mg/kg).
# Repeat with the standard error dose to get standard range of error in mg/kg (6.87 mg/kg).


### this subsets dataset (woodlice)

L_sclopetarius_LD50_woodlice <- LD50_data24[LD50_data24$species == "Larinioides_sclopetarius" & LD50_data24$prey_species == "Porcellio_scaber", ]


model_L_sclopetarius_LD50_woodlice <- glm(cbind(L_sclopetarius_LD50_woodlice$dead, 
                                                L_sclopetarius_LD50_woodlice$alive) ~ (L_sclopetarius_LD50_woodlice$venom_conc_percent),
                                                binomial(link = probit))
                                            
summary(model_L_sclopetarius_LD50_woodlice)


### plotting the data

### this just generates a sequence of numbers
ypredicts <- seq(0, 1, 0.01)
xpredicts <- dose.p(model_L_sclopetarius_LD50_woodlice,p=c(ypredicts))


plot(L_sclopetarius_LD50_woodlice$dead/20 ~ (L_sclopetarius_LD50_woodlice$venom_conc_percent),
     ylab = "Proportion_dead",
     xlab = "Dosage",
     pch = 16)

lines(unlist(xpredicts), ypredicts)


### LD50 ###
L_sclopetarius_LD50_woodlice_result <- dose.p(model_L_sclopetarius_LD50_woodlice, p=c(0.5))
L_sclopetarius_LD50_woodlice_result
#          Dose     SE
# p = 0.5: 1.446297 0.5916746


### Converting % dose to mg/kg ###

# The calculated dose is the % concentration dose (mg/ul) required to achieve an LD50 in woodlice of median mass 0.00009 kg.
# To get LD50 mg/kg, you first figure out how much 100% of the sample equals (0.2315202231520223 mg/ul).
# 100% then divided by 100 to get 1% = 0.002315 mg/ul.
# 1% multiplied by % dose = 1.45 = 0.003357 mg/ul.
# 0.003357 mg/ul is then divided by woodlice median mass to get mg/kg (0.003357/0.00009 = 37.3 mg/kg).
# Repeat with the standard error dose to get standard range of error in mg/kg (15.18 mg/kg).


########################################
##### Let's try with 1hr peak data #####
########################################


#################################
### First, calculate ED50 1hr ###
#################################

### this subsets dataset

L_sclopetarius_ED50_crickets_1hr <- ED50_data_peaks_1hr[ED50_data_peaks_1hr$species == "Larinioides_sclopetarius" & ED50_data_peaks_1hr$prey_species == "Acheta_domesticus", ] 

model_L_sclopetarius_ED50_crickets_1hr <- glm(cbind(L_sclopetarius_ED50_crickets_1hr$incapacitated_or_dead, 
                                                L_sclopetarius_ED50_crickets_1hr$alive) ~ (L_sclopetarius_ED50_crickets_1hr$venom_conc_percent), 
                                                binomial(link = probit))
                                            
summary(model_L_sclopetarius_ED50_crickets_1hr)


### plotting the data

### this just generates a sequence of numbers
ypredicts <- seq(0, 1, 0.01)
xpredicts <- dose.p(model_L_sclopetarius_ED50_crickets_1hr,p=c(ypredicts))


plot(L_sclopetarius_ED50_crickets_1hr$incapacitated_or_dead/20 ~ (L_sclopetarius_ED50_crickets_1hr$venom_conc_percent),
     ylab = "Proportion_incap",
     xlab = "Dosage",
     pch = 16)

lines(unlist(xpredicts), ypredicts)


L_sclopetarius_ED50_crickets_result_1hr <- dose.p(model_L_sclopetarius_ED50_crickets_1hr, p=c(0.5))
L_sclopetarius_ED50_crickets_result_1hr
#                 Dose       SE
# p = 0.5: 0.3592372 0.05942168


### Converting % dose to mg/kg ###

# The calculated dose is the % concentration dose (mg/ul) required to achieve an ED50 in crickets of median mass 0.0003 kg.
# To get ED50 mg/kg, you first figure out how much 100% of the sample equals (0.2315202231520223 mg/ul).
# 100% then divided by 100 to get 1% = 0.002315 mg/ul.
# 1% multiplied by % dose = 0.36 = 0.000833 mg/ul.
# 0.000833 mg/ul is then divided by cricket median mass to get mg/kg (0.000833/0.0003 = 2.78 mg/kg).
# Repeat with the standard error dose to get standard range of error in mg/kg (0.46 mg/kg).


### this subsets dataset (woodlice)

L_sclopetarius_ED50_woodlice_1hr <- ED50_data_peaks_1hr[ED50_data_peaks_1hr$species == "Larinioides_sclopetarius" & ED50_data_peaks_1hr$prey_species == "Porcellio_scaber", ]


model_L_sclopetarius_ED50_woodlice_1hr <- glm(cbind(L_sclopetarius_ED50_woodlice_1hr$incapacitated_or_dead, 
                                                L_sclopetarius_ED50_woodlice_1hr$alive) ~ (L_sclopetarius_ED50_woodlice_1hr$venom_conc_percent),
                                                binomial(link = probit))
                                            
summary(model_L_sclopetarius_ED50_woodlice_1hr)


### plotting the data

### this just generates a sequence of numbers
ypredicts <- seq(0, 1, 0.01)
xpredicts <- dose.p(model_L_sclopetarius_ED50_woodlice_1hr,p=c(ypredicts))


plot(L_sclopetarius_ED50_woodlice_1hr$incapacitated_or_dead/20 ~ (L_sclopetarius_ED50_woodlice_1hr$venom_conc_percent),
     ylab = "Proportion_incap",
     xlab = "Dosage",
     pch = 16)

lines(unlist(xpredicts), ypredicts)


L_sclopetarius_ED50_woodlice_result_1hr <- dose.p(model_L_sclopetarius_ED50_woodlice_1hr, p=c(0.5))
L_sclopetarius_ED50_woodlice_result_1hr
#           Dose     SE
# p = 0.5:  1.093098 0.1360672


### Converting % dose to mg/kg ###

# The calculated dose is the % concentration dose (mg/ul) required to achieve an ED50 in woodlice of median mass 0.00009 kg.
# To get ED50 mg/kg, you first figure out how much 100% of the sample equals (0.2315202231520223 mg/ul).
# 100% then divided by 100 to get 1% = 0.002315 mg/ul.
# 1% multiplied by % dose = 1.09 = 0.002523 mg/ul.
# 0.002523 mg/ul is then divided by woodlice median mass to get mg/kg (0.002523/0.00009 = 28.03 mg/kg).
# Repeat with the standard error dose to get standard range of error in mg/kg (3.5 mg/kg).
```


```{r Meta menardi ED50 and LD50}

#############################
### First, calculate ED50 ###
#############################

### this subsets dataset

M_menardi_ED50_crickets <- ED50_data_peaks[ED50_data_peaks$species == "Meta_menardi" & ED50_data_peaks$prey_species == "Acheta_domesticus", ]

model_M_menardi_ED50_crickets <- glm(cbind(M_menardi_ED50_crickets$incapacitated_or_dead, 
                                           M_menardi_ED50_crickets$alive) ~ (M_menardi_ED50_crickets$venom_conc_percent), 
                                           binomial(link = probit))
                                       
summary(model_M_menardi_ED50_crickets)


### plotting the data

### this just generates a sequence of numbers
ypredicts <- seq(0, 1, 0.01)
xpredicts <- dose.p(model_M_menardi_ED50_crickets,p=c(ypredicts))


plot(M_menardi_ED50_crickets$incapacitated_or_dead/20 ~ (M_menardi_ED50_crickets$venom_conc_percent),
     ylab = "Proportion_incap",
     xlab = "Dosage",
     pch = 16)

lines(unlist(xpredicts), ypredicts)


M_menardi_ED50_crickets_result <- dose.p(model_M_menardi_ED50_crickets, p=c(0.5))
M_menardi_ED50_crickets_result
#                      Dose       SE
#     p = 0.5: 0.2929342 0.085313732


### Converting % dose to mg/kg ###

# The calculated dose is the % concentration dose (mg/ul) required to achieve an ED50 in crickets of median mass 0.0003 kg.
# To get ED50 mg/kg, you first figure out how much 100% of the sample equals (0.0761904761904762 mg/ul).
# 100% then divided by 100 to get 1% = 0.000762 mg/ul.
# 1% multiplied by % dose = 0.29 = 0.000221 mg/ul.
# 0.000221 mg/ul is then divided by cricket median mass to get mg/kg (0.000221/0.0003 = 0.74 mg/kg).
# Repeat with the standard error dose to get standard range of error in mg/kg (0.20 mg/kg).


### this subsets dataset (woodlice)

M_menardi_ED50_woodlice <- ED50_data_peaks[ED50_data_peaks$species == "Meta_menardi" & ED50_data_peaks$prey_species == "Porcellio_scaber", ]


model_M_menardi_ED50_woodlice <- glm(cbind(M_menardi_ED50_woodlice$incapacitated_or_dead, 
                                           M_menardi_ED50_woodlice$alive) ~ (M_menardi_ED50_woodlice$venom_conc_percent),
                                           binomial(link = probit))
                                       
summary(model_M_menardi_ED50_woodlice)


### plotting the data

### this just generates a sequence of numbers
ypredicts <- seq(0, 1, 0.01)
xpredicts <- dose.p(model_M_menardi_ED50_woodlice,p=c(ypredicts))


plot(M_menardi_ED50_woodlice$incapacitated_or_dead/20 ~ (M_menardi_ED50_woodlice$venom_conc_percent),
     ylab = "Proportion_incap",
     xlab = "Dosage",
     pch = 16)

lines(unlist(xpredicts), ypredicts)


M_menardi_ED50_woodlice_result <- dose.p(model_M_menardi_ED50_woodlice, p=c(0.5))
M_menardi_ED50_woodlice_result
#                   Dose        SE
#     p = 0.5: 0.3788829 0.3391901


### Converting % dose to mg/kg ###

# The calculated dose is the % concentration dose (mg/ul) required to achieve an ED50 in woodlice of median mass 0.00009 kg.
# To get ED50 mg/kg, you first figure out how much 100% of the sample equals (0.0761904761904762 mg/ul).
# 100% then divided by 100 to get 1% = 0.000762 mg/ul.
# 1% multiplied by % dose = 0.38 = 0.000290 mg/ul.
# 0.000290 mg/ul is then divided by woodlice median mass to get mg/kg (0.000290/0.00009 = 3.22 mg/kg).
# Repeat with the standard error dose to get standard range of error in mg/kg (2.88 mg/kg).


##########################
### Now calculate LD50 ###
##########################

### this subsets dataset

M_menardi_LD50_crickets <- LD50_data24[LD50_data24$species == "Meta_menardi" & LD50_data24$prey_species == "Acheta_domesticus", ]

model_M_menardi_LD50_crickets <- glm(cbind(M_menardi_LD50_crickets$dead, 
                                           M_menardi_LD50_crickets$alive) ~ (M_menardi_LD50_crickets$venom_conc_percent), 
                                           binomial(link = probit))
                                       
summary(model_M_menardi_LD50_crickets)


### plotting the data

### this just generates a sequence of numbers
ypredicts <- seq(0, 1, 0.01)
xpredicts <- dose.p(model_M_menardi_LD50_crickets,p=c(ypredicts))


plot(M_menardi_LD50_crickets$dead/20 ~ (M_menardi_LD50_crickets$venom_conc_percent),
     ylab = "Proportion_dead",
     xlab = "Dosage",
     pch = 16)

lines(unlist(xpredicts), ypredicts)


### LD50 ###
M_menardi_LD50_crickets_result <- dose.p(model_M_menardi_LD50_crickets, p=c(0.5))
M_menardi_LD50_crickets_result
#          Dose     SE
# p = 0.5: 4.907392 1.078293


### Converting % dose to mg/kg ###

# The calculated dose is the % concentration dose (mg/ul) required to achieve an LD50 in crickets of median mass 0.0003 kg.
# To get LD50 mg/kg, you first figure out how much 100% of the sample equals (0.076190 mg/ul).
# 100% then divided by 100 to get 1% = 0.000762 mg/ul.
# 1% multiplied by % dose = 4.91 = 0.003741 mg/ul.
# 0.003741 mg/ul is then divided by crickets median mass to get mg/kg (0.003741/0.0003 = 12.47 mg/kg).
# Repeat with the standard error dose to get standard range of error in mg/kg (2.74 mg/kg).


### this subsets dataset (woodlice)

M_menardi_LD50_woodlice <- LD50_data24[LD50_data24$species == "Meta_menardi" & LD50_data24$prey_species == "Porcellio_scaber", ]


model_M_menardi_LD50_woodlice <- glm(cbind(M_menardi_LD50_woodlice$dead, 
                                           M_menardi_LD50_woodlice$alive) ~ (M_menardi_LD50_woodlice$venom_conc_percent),
                                           binomial(link = probit))
                                       
summary(model_M_menardi_LD50_woodlice)


### plotting the data

### this just generates a sequence of numbers
ypredicts <- seq(0, 1, 0.01)
xpredicts <- dose.p(model_M_menardi_LD50_woodlice,p=c(ypredicts))


plot(M_menardi_LD50_woodlice$dead/20 ~ (M_menardi_LD50_woodlice$venom_conc_percent),
     ylab = "Proportion_dead",
     xlab = "Dosage",
     pch = 16)

lines(unlist(xpredicts), ypredicts)


### LD50 ###
M_menardi_LD50_woodlice_result <- dose.p(model_M_menardi_LD50_woodlice, p=c(0.5))
M_menardi_LD50_woodlice_result
#          Dose     SE
# p = 0.5: 8.855659 6.913721


### Converting % dose to mg/kg ###

# The calculated dose is the % concentration dose (mg/ul) required to achieve an LD50 in woodlice of median mass 0.00009 kg.
# To get LD50 mg/kg, you first figure out how much 100% of the sample equals (0.076190 mg/ul).
# 100% then divided by 100 to get 1% = 0.000762 mg/ul.
# 1% multiplied by % dose = 8.86 = 0.006751 mg/ul.
# 0.006751 mg/ul is then divided by woodlice median mass to get mg/kg (0.006751/0.00009 = 75.01 mg/kg).
# Repeat with the standard error dose to get standard range of error in mg/kg (58.5 mg/kg).


########################################
##### Let's try with 1hr peak data #####
########################################


#################################
### First, calculate ED50 1hr ###
#################################

### this subsets dataset

M_menardi_ED50_crickets_1hr <- ED50_data_peaks_1hr[ED50_data_peaks_1hr$species == "Meta_menardi" & ED50_data_peaks_1hr$prey_species == "Acheta_domesticus", ]

model_M_menardi_ED50_crickets_1hr <- glm(cbind(M_menardi_ED50_crickets_1hr$incapacitated_or_dead, 
                                           M_menardi_ED50_crickets_1hr$alive) ~ (M_menardi_ED50_crickets_1hr$venom_conc_percent), 
                                           binomial(link = probit))
                                       
summary(model_M_menardi_ED50_crickets_1hr)


### plotting the data

### this just generates a sequence of numbers
ypredicts <- seq(0, 1, 0.01)
xpredicts <- dose.p(model_M_menardi_ED50_crickets_1hr,p=c(ypredicts))


plot(M_menardi_ED50_crickets_1hr$incapacitated_or_dead/20 ~ (M_menardi_ED50_crickets_1hr$venom_conc_percent),
     ylab = "Proportion_incap",
     xlab = "Dosage",
     pch = 16)

lines(unlist(xpredicts), ypredicts)


M_menardi_ED50_crickets_result_1hr <- dose.p(model_M_menardi_ED50_crickets_1hr, p=c(0.5))
M_menardi_ED50_crickets_result_1hr
#                      Dose       SE
#      p = 0.5: 0.3178383 0.07945507


### Converting % dose to mg/kg ###

# The calculated dose is the % concentration dose (mg/ul) required to achieve an ED50 in crickets of median mass 0.0003 kg.
# To get ED50 mg/kg, you first figure out how much 100% of the sample equals (0.0761904761904762 mg/ul).
# 100% then divided by 100 to get 1% = 0.000762 mg/ul.
# 1% multiplied by % dose = 0.32 = 0.0002438 mg/ul.
# 0.0002438 mg/ul is then divided by cricket median mass to get mg/kg (0.0002438/0.0003 = 0.81 mg/kg).
# Repeat with the standard error dose to get standard range of error in mg/kg (0.20 mg/kg).


### this subsets dataset (woodlice)

M_menardi_ED50_woodlice_1hr <- ED50_data_peaks_1hr[ED50_data_peaks_1hr$species == "Meta_menardi" & ED50_data_peaks_1hr$prey_species == "Porcellio_scaber", ]


model_M_menardi_ED50_woodlice_1hr <- glm(cbind(M_menardi_ED50_woodlice_1hr$incapacitated_or_dead, 
                                           M_menardi_ED50_woodlice_1hr$alive) ~ (M_menardi_ED50_woodlice_1hr$venom_conc_percent),
                                           binomial(link = probit))
                                       
summary(model_M_menardi_ED50_woodlice_1hr)


### plotting the data

### this just generates a sequence of numbers
ypredicts <- seq(0, 1, 0.01)
xpredicts <- dose.p(model_M_menardi_ED50_woodlice_1hr,p=c(ypredicts))


plot(M_menardi_ED50_woodlice_1hr$incapacitated_or_dead/20 ~ (M_menardi_ED50_woodlice_1hr$venom_conc_percent),
     ylab = "Proportion_incap",
     xlab = "Dosage",
     pch = 16)

lines(unlist(xpredicts), ypredicts)


M_menardi_ED50_woodlice_result_1hr <- dose.p(model_M_menardi_ED50_woodlice_1hr, p=c(0.5))
M_menardi_ED50_woodlice_result_1hr
#              Dose      SE
#     p = 0.5: 0.3684871 0.4092125


### Converting % dose to mg/kg ###

# The calculated dose is the % concentration dose (mg/ul) required to achieve an ED50 in woodlice of median mass 0.00009 kg.
# To get ED50 mg/kg, you first figure out how much 100% of the sample equals (0.0761904761904762 mg/ul).
# 100% then divided by 100 to get 1% = 0.000762 mg/ul.
# 1% multiplied by % dose = 0.37 = 0.0002819 mg/ul.
# 0.0002819 mg/ul is then divided by woodlice median mass to get mg/kg (0.0002819/0.00009 = 3.13 mg/kg).
# Repeat with the standard error dose to get standard range of error in mg/kg (3.46 mg/kg).

```


```{r Monocentropus balfouri ED50 and LD50}

#############################
### First, calculate ED50 ###
#############################

### this subsets dataset

M_balfouri_ED50_crickets <- ED50_data_peaks[ED50_data_peaks$species == "Monocentropus_balfouri" & ED50_data_peaks$prey_species == "Acheta_domesticus", ]

model_M_balfouri_ED50_crickets <- glm(cbind(M_balfouri_ED50_crickets$incapacitated_or_dead, 
                                            M_balfouri_ED50_crickets$alive) ~ (M_balfouri_ED50_crickets$venom_conc_percent), 
                                            binomial(link = probit))
                                        
summary(model_M_balfouri_ED50_crickets)


##### plotting the data

#this just generates a sequence of numbers
ypredicts <- seq(0, 1, 0.01)
xpredicts <- dose.p(model_M_balfouri_ED50_crickets,p=c(ypredicts))


plot(M_balfouri_ED50_crickets$incapacitated_or_dead/20 ~ (M_balfouri_ED50_crickets$venom_conc_percent),
     ylab = "Proportion_incap",
     xlab = "Dosage",
     pch = 16)

lines(unlist(xpredicts), ypredicts)


M_balfouri_ED50_crickets_result <- dose.p(model_M_balfouri_ED50_crickets, p=c(0.5))
M_balfouri_ED50_crickets_result
#                  Dose        SE
#    p = 0.5: 0.8050086 0.2500554


### Converting % dose to mg/kg ###

# The calculated dose is the % concentration dose (mg/ul) required to achieve an ED50 in crickets of median mass 0.0003 kg.
# To get ED50 mg/kg, you first figure out how much 100% of the sample equals (0.1105527638190955 mg/ul).
# 100% then divided by 100 to get 1% = 0.001105 mg/ul.
# 1% multiplied by % dose = 0.81 = 0.000895 mg/ul.
# 0.000895 mg/ul is then divided by cricket median mass to get mg/kg (0.000895/0.0003 = 2.98 mg/kg).
# Repeat with the standard error dose to get standard range of error in mg/kg (0.92 mg/kg).


### this subsets dataset (woodlice)

M_balfouri_ED50_woodlice <- ED50_data_peaks[ED50_data_peaks$species == "Monocentropus_balfouri" & ED50_data_peaks$prey_species == "Porcellio_scaber", ]


model_M_balfouri_ED50_woodlice <- glm(cbind(M_balfouri_ED50_woodlice$incapacitated_or_dead, 
                                            M_balfouri_ED50_woodlice$alive) ~ (M_balfouri_ED50_woodlice$venom_conc_percent),
                                            binomial(link = probit))
                                     
summary(model_M_balfouri_ED50_woodlice)


### plotting the data

### this just generates a sequence of numbers
ypredicts <- seq(0, 1, 0.01)
xpredicts <- dose.p(model_M_balfouri_ED50_woodlice,p=c(ypredicts))


plot(M_balfouri_ED50_woodlice$incapacitated_or_dead/20 ~ (M_balfouri_ED50_woodlice$venom_conc_percent),
     ylab = "Proportion_incap",
     xlab = "Dosage",
     pch = 16)

lines(unlist(xpredicts), ypredicts)


M_balfouri_ED50_woodlice_result <- dose.p(model_M_balfouri_ED50_woodlice, p=c(0.5))
M_balfouri_ED50_woodlice_result
#                  Dose        SE
#    p = 0.5: 0.6586176 0.5243086


### Converting % dose to mg/kg ###

# The calculated dose is the % concentration dose (mg/ul) required to achieve an ED50 in woodlice of median mass 0.00009 kg.
# To get ED50 mg/kg, you first figure out how much 100% of the sample equals (0.1105527638190955 mg/ul).
# 100% then divided by 100 to get 1% = 0.001105 mg/ul.
# 1% multiplied by % dose = 0.66 = 0.000729 mg/ul.
# 0.000729 mg/ul is then divided by woodlice median mass to get mg/kg (0.000729/0.00009 = 8.1 mg/kg).
# Repeat with the standard error dose to get standard range of error in mg/kg (6.39 mg/kg).


##########################
### Now calculate LD50 ###
##########################


M_balfouri_LD50_crickets <- LD50_data24[LD50_data24$species == "Monocentropus_balfouri" & LD50_data24$prey_species == "Acheta_domesticus", ]

model_M_balfouri_LD50_crickets <- glm(cbind(M_balfouri_LD50_crickets$dead, 
                                            M_balfouri_LD50_crickets$alive) ~ (M_balfouri_LD50_crickets$venom_conc_percent), 
                                            binomial(link = probit))
                                        
summary(model_M_balfouri_LD50_crickets)


##### plotting the data

#this just generates a sequence of numbers
ypredicts <- seq(0, 1, 0.01)
xpredicts <- dose.p(model_M_balfouri_LD50_crickets,p=c(ypredicts))


plot(M_balfouri_LD50_crickets$dead/20 ~ (M_balfouri_LD50_crickets$venom_conc_percent),
     ylab = "Proportion_dead",
     xlab = "Dosage",
     pch = 16)

lines(unlist(xpredicts), ypredicts)


### LD50 ###
M_balfouri_LD50_crickets_result <- dose.p(model_M_balfouri_LD50_crickets, p=c(0.5))
M_balfouri_LD50_crickets_result
#          Dose     SE
# p = 0.5: 4.124566 0.4507631


### Converting % dose to mg/kg ###

# The calculated dose is the % concentration dose (mg/ul) required to achieve an LD50 in crickets of median mass 0.0003 kg.
# To get LD50 mg/kg, you first figure out how much 100% of the sample equals (0.110545 mg/ul).
# 100% then divided by 100 to get 1% = 0.001105 mg/ul.
# 1% multiplied by % dose = 4.12 = 0.004553 mg/ul.
# 0.004553 mg/ul is then divided by crickets median mass to get mg/kg (0.004553/0.0003 = 15.18 mg/kg).
# Repeat with the standard error dose to get standard range of error in mg/kg (1.66 mg/kg).


### this subsets dataset (woodlice)

M_balfouri_LD50_woodlice <- LD50_data24[LD50_data24$species == "Monocentropus_balfouri" & LD50_data24$prey_species == "Porcellio_scaber", ]


model_M_balfouri_LD50_woodlice <- glm(cbind(M_balfouri_LD50_woodlice$dead, 
                                            M_balfouri_LD50_woodlice$alive) ~ (M_balfouri_LD50_woodlice$venom_conc_percent),
                                            binomial(link = probit))
                                        
summary(model_M_balfouri_LD50_woodlice)


### plotting the data

### this just generates a sequence of numbers
ypredicts <- seq(0, 1, 0.01)
xpredicts <- dose.p(model_M_balfouri_LD50_woodlice,p=c(ypredicts))


plot(M_balfouri_LD50_woodlice$dead/20 ~ (M_balfouri_LD50_woodlice$venom_conc_percent),
     ylab = "Proportion_dead",
     xlab = "Dosage",
     pch = 16)

lines(unlist(xpredicts), ypredicts)


### LD50 ###
M_balfouri_LD50_woodlice_result <- dose.p(model_M_balfouri_LD50_woodlice, p=c(0.5))
M_balfouri_LD50_woodlice_result
#          Dose      SE
# p = 0.5: 0.6890831 0.5269044


### Converting % dose to mg/kg ###

# The calculated dose is the % concentration dose (mg/ul) required to achieve an LD50 in woodlice of median mass 0.00009 kg.
# To get LD50 mg/kg, you first figure out how much 100% of the sample equals (0.110545 mg/ul).
# 100% then divided by 100 to get 1% = 0.001105 mg/ul.
# 1% multiplied by % dose = 0.69 = 0.000762 mg/ul.
# 0.000762 mg/ul is then divided by woodlice median mass to get mg/kg (0.000762/0.00009 = 8.47 mg/kg).
# Repeat with the standard error dose to get standard range of error in mg/kg (6.51 mg/kg).


########################################
##### Let's try with 1hr peak data #####
########################################


#################################
### First, calculate ED50 1hr ###
#################################

### this subsets dataset

M_balfouri_ED50_crickets_1hr <- ED50_data_peaks_1hr[ED50_data_peaks_1hr$species == "Monocentropus_balfouri" & ED50_data_peaks_1hr$prey_species == "Acheta_domesticus", ]

model_M_balfouri_ED50_crickets_1hr <- glm(cbind(M_balfouri_ED50_crickets_1hr$incapacitated_or_dead, 
                                            M_balfouri_ED50_crickets_1hr$alive) ~ (M_balfouri_ED50_crickets_1hr$venom_conc_percent), 
                                            binomial(link = probit))
                                        
summary(model_M_balfouri_ED50_crickets_1hr)


##### plotting the data

#this just generates a sequence of numbers
ypredicts <- seq(0, 1, 0.01)
xpredicts <- dose.p(model_M_balfouri_ED50_crickets_1hr,p=c(ypredicts))


plot(M_balfouri_ED50_crickets_1hr$incapacitated_or_dead/20 ~ (M_balfouri_ED50_crickets_1hr$venom_conc_percent),
     ylab = "Proportion_incap",
     xlab = "Dosage",
     pch = 16)

lines(unlist(xpredicts), ypredicts)


M_balfouri_ED50_crickets_result_1hr <- dose.p(model_M_balfouri_ED50_crickets_1hr, p=c(0.5))
M_balfouri_ED50_crickets_result_1hr
#                  Dose        SE
#    p = 0.5: 0.8050086 0.2500554


### Converting % dose to mg/kg ###

# The calculated dose is the % concentration dose (mg/ul) required to achieve an ED50 in crickets of median mass 0.0003 kg.
# To get ED50 mg/kg, you first figure out how much 100% of the sample equals (0.1105527638190955 mg/ul).
# 100% then divided by 100 to get 1% = 0.001105 mg/ul.
# 1% multiplied by % dose = 0.81 = 0.000895 mg/ul.
# 0.000895 mg/ul is then divided by cricket median mass to get mg/kg (0.000895/0.0003 = 2.98 mg/kg).
# Repeat with the standard error dose to get standard range of error in mg/kg (0.92 mg/kg).


### this subsets dataset (woodlice)

M_balfouri_ED50_woodlice_1hr <- ED50_data_peaks_1hr[ED50_data_peaks_1hr$species == "Monocentropus_balfouri" & ED50_data_peaks_1hr$prey_species == "Porcellio_scaber", ]


model_M_balfouri_ED50_woodlice_1hr <- glm(cbind(M_balfouri_ED50_woodlice_1hr$incapacitated_or_dead, 
                                            M_balfouri_ED50_woodlice_1hr$alive) ~ (M_balfouri_ED50_woodlice_1hr$venom_conc_percent),
                                            binomial(link = probit))
                                     
summary(model_M_balfouri_ED50_woodlice_1hr)


### plotting the data

### this just generates a sequence of numbers
ypredicts <- seq(0, 1, 0.01)
xpredicts <- dose.p(model_M_balfouri_ED50_woodlice_1hr,p=c(ypredicts))


plot(M_balfouri_ED50_woodlice_1hr$incapacitated_or_dead/20 ~ (M_balfouri_ED50_woodlice_1hr$venom_conc_percent),
     ylab = "Proportion_incap",
     xlab = "Dosage",
     pch = 16)

lines(unlist(xpredicts), ypredicts)


M_balfouri_ED50_woodlice_result_1hr <- dose.p(model_M_balfouri_ED50_woodlice_1hr, p=c(0.5))
M_balfouri_ED50_woodlice_result_1hr
#             Dose      SE
#    p = 0.5: 0.9826991 0.4552866


### Converting % dose to mg/kg ###

# The calculated dose is the % concentration dose (mg/ul) required to achieve an ED50 in woodlice of median mass 0.00009 kg.
# To get ED50 mg/kg, you first figure out how much 100% of the sample equals (0.1105527638190955 mg/ul).
# 100% then divided by 100 to get 1% = 0.001105 mg/ul.
# 1% multiplied by % dose = 0.98 = 0.0010829 mg/ul.
# 0.0010829 mg/ul is then divided by woodlice median mass to get mg/kg (0.0010829/0.00009 = 12.03 mg/kg).
# Repeat with the standard error dose to get standard range of error in mg/kg (5.59 mg/kg).

```


```{r Pholcus phalangioides ED50 and LD50 }

#############################
### First, calculate ED50 ###
#############################

### this subsets dataset

P_phalangioides_ED50_crickets <- ED50_data_peaks[ED50_data_peaks$species == "Pholcus_phalangioides" & ED50_data_peaks$prey_species == "Acheta_domesticus", ]

model_P_phalangioides_ED50_crickets <- glm(cbind(P_phalangioides_ED50_crickets$incapacitated_or_dead, 
                                                 P_phalangioides_ED50_crickets$alive) ~ (P_phalangioides_ED50_crickets$venom_conc_percent), 
                                                 binomial(link = probit))
                                             
summary(model_P_phalangioides_ED50_crickets)


### plotting the data

### this just generates a sequence of numbers
ypredicts <- seq(0, 1, 0.01)
xpredicts <- dose.p(model_P_phalangioides_ED50_crickets,p=c(ypredicts))


plot(P_phalangioides_ED50_crickets$incapacitated_or_dead/20 ~ (P_phalangioides_ED50_crickets$venom_conc_percent),
     ylab = "Proportion_incap",
     xlab = "Dosage",
     pch = 16)

lines(unlist(xpredicts), ypredicts)


P_phalangioides_ED50_crickets_result <- dose.p(model_P_phalangioides_ED50_crickets, p=c(0.5))
P_phalangioides_ED50_crickets_result
#                  Dose       SE
#   p = 0.5: 0.2910002 0.1542442


### Converting % dose to mg/kg ###

# The calculated dose is the % concentration dose (mg/ul) required to achieve an ED50 in crickets of median mass 0.0003 kg.
# To get ED50 mg/kg, you first figure out how much 100% of the sample equals (0.0822784810126582 mg/ul).
# 100% then divided by 100 to get 1% = 0.000823 mg/ul.
# 1% multiplied by % dose = 0.29 = 0.000239 mg/ul.
# 0.000239 mg/ul is then divided by cricket median mass to get mg/kg (0.000239/0.0003 = 0.80 mg/kg).
# Repeat with the standard error dose to get standard range of error in mg/kg (0.41 mg/kg).


### this subsets dataset (woodlice)

P_phalangioides_ED50_woodlice <- ED50_data_peaks[ED50_data_peaks$species == "Pholcus_phalangioides" & ED50_data_peaks$prey_species == "Porcellio_scaber", ]


model_P_phalangioides_ED50_woodlice <- glm(cbind(P_phalangioides_ED50_woodlice$incapacitated_or_dead, 
                                                 P_phalangioides_ED50_woodlice$alive) ~ (P_phalangioides_ED50_woodlice$venom_conc_percent),
                                                 binomial(link = probit))
                                             
summary(model_P_phalangioides_ED50_woodlice)


### plotting the data

### this just generates a sequence of numbers
ypredicts <- seq(0, 1, 0.01)
xpredicts <- dose.p(model_P_phalangioides_ED50_woodlice,p=c(ypredicts))


plot(P_phalangioides_ED50_woodlice$incapacitated_or_dead/20 ~ (P_phalangioides_ED50_woodlice$venom_conc_percent),
     ylab = "Proportion_incap",
     xlab = "Dosage",
     pch = 16)

lines(unlist(xpredicts), ypredicts)


P_phalangioides_ED50_woodlice_result <- dose.p(model_P_phalangioides_ED50_woodlice, p=c(0.5))
P_phalangioides_ED50_woodlice_result
#                 Dose        SE
#  p = 0.5:  0.4888637 0.1799947


### Converting % dose to mg/kg ###

# The calculated dose is the % concentration dose (mg/ul) required to achieve an ED50 in woodlice of median mass 0.00009 kg.
# To get ED50 mg/kg, you first figure out how much 100% of the sample equals (0.0822784810126582 mg/ul).
# 100% then divided by 100 to get 1% = 0.000823 mg/ul.
# 1% multiplied by % dose = 0.49 = 0.000403 mg/ul.
# 0.000403 mg/ul is then divided by woodlice median mass to get mg/kg (0.000403/0.00009 = 4.48 mg/kg).
# Repeat with the standard error dose to get standard range of error in mg/kg (1.64 mg/kg).


##########################
### Now calculate LD50 ###
##########################


### this subsets dataset

P_phalangioides_LD50_crickets <- LD50_data24[LD50_data24$species == "Pholcus_phalangioides" & LD50_data24$prey_species == "Acheta_domesticus", ]

model_P_phalangioides_LD50_crickets <- glm(cbind(P_phalangioides_LD50_crickets$dead, 
                                                 P_phalangioides_LD50_crickets$alive) ~ (P_phalangioides_LD50_crickets$venom_conc_percent), 
                                                 binomial(link = probit))
                                             
summary(model_P_phalangioides_LD50_crickets)


### plotting the data

### this just generates a sequence of numbers
ypredicts <- seq(0, 1, 0.01)
xpredicts <- dose.p(model_P_phalangioides_LD50_crickets,p=c(ypredicts))


plot(P_phalangioides_LD50_crickets$dead/20 ~ (P_phalangioides_LD50_crickets$venom_conc_percent),
     ylab = "Proportion_dead",
     xlab = "Dosage",
     pch = 16)

lines(unlist(xpredicts), ypredicts)


### LD50 ###
P_phalangioides_LD50_crickets_result <- dose.p(model_P_phalangioides_LD50_crickets, p=c(0.5))
P_phalangioides_LD50_crickets_result
#          Dose     SE
# p = 0.5: 2.302703 0.8170344


### Converting % dose to mg/kg ###

# The calculated dose is the % concentration dose (mg/ul) required to achieve an LD50 in crickets of median mass 0.0003 kg.
# To get LD50 mg/kg, you first figure out how much 100% of the sample equals (0.082278 mg/ul).
# 100% then divided by 100 to get 1% = 0.000823 mg/ul.
# 1% multiplied by % dose = 2.30 = 0.001893 mg/ul.
# 0.001893 mg/ul is then divided by cricket median mass to get mg/kg (0.001893/0.0003 = 6.31 mg/kg).
# Repeat with the standard error dose to get standard range of error in mg/kg (2.25 mg/kg).


### this subsets dataset (woodlice)

P_phalangioides_LD50_woodlice <- LD50_data24[LD50_data24$species == "Pholcus_phalangioides" & LD50_data24$prey_species == "Porcellio_scaber", ]


model_P_phalangioides_LD50_woodlice <- glm(cbind(P_phalangioides_LD50_woodlice$dead, 
                                                 P_phalangioides_LD50_woodlice$alive) ~ (P_phalangioides_LD50_woodlice$venom_conc_percent),
                                                 binomial(link = probit))
                                             
summary(model_P_phalangioides_LD50_woodlice)


### plotting the data

### this just generates a sequence of numbers
ypredicts <- seq(0, 1, 0.01)
xpredicts <- dose.p(model_P_phalangioides_LD50_woodlice,p=c(ypredicts))


plot(P_phalangioides_LD50_woodlice$dead/20 ~ (P_phalangioides_LD50_woodlice$venom_conc_percent),
     ylab = "Proportion_dead",
     xlab = "Dosage",
     pch = 16)

lines(unlist(xpredicts), ypredicts)


### LD50 ###
P_phalangioides_LD50_woodlice_result <- dose.p(model_P_phalangioides_LD50_woodlice, p=c(0.5))
P_phalangioides_LD50_woodlice_result
#          Dose    SE
# p = 0.5: 1.37646 0.172158


### Converting % dose to mg/kg ###

# The calculated dose is the % concentration dose (mg/ul) required to achieve an LD50 in woodlice of median mass 0.00009 kg.
# To get LD50 mg/kg, you first figure out how much 100% of the sample equals (0.082278 mg/ul).
# 100% then divided by 100 to get 1% = 0.000823 mg/ul.
# 1% multiplied by % dose = 1.38 = 0.001136 mg/ul.
# 0.001136 mg/ul is then divided by woodlice median mass to get mg/kg (0.001136/0.00009 = 12.62 mg/kg).
# Repeat with the standard error dose to get standard range of error in mg/kg (1.56 mg/kg).


########################################
##### Let's try with 1hr peak data #####
########################################


#############################
### First, calculate ED50 ###
#############################

### this subsets dataset

P_phalangioides_ED50_crickets_1hr <- ED50_data_peaks_1hr[ED50_data_peaks_1hr$species == "Pholcus_phalangioides" & ED50_data_peaks_1hr$prey_species == "Acheta_domesticus", ]

model_P_phalangioides_ED50_crickets_1hr <- glm(cbind(P_phalangioides_ED50_crickets_1hr$incapacitated_or_dead, 
                                                 P_phalangioides_ED50_crickets_1hr$alive) ~ (P_phalangioides_ED50_crickets_1hr$venom_conc_percent), 
                                                 binomial(link = probit))
                                             
summary(model_P_phalangioides_ED50_crickets_1hr)


### plotting the data

### this just generates a sequence of numbers
ypredicts <- seq(0, 1, 0.01)
xpredicts <- dose.p(model_P_phalangioides_ED50_crickets_1hr,p=c(ypredicts))


plot(P_phalangioides_ED50_crickets_1hr$incapacitated_or_dead/20 ~ (P_phalangioides_ED50_crickets_1hr$venom_conc_percent),
     ylab = "Proportion_incap",
     xlab = "Dosage",
     pch = 16)

lines(unlist(xpredicts), ypredicts)


P_phalangioides_ED50_crickets_result_1hr <- dose.p(model_P_phalangioides_ED50_crickets_1hr, p=c(0.5))
P_phalangioides_ED50_crickets_result_1hr
#            Dose    SE
#   p = 0.5: 0.77718 0.1614755


### Converting % dose to mg/kg ###

# The calculated dose is the % concentration dose (mg/ul) required to achieve an ED50 in crickets of median mass 0.0003 kg.
# To get ED50 mg/kg, you first figure out how much 100% of the sample equals (0.0822784810126582 mg/ul).
# 100% then divided by 100 to get 1% = 0.000823 mg/ul.
# 1% multiplied by % dose = 0.78 = 0.0006419 mg/ul.
# 0.0006419 mg/ul is then divided by cricket median mass to get mg/kg (0.0006419/0.0003 = 2.14 mg/kg).
# Repeat with the standard error dose to get standard range of error in mg/kg (0.44 mg/kg).


### this subsets dataset (woodlice)

P_phalangioides_ED50_woodlice_1hr <- ED50_data_peaks_1hr[ED50_data_peaks_1hr$species == "Pholcus_phalangioides" & ED50_data_peaks_1hr$prey_species == "Porcellio_scaber", ]


model_P_phalangioides_ED50_woodlice_1hr <- glm(cbind(P_phalangioides_ED50_woodlice_1hr$incapacitated_or_dead, 
                                                 P_phalangioides_ED50_woodlice_1hr$alive) ~ (P_phalangioides_ED50_woodlice_1hr$venom_conc_percent),
                                                 binomial(link = probit))
                                             
summary(model_P_phalangioides_ED50_woodlice_1hr)


### plotting the data

### this just generates a sequence of numbers
ypredicts <- seq(0, 1, 0.01)
xpredicts <- dose.p(model_P_phalangioides_ED50_woodlice_1hr,p=c(ypredicts))


plot(P_phalangioides_ED50_woodlice_1hr$incapacitated_or_dead/20 ~ (P_phalangioides_ED50_woodlice_1hr$venom_conc_percent),
     ylab = "Proportion_incap",
     xlab = "Dosage",
     pch = 16)

lines(unlist(xpredicts), ypredicts)


P_phalangioides_ED50_woodlice_result_1hr <- dose.p(model_P_phalangioides_ED50_woodlice_1hr, p=c(0.5))
P_phalangioides_ED50_woodlice_result_1hr
#            Dose      SE
#  p = 0.5:  0.9361665 0.1402735


### Converting % dose to mg/kg ###

# The calculated dose is the % concentration dose (mg/ul) required to achieve an ED50 in woodlice of median mass 0.00009 kg.
# To get ED50 mg/kg, you first figure out how much 100% of the sample equals (0.0822784810126582 mg/ul).
# 100% then divided by 100 to get 1% = 0.000823 mg/ul.
# 1% multiplied by % dose = 0.94 = 0.0007736 mg/ul.
# 0.0007736 mg/ul is then divided by woodlice median mass to get mg/kg (0.0007736/0.00009 = 8.59 mg/kg).
# Repeat with the standard error dose to get standard range of error in mg/kg (1.28 mg/kg).


```


```{r Phormictopus cancerides ED50 and LD50}

#############################
### First, calculate ED50 ###
#############################

### this subsets dataset

P_cancerides_ED50_crickets <- ED50_data_peaks[ED50_data_peaks$species == "Phormictopus_cancerides" & ED50_data_peaks$prey_species == "Acheta_domesticus", ]

model_P_cancerides_ED50_crickets <- glm(cbind(P_cancerides_ED50_crickets$incapacitated_or_dead, 
                                              P_cancerides_ED50_crickets$alive) ~ (P_cancerides_ED50_crickets$venom_conc_percent), 
                                              binomial(link = probit))
                                          
summary(model_P_cancerides_ED50_crickets)


### plotting the data

### this just generates a sequence of numbers
ypredicts <- seq(0, 1, 0.01)
xpredicts <- dose.p(model_P_cancerides_ED50_crickets,p=c(ypredicts))


plot(P_cancerides_ED50_crickets$incapacitated_or_dead/20 ~ (P_cancerides_ED50_crickets$venom_conc_percent),
     ylab = "Proportion_incap",
     xlab = "Dosage",
     pch = 16)

lines(unlist(xpredicts), ypredicts)


P_cancerides_ED50_crickets_result <- dose.p(model_P_cancerides_ED50_crickets, p=c(0.5))
P_cancerides_ED50_crickets_result
#                Dose       SE
#   p = 0.5: 2.720987 0.5766899


### Converting % dose to mg/kg ###

# The calculated dose is the % concentration dose (mg/ul) required to achieve an ED50 in crickets of median mass 0.0003 kg.
# To get ED50 mg/kg, you first figure out how much 100% of the sample equals (0.1842900302114804 mg/ul).
# 100% then divided by 100 to get 1% = 0.001843 mg/ul.
# 1% multiplied by % dose = 2.72 = 0.005013 mg/ul.
# 0.005013 mg/ul is then divided by cricket median mass to get mg/kg (0.005013/0.0003 = 16.71 mg/kg).
# Repeat with the standard error dose to get standard range of error in mg/kg (3.56 mg/kg).


### this subsets dataset (woodlice)

P_cancerides_ED50_woodlice <- ED50_data_peaks[ED50_data_peaks$species == "Phormictopus_cancerides" & ED50_data_peaks$prey_species == "Porcellio_scaber", ]


model_P_cancerides_ED50_woodlice <- glm(cbind(P_cancerides_ED50_woodlice$incapacitated_or_dead, 
                                              P_cancerides_ED50_woodlice$alive) ~ (P_cancerides_ED50_woodlice$venom_conc_percent),
                                              binomial(link = probit))
                                          
summary(model_P_cancerides_ED50_woodlice)


### plotting the data

### this just generates a sequence of numbers
ypredicts <- seq(0, 1, 0.01)
xpredicts <- dose.p(model_P_cancerides_ED50_woodlice,p=c(ypredicts))


plot(P_cancerides_ED50_woodlice$incapacitated_or_dead/20 ~ (P_cancerides_ED50_woodlice$venom_conc_percent),
     ylab = "Proportion_incap",
     xlab = "Dosage",
     pch = 16)

lines(unlist(xpredicts), ypredicts)


P_cancerides_ED50_woodlice_result <- dose.p(model_P_cancerides_ED50_woodlice, p=c(0.5))
P_cancerides_ED50_woodlice_result
#                Dose       SE
#  p = 0.5: 1.373494 0.1979401


### Converting % dose to mg/kg ###

# The calculated dose is the % concentration dose (mg/ul) required to achieve an ED50 in woodlice of median mass 0.00009 kg.
# To get ED50 mg/kg, you first figure out how much 100% of the sample equals (0.1842900302114804 mg/ul).
# 100% then divided by 100 to get 1% = 0.001843 mg/ul.
# 1% multiplied by % dose = 1.37 = 0.002525 mg/ul.
# 0.002525 mg/ul is then divided by woodlice median mass to get mg/kg (0.002525/0.00009 = 28.06 mg/kg).
# Repeat with the standard error dose to get standard range of error in mg/kg (3.87 mg/kg).


##########################
### Now calculate LD50 ###
##########################

### this subsets dataset

P_cancerides_LD50_crickets <- LD50_data24[LD50_data24$species == "Phormictopus_cancerides" & LD50_data24$prey_species == "Acheta_domesticus", ]

model_P_cancerides_LD50_crickets <- glm(cbind(P_cancerides_LD50_crickets$dead, 
                                              P_cancerides_LD50_crickets$alive) ~ (P_cancerides_LD50_crickets$venom_conc_percent), 
                                              binomial(link = probit))
                                          
summary(model_P_cancerides_LD50_crickets)


### plotting the data

### this just generates a sequence of numbers
ypredicts <- seq(0, 1, 0.01)
xpredicts <- dose.p(model_P_cancerides_LD50_crickets,p=c(ypredicts))


plot(P_cancerides_LD50_crickets$dead/20 ~ (P_cancerides_LD50_crickets$venom_conc_percent),
     ylab = "Proportion_dead",
     xlab = "Dosage",
     pch = 16)

lines(unlist(xpredicts), ypredicts)


### LD50 ###
P_cancerides_LD50_crickets_result <- dose.p(model_P_cancerides_LD50_crickets, p=c(0.5))
P_cancerides_LD50_crickets_result
#          Dose     SE
# p = 0.5: 9.575621 8.78483


### Converting % dose to mg/kg ###

# The calculated dose is the % concentration dose (mg/ul) required to achieve an LD50 in crickets of median mass 0.0003 kg.
# To get LD50 mg/kg, you first figure out how much 100% of the sample equals (0.1842900302114804 mg/ul).
# 100% then divided by 100 to get 1% = 0.001843 mg/ul.
# 1% multiplied by % dose = 9.58 = 0.017656 mg/ul.
# 0.017656 mg/ul is then divided by cricket median mass to get mg/kg (0.017656/0.0003 = 58.85 mg/kg).
# Repeat with the standard error dose to get standard range of error in mg/kg (53.94 mg/kg).


### this subsets dataset (woodlice)

P_cancerides_LD50_woodlice <- LD50_data24[LD50_data24$species == "Phormictopus_cancerides" & LD50_data24$prey_species == "Porcellio_scaber", ]


model_P_cancerides_LD50_woodlice <- glm(cbind(P_cancerides_LD50_woodlice$dead, 
                                              P_cancerides_LD50_woodlice$alive) ~ (P_cancerides_LD50_woodlice$venom_conc_percent),
                                              binomial(link = probit))
                                          
summary(model_P_cancerides_LD50_woodlice)


### plotting the data

### this just generates a sequence of numbers
ypredicts <- seq(0, 1, 0.01)
xpredicts <- dose.p(model_P_cancerides_LD50_woodlice,p=c(ypredicts))


plot(P_cancerides_LD50_woodlice$dead/20 ~ (P_cancerides_LD50_woodlice$venom_conc_percent),
     ylab = "Proportion_dead",
     xlab = "Dosage",
     pch = 16)

lines(unlist(xpredicts), ypredicts)


### LD50 ###
P_cancerides_LD50_woodlice_result <- dose.p(model_P_cancerides_LD50_woodlice, p=c(0.5))
P_cancerides_LD50_woodlice_result
#          Dose     SE
# p = 0.5: 4.895905 4.814836


### Converting % dose to mg/kg ###

# The calculated dose is the % concentration dose (mg/ul) required to achieve an LD50 in woodlice of median mass 0.00009 kg.
# To get LD50 mg/kg, you first figure out how much 100% of the sample equals (0.1842900302114804 mg/ul).
# 100% then divided by 100 to get 1% = 0.001843 mg/ul.
# 1% multiplied by % dose = 4.9 = 0.009307 mg/ul.
# 0.009307 mg/ul is then divided by woodlice median mass to get mg/kg (0.009307/0.00009 = 100.34 mg/kg).
# Repeat with the standard error dose to get standard range of error in mg/kg (98.5 mg/kg).


########################################
##### Let's try with 1hr peak data #####
########################################


#################################
### First, calculate ED50 1hr ###
#################################

### this subsets dataset

P_cancerides_ED50_crickets_1hr <- ED50_data_peaks_1hr[ED50_data_peaks_1hr$species == "Phormictopus_cancerides" & ED50_data_peaks_1hr$prey_species == "Acheta_domesticus", ]

model_P_cancerides_ED50_crickets_1hr <- glm(cbind(P_cancerides_ED50_crickets_1hr$incapacitated_or_dead, 
                                              P_cancerides_ED50_crickets_1hr$alive) ~ (P_cancerides_ED50_crickets_1hr$venom_conc_percent), 
                                              binomial(link = probit))
                                          
summary(model_P_cancerides_ED50_crickets_1hr)


### plotting the data

### this just generates a sequence of numbers
ypredicts <- seq(0, 1, 0.01)
xpredicts <- dose.p(model_P_cancerides_ED50_crickets_1hr,p=c(ypredicts))


plot(P_cancerides_ED50_crickets_1hr$incapacitated_or_dead/20 ~ (P_cancerides_ED50_crickets_1hr$venom_conc_percent),
     ylab = "Proportion_incap",
     xlab = "Dosage",
     pch = 16)

lines(unlist(xpredicts), ypredicts)


P_cancerides_ED50_crickets_result_1hr <- dose.p(model_P_cancerides_ED50_crickets_1hr, p=c(0.5))
P_cancerides_ED50_crickets_result_1hr
#                Dose       SE
#   p = 0.5: 2.720987 0.5766899


### Converting % dose to mg/kg ###

# The calculated dose is the % concentration dose (mg/ul) required to achieve an ED50 in crickets of median mass 0.0003 kg.
# To get ED50 mg/kg, you first figure out how much 100% of the sample equals (0.1842900302114804 mg/ul).
# 100% then divided by 100 to get 1% = 0.001843 mg/ul.
# 1% multiplied by % dose = 2.72 = 0.005013 mg/ul.
# 0.005013 mg/ul is then divided by cricket median mass to get mg/kg (0.005013/0.0003 = 16.71 mg/kg).
# Repeat with the standard error dose to get standard range of error in mg/kg (3.56 mg/kg).


### this subsets dataset (woodlice)

P_cancerides_ED50_woodlice_1hr <- ED50_data_peaks_1hr[ED50_data_peaks_1hr$species == "Phormictopus_cancerides" & ED50_data_peaks_1hr$prey_species == "Porcellio_scaber", ]


model_P_cancerides_ED50_woodlice_1hr <- glm(cbind(P_cancerides_ED50_woodlice_1hr$incapacitated_or_dead, 
                                              P_cancerides_ED50_woodlice_1hr$alive) ~ (P_cancerides_ED50_woodlice_1hr$venom_conc_percent),
                                              binomial(link = probit))
                                          
summary(model_P_cancerides_ED50_woodlice_1hr)


### plotting the data

### this just generates a sequence of numbers
ypredicts <- seq(0, 1, 0.01)
xpredicts <- dose.p(model_P_cancerides_ED50_woodlice_1hr,p=c(ypredicts))


plot(P_cancerides_ED50_woodlice_1hr$incapacitated_or_dead/20 ~ (P_cancerides_ED50_woodlice_1hr$venom_conc_percent),
     ylab = "Proportion_incap",
     xlab = "Dosage",
     pch = 16)

lines(unlist(xpredicts), ypredicts)


P_cancerides_ED50_woodlice_result_1hr <- dose.p(model_P_cancerides_ED50_woodlice_1hr, p=c(0.5))
P_cancerides_ED50_woodlice_result_1hr
#           Dose    SE
#  p = 0.5: 2.59003 0.4099108


### Converting % dose to mg/kg ###

# The calculated dose is the % concentration dose (mg/ul) required to achieve an ED50 in woodlice of median mass 0.00009 kg.
# To get ED50 mg/kg, you first figure out how much 100% of the sample equals (0.1842900302114804 mg/ul).
# 100% then divided by 100 to get 1% = 0.001843 mg/ul.
# 1% multiplied by % dose = 2.59 = 0.00477 mg/ul.
# 0.00477 mg/ul is then divided by woodlice median mass to get mg/kg (0.00477/0.00009 = 53.0 mg/kg).
# Repeat with the standard error dose to get standard range of error in mg/kg (8.39 mg/kg).


```


```{r Piloctenus haematostoma ED50 and LD50}

#############################
### First, calculate ED50 ###
#############################

### this subsets dataset

P_haematostoma_ED50_crickets <- ED50_data_peaks[ED50_data_peaks$species == "Piloctenus_haematostoma" & ED50_data_peaks$prey_species == "Acheta_domesticus",]

model_P_haematostoma_ED50_crickets <- glm(cbind(P_haematostoma_ED50_crickets$incapacitated_or_dead, 
                                                P_haematostoma_ED50_crickets$alive) ~ (P_haematostoma_ED50_crickets$venom_conc_percent), 
                                                binomial(link = probit))
                                            
summary(model_P_haematostoma_ED50_crickets)


### plotting the data

### this just generates a sequence of numbers
ypredicts <- seq(0, 1, 0.01)
xpredicts <- dose.p(model_P_haematostoma_ED50_crickets,p=c(ypredicts))


plot(P_haematostoma_ED50_crickets$incapacitated_or_dead/20 ~ (P_haematostoma_ED50_crickets$venom_conc_percent),
     ylab = "Proportion_incap",
     xlab = "Dosage",
     pch = 16)

lines(unlist(xpredicts), ypredicts)


P_haematostoma_ED50_crickets_result <- dose.p(model_P_haematostoma_ED50_crickets, p=c(0.5))
P_haematostoma_ED50_crickets_result
#              Dose       SE
#  p = 0.5: 8.06229 1.741125


### Converting % dose to mg/kg ###

# The calculated dose is the % concentration dose (mg/ul) required to achieve an ED50 in crickets of median mass 0.0003 kg.
# To get ED50 mg/kg, you first figure out how much 100% of the sample equals (0.0099042588312975 mg/ul).
# 100% then divided by 100 to get 1% = 0.000099 mg/ul.
# 1% multiplied by % dose = 8.06 = 0.000798 mg/ul.
# 0.000798 mg/ul is then divided by cricket median mass to get mg/kg (0.000798/0.0003 = 2.66 mg/kg).
# Repeat with the standard error dose to get standard range of error in mg/kg (0.57 mg/kg).


### this subsets dataset (woodlice)

P_haematostoma_ED50_woodlice <- ED50_data_peaks[ED50_data_peaks$species == "Piloctenus_haematostoma" & ED50_data_peaks$prey_species == "Porcellio_scaber", ]


model_P_haematostoma_ED50_woodlice <- glm(cbind(P_haematostoma_ED50_woodlice$incapacitated_or_dead, 
                                                P_haematostoma_ED50_woodlice$alive) ~ (P_haematostoma_ED50_woodlice$venom_conc_percent),
                                                binomial(link = probit))
                                            
summary(model_P_haematostoma_ED50_woodlice)


### plotting the data

### this just generates a sequence of numbers
ypredicts <- seq(0, 1, 0.01)
xpredicts <- dose.p(model_P_haematostoma_ED50_woodlice,p=c(ypredicts))


plot(P_haematostoma_ED50_woodlice$incapacitated_or_dead/20 ~ (P_haematostoma_ED50_woodlice$venom_conc_percent),
     ylab = "Proportion_incap",
     xlab = "Dosage",
     pch = 16)

lines(unlist(xpredicts), ypredicts)


P_haematostoma_ED50_woodlice_result <- dose.p(model_P_haematostoma_ED50_woodlice, p=c(0.5))
P_haematostoma_ED50_woodlice_result
#                      Dose       SE
#         p = 0.5: 9.376282 2.856322


### Converting % dose to mg/kg ###

# The calculated dose is the % concentration dose (mg/ul) required to achieve an ED50 in woodlice of median mass 0.00009 kg.
# To get ED50 mg/kg, you first figure out how much 100% of the sample equals (0.0099042588312975 mg/ul).
# 100% then divided by 100 to get 1% = 0.000099 mg/ul.
# 1% multiplied by % dose = 9.38 = 0.000929 mg/ul.
# 0.000929 mg/ul is then divided by woodlice median mass to get mg/kg (0.000929/0.00009 = 10.32 mg/kg).
# Repeat with the standard error dose to get standard range of error in mg/kg (3.14 mg/kg).


##########################
### Now calculate LD50 ###
##########################

### this subsets dataset

P_haematostoma_LD50_crickets <- LD50_data24[LD50_data24$species == "Piloctenus_haematostoma" & LD50_data24$prey_species == "Acheta_domesticus",]

model_P_haematostoma_LD50_crickets <- glm(cbind(P_haematostoma_LD50_crickets$dead, 
                                                P_haematostoma_LD50_crickets$alive) ~ (P_haematostoma_LD50_crickets$venom_conc_percent), 
                                                binomial(link = probit))
                                            
summary(model_P_haematostoma_LD50_crickets)


### plotting the data

### this just generates a sequence of numbers
ypredicts <- seq(0, 1, 0.01)
xpredicts <- dose.p(model_P_haematostoma_LD50_crickets,p=c(ypredicts))


plot(P_haematostoma_LD50_crickets$dead/20 ~ (P_haematostoma_LD50_crickets$venom_conc_percent),
     ylab = "Proportion_dead",
     xlab = "Dosage",
     pch = 16)

lines(unlist(xpredicts), ypredicts)


### LD50 ###
P_haematostoma_LD50_crickets_result <- dose.p(model_P_haematostoma_LD50_crickets, p=c(0.5))
P_haematostoma_LD50_crickets_result
#          Dose     SE
# p = 0.5: 17.81783 5.70640


### Converting % dose to mg/kg ###

# The calculated dose is the % concentration dose (mg/ul) required to achieve an LD50 in crickets of median mass 0.0003 kg.
# To get LD50 mg/kg, you first figure out how much 100% of the sample equals (0.0099042588312975 mg/ul).
# 100% then divided by 100 to get 1% = 0.000099 mg/ul.
# 1% multiplied by % dose = 17.82 = 0.001764 mg/ul.
# 0.001764 mg/ul is then divided by cricket median mass to get mg/kg (0.001764/0.0003 = 5.88 mg/kg).
# Repeat with the standard error dose to get standard range of error in mg/kg (1.88 mg/kg).


### this subsets dataset (woodlice)

P_haematostoma_LD50_woodlice <- LD50_data24[LD50_data24$species == "Piloctenus_haematostoma" & LD50_data24$prey_species == "Porcellio_scaber", ]


model_P_haematostoma_LD50_woodlice <- glm(cbind(P_haematostoma_LD50_woodlice$dead, 
                                                P_haematostoma_LD50_woodlice$alive) ~ (P_haematostoma_LD50_woodlice$venom_conc_percent),
                                                binomial(link = probit))
                                            
summary(model_P_haematostoma_LD50_woodlice)


### plotting the data

### this just generates a sequence of numbers
ypredicts <- seq(0, 1, 0.01)
xpredicts <- dose.p(model_P_haematostoma_LD50_woodlice,p=c(ypredicts))


plot(P_haematostoma_LD50_woodlice$dead/20 ~ (P_haematostoma_LD50_woodlice$venom_conc_percent),
     ylab = "Proportion_dead",
     xlab = "Dosage",
     pch = 16)

lines(unlist(xpredicts), ypredicts)


### LD50 ###
P_haematostoma_LD50_woodlice_result <- dose.p(model_P_haematostoma_LD50_woodlice, p=c(0.5))
P_haematostoma_LD50_woodlice_result
#          Dose     SE
# p = 0.5: 17.46154 8.614369


### Converting % dose to mg/kg ###

# The calculated dose is the % concentration dose (mg/ul) required to achieve an LD50 in woodlice of median mass 0.00009 kg.
# To get LD50 mg/kg, you first figure out how much 100% of the sample equals (0.0099042588312975 mg/ul).
# 100% then divided by 100 to get 1% = 0.000099 mg/ul.
# 1% multiplied by % dose = 17.46 = 0.001729 mg/ul.
# 0.001729 mg/ul is then divided by woodlice median mass to get mg/kg (0.001729/0.00009 = 19.21 mg/kg).
# Repeat with the standard error dose to get standard range of error in mg/kg (9.47 mg/kg).


########################################
##### Let's try with 1hr peak data #####
########################################


#################################
### First, calculate ED50 1hr ###
#################################

### this subsets dataset

P_haematostoma_ED50_crickets_1hr <- ED50_data_peaks_1hr[ED50_data_peaks_1hr$species == "Piloctenus_haematostoma" & ED50_data_peaks_1hr$prey_species == "Acheta_domesticus",]

model_P_haematostoma_ED50_crickets_1hr <- glm(cbind(P_haematostoma_ED50_crickets_1hr$incapacitated_or_dead, 
                                                P_haematostoma_ED50_crickets_1hr$alive) ~ (P_haematostoma_ED50_crickets_1hr$venom_conc_percent), 
                                                binomial(link = probit))
                                            
summary(model_P_haematostoma_ED50_crickets_1hr)


### plotting the data

### this just generates a sequence of numbers
ypredicts <- seq(0, 1, 0.01)
xpredicts <- dose.p(model_P_haematostoma_ED50_crickets_1hr,p=c(ypredicts))


plot(P_haematostoma_ED50_crickets_1hr$incapacitated_or_dead/20 ~ (P_haematostoma_ED50_crickets_1hr$venom_conc_percent),
     ylab = "Proportion_incap",
     xlab = "Dosage",
     pch = 16)

lines(unlist(xpredicts), ypredicts)


P_haematostoma_ED50_crickets_result_1hr <- dose.p(model_P_haematostoma_ED50_crickets_1hr, p=c(0.5))
P_haematostoma_ED50_crickets_result_1hr
#              Dose       SE
#  p = 0.5: 8.06229 1.741125


### Converting % dose to mg/kg ###

# The calculated dose is the % concentration dose (mg/ul) required to achieve an ED50 in crickets of median mass 0.0003 kg.
# To get ED50 mg/kg, you first figure out how much 100% of the sample equals (0.0099042588312975 mg/ul).
# 100% then divided by 100 to get 1% = 0.000099 mg/ul.
# 1% multiplied by % dose = 8.06 = 0.000798 mg/ul.
# 0.000798 mg/ul is then divided by cricket median mass to get mg/kg (0.000798/0.0003 = 2.66 mg/kg).
# Repeat with the standard error dose to get standard range of error in mg/kg (0.57 mg/kg).


### this subsets dataset (woodlice)

P_haematostoma_ED50_woodlice_1hr <- ED50_data_peaks_1hr[ED50_data_peaks_1hr$species == "Piloctenus_haematostoma" & ED50_data_peaks_1hr$prey_species == "Porcellio_scaber", ]


model_P_haematostoma_ED50_woodlice_1hr <- glm(cbind(P_haematostoma_ED50_woodlice_1hr$incapacitated_or_dead, 
                                                P_haematostoma_ED50_woodlice_1hr$alive) ~ (P_haematostoma_ED50_woodlice_1hr$venom_conc_percent),
                                                binomial(link = probit))
                                            
summary(model_P_haematostoma_ED50_woodlice_1hr)


### plotting the data

### this just generates a sequence of numbers
ypredicts <- seq(0, 1, 0.01)
xpredicts <- dose.p(model_P_haematostoma_ED50_woodlice_1hr,p=c(ypredicts))


plot(P_haematostoma_ED50_woodlice_1hr$incapacitated_or_dead/20 ~ (P_haematostoma_ED50_woodlice_1hr$venom_conc_percent),
     ylab = "Proportion_incap",
     xlab = "Dosage",
     pch = 16)

lines(unlist(xpredicts), ypredicts)


P_haematostoma_ED50_woodlice_result_1hr <- dose.p(model_P_haematostoma_ED50_woodlice_1hr, p=c(0.5))
P_haematostoma_ED50_woodlice_result_1hr
#                  Dose     SE
#         p = 0.5: 23.06114 17.11145


### Converting % dose to mg/kg ###

# The calculated dose is the % concentration dose (mg/ul) required to achieve an ED50 in woodlice of median mass 0.00009 kg.
# To get ED50 mg/kg, you first figure out how much 100% of the sample equals (0.0099042588312975 mg/ul).
# 100% then divided by 100 to get 1% = 0.000099 mg/ul.
# 1% multiplied by % dose = 23.06 = 0.002283 mg/ul.
# 0.002283 mg/ul is then divided by woodlice median mass to get mg/kg (0.002283/0.00009 = 25.37 mg/kg).
# Repeat with the standard error dose to get standard range of error in mg/kg (18.82 mg/kg).


```


```{r Steatoda nobilis ED50 and LD50}

#############################
### First, calculate ED50 ###
#############################

### this subsets dataset

S_nobilis_ED50_crickets <- ED50_data_peaks[ED50_data_peaks$species == "Steatoda_nobilis" & ED50_data_peaks$prey_species == "Acheta_domesticus", ]

model_S_nobilis_ED50_crickets <- glm(cbind(S_nobilis_ED50_crickets$incapacitated_or_dead, 
                                           S_nobilis_ED50_crickets$alive) ~ (S_nobilis_ED50_crickets$venom_conc_percent), 
                                           binomial(link = probit))
                                       
summary(model_S_nobilis_ED50_crickets)


### plotting the data

### this just generates a sequence of numbers
ypredicts <- seq(0, 1, 0.01)
xpredicts <- dose.p(model_S_nobilis_ED50_crickets,p=c(ypredicts))


plot(S_nobilis_ED50_crickets$incapacitated_or_dead/20 ~ (S_nobilis_ED50_crickets$venom_conc_percent),
     ylab = "Proportion_incap",
     xlab = "Dosage",
     pch = 16)

lines(unlist(xpredicts), ypredicts)


S_nobilis_ED50_crickets_result <- dose.p(model_S_nobilis_ED50_crickets, p=c(0.5))
S_nobilis_ED50_crickets_result
#                 Dose         SE
#  p = 0.5: 0.01099826 0.01004753


### Converting % dose to mg/kg ###

# The calculated dose is the % concentration dose (mg/ul) required to achieve an ED50 in crickets of median mass 0.0003 kg.
# To get ED50 mg/kg, you first figure out how much 100% of the sample equals (0.0988002822865208 mg/ul).
# 100% then divided by 100 to get 1% = 0.000988 mg/ul.
# 1% multiplied by % dose = 0.011 = 0.000010868 mg/ul.
# 0.000010868 mg/ul is then divided by cricket median mass to get mg/kg (0.000010868/0.0003 = 0.036 mg/kg).
# Repeat with the standard error dose to get standard range of error in mg/kg (0.033 mg/kg).


### this subsets dataset (woodlice)

S_nobilis_ED50_woodlice <- ED50_data_peaks[ED50_data_peaks$species == "Steatoda_nobilis" & ED50_data_peaks$prey_species == "Porcellio_scaber", ]


model_S_nobilis_ED50_woodlice <- glm(cbind(S_nobilis_ED50_woodlice$incapacitated_or_dead, 
                                           S_nobilis_ED50_woodlice$alive) ~ (S_nobilis_ED50_woodlice$venom_conc_percent),
                                           binomial(link = probit))
                                       
summary(model_S_nobilis_ED50_woodlice)


### plotting the data

### this just generates a sequence of numbers
ypredicts <- seq(0, 1, 0.01)
xpredicts <- dose.p(model_S_nobilis_ED50_woodlice,p=c(ypredicts))


plot(S_nobilis_ED50_woodlice$incapacitated_or_dead/20 ~ (S_nobilis_ED50_woodlice$venom_conc_percent),
     ylab = "Proportion_incap",
     xlab = "Dosage",
     pch = 16)

lines(unlist(xpredicts), ypredicts)


S_nobilis_ED50_woodlice_result <- dose.p(model_S_nobilis_ED50_woodlice, p=c(0.5))
S_nobilis_ED50_woodlice_result
#                  Dose         SE
#   p = 0.5: 0.07870573 0.07537188


### Converting % dose to mg/kg ###

# The calculated dose is the % concentration dose (mg/ul) required to achieve an ED50 in woodlice of median mass 0.00009 kg.
# To get ED50 mg/kg, you first figure out how much 100% of the sample equals (0.0988002822865208 mg/ul).
# 100% then divided by 100 to get 1% = 0.000988 mg/ul.
# 1% multiplied by % dose = 0.079 = 0.000078 mg/ul.
# 0.000078 mg/ul is then divided by woodlice median mass to get mg/kg (0.000078/0.00009 = 0.87 mg/kg).
# Repeat with the standard error dose to get standard range of error in mg/kg (0.82 mg/kg).


##########################
### Now calculate LD50 ###
##########################


### this subsets dataset

S_nobilis_LD50_crickets <- LD50_data24[LD50_data24$species == "Steatoda_nobilis" & LD50_data24$prey_species == "Acheta_domesticus", ]

model_S_nobilis_LD50_crickets <- glm(cbind(S_nobilis_LD50_crickets$dead, 
                                           S_nobilis_LD50_crickets$alive) ~ (S_nobilis_LD50_crickets$venom_conc_percent), 
                                           binomial(link = probit))
                                       
summary(model_S_nobilis_LD50_crickets)


### plotting the data

### this just generates a sequence of numbers
ypredicts <- seq(0, 1, 0.01)
xpredicts <- dose.p(model_S_nobilis_LD50_crickets,p=c(ypredicts))


plot(S_nobilis_LD50_crickets$dead/20 ~ (S_nobilis_LD50_crickets$venom_conc_percent),
     ylab = "Proportion_dead",
     xlab = "Dosage",
     pch = 16)

lines(unlist(xpredicts), ypredicts)


### LD50 ###
S_nobilis_LD50_crickets_result <- dose.p(model_S_nobilis_LD50_crickets, p=c(0.5))
S_nobilis_LD50_crickets_result
#          Dose      SE
# p = 0.5: 0.2200633 0.05887277


### Converting % dose to mg/kg ###

# The calculated dose is the % concentration dose (mg/ul) required to achieve an LD50 in crickets of median mass 0.0003 kg.
# To get LD50 mg/kg, you first figure out how much 100% of the sample equals (0.098793 mg/ul).
# 100% then divided by 100 to get 1% = 0.000988 mg/ul.
# 1% multiplied by % dose = 0.22 = 0.000217 mg/ul.
# 0.000217 mg/ul is then divided by cricket median mass to get mg/kg (0.000217/0.0003 = 0.72 mg/kg).
# Repeat with the standard error dose to get standard range of error in mg/kg (0.2 mg/kg).


### this subsets dataset (woodlice)

S_nobilis_LD50_woodlice <- LD50_data24[LD50_data24$species == "Steatoda_nobilis" & LD50_data24$prey_species == "Porcellio_scaber", ]


model_S_nobilis_LD50_woodlice <- glm(cbind(S_nobilis_LD50_woodlice$dead, 
                                           S_nobilis_LD50_woodlice$alive) ~ (S_nobilis_LD50_woodlice$venom_conc_percent),
                                           binomial(link = probit))
                                       
summary(model_S_nobilis_LD50_woodlice)


### plotting the data

### this just generates a sequence of numbers
ypredicts <- seq(0, 1, 0.01)
xpredicts <- dose.p(model_S_nobilis_LD50_woodlice,p=c(ypredicts))


plot(S_nobilis_LD50_woodlice$dead/20 ~ (S_nobilis_LD50_woodlice$venom_conc_percent),
     ylab = "Proportion_dead",
     xlab = "Dosage",
     pch = 16)

lines(unlist(xpredicts), ypredicts)


### LD50 ###
S_nobilis_LD50_woodlice_result <- dose.p(model_S_nobilis_LD50_woodlice, p=c(0.5))
S_nobilis_LD50_woodlice_result
#          Dose      SE
# p = 0.5: 0.6807888 0.6111437
# 

### Converting % dose to mg/kg ###

# The calculated dose is the % concentration dose (mg/ul) required to achieve an LD50 in woodlice of median mass 0.00009 kg.
# To get LD50 mg/kg, you first figure out how much 100% of the sample equals (0.098793 mg/ul).
# 100% then divided by 100 to get 1% = 0.000988 mg/ul.
# 1% multiplied by % dose = 0.68 = 0.000672 mg/ul.
# 0.000672 mg/ul is then divided by woodlice median mass to get mg/kg (0.000672/0.00009 = 7.47 mg/kg).
# Repeat with the standard error dose to get standard range of error in mg/kg (6.7 mg/kg).


########################################
##### Let's try with 1hr peak data #####
########################################


#################################
### First, calculate ED50 1hr ###
#################################

### this subsets dataset

S_nobilis_ED50_crickets_1hr <- ED50_data_peaks_1hr[ED50_data_peaks_1hr$species == "Steatoda_nobilis" & ED50_data_peaks_1hr$prey_species == "Acheta_domesticus", ]

model_S_nobilis_ED50_crickets_1hr <- glm(cbind(S_nobilis_ED50_crickets_1hr$incapacitated_or_dead, 
                                           S_nobilis_ED50_crickets_1hr$alive) ~ (S_nobilis_ED50_crickets_1hr$venom_conc_percent), 
                                           binomial(link = probit))
                                       
summary(model_S_nobilis_ED50_crickets_1hr)


### plotting the data

### this just generates a sequence of numbers
ypredicts <- seq(0, 1, 0.01)
xpredicts <- dose.p(model_S_nobilis_ED50_crickets_1hr,p=c(ypredicts))


plot(S_nobilis_ED50_crickets_1hr$incapacitated_or_dead/20 ~ (S_nobilis_ED50_crickets_1hr$venom_conc_percent),
     ylab = "Proportion_incap",
     xlab = "Dosage",
     pch = 16)

lines(unlist(xpredicts), ypredicts)


S_nobilis_ED50_crickets_result_1hr <- dose.p(model_S_nobilis_ED50_crickets_1hr, p=c(0.5))
S_nobilis_ED50_crickets_result_1hr
#           Dose       SE
#  p = 0.5: 0.01582886 0.008639478


### Converting % dose to mg/kg ###

# The calculated dose is the % concentration dose (mg/ul) required to achieve an ED50 in crickets of median mass 0.0003 kg.
# To get ED50 mg/kg, you first figure out how much 100% of the sample equals (0.0988002822865208 mg/ul).
# 100% then divided by 100 to get 1% = 0.000988 mg/ul.
# 1% multiplied by % dose = 0.016 = 0.000015808 mg/ul.
# 0.000015808 mg/ul is then divided by cricket median mass to get mg/kg (0.000015808/0.0003 = 0.053 mg/kg).
# Repeat with the standard error dose to get standard range of error in mg/kg (0.028 mg/kg).


### this subsets dataset (woodlice)

S_nobilis_ED50_woodlice_1hr <- ED50_data_peaks_1hr[ED50_data_peaks_1hr$species == "Steatoda_nobilis" & ED50_data_peaks_1hr$prey_species == "Porcellio_scaber", ]


model_S_nobilis_ED50_woodlice_1hr <- glm(cbind(S_nobilis_ED50_woodlice_1hr$incapacitated_or_dead, 
                                           S_nobilis_ED50_woodlice_1hr$alive) ~ (S_nobilis_ED50_woodlice_1hr$venom_conc_percent),
                                           binomial(link = probit))
                                       
summary(model_S_nobilis_ED50_woodlice_1hr)


### plotting the data

### this just generates a sequence of numbers
ypredicts <- seq(0, 1, 0.01)
xpredicts <- dose.p(model_S_nobilis_ED50_woodlice_1hr,p=c(ypredicts))


plot(S_nobilis_ED50_woodlice_1hr$incapacitated_or_dead/20 ~ (S_nobilis_ED50_woodlice_1hr$venom_conc_percent),
     ylab = "Proportion_incap",
     xlab = "Dosage",
     pch = 16)

lines(unlist(xpredicts), ypredicts)


S_nobilis_ED50_woodlice_result_1hr <- dose.p(model_S_nobilis_ED50_woodlice_1hr, p=c(0.5))
S_nobilis_ED50_woodlice_result_1hr
#            Dose     SE
#   p = 0.5: 0.174228 0.1263925


### Converting % dose to mg/kg ###

# The calculated dose is the % concentration dose (mg/ul) required to achieve an ED50 in woodlice of median mass 0.00009 kg.
# To get ED50 mg/kg, you first figure out how much 100% of the sample equals (0.0988002822865208 mg/ul).
# 100% then divided by 100 to get 1% = 0.000988 mg/ul.
# 1% multiplied by % dose = 0.17 = 0.00016796 mg/ul.
# 0.00016796 mg/ul is then divided by woodlice median mass to get mg/kg (0.00016796/0.00009 = 1.87 mg/kg).
# Repeat with the standard error dose to get standard range of error in mg/kg (1.39 mg/kg).


```


###########################################################################################################################
############################################ ED50 ~ LD50 Relationship ANALYSIS ############################################
###########################################################################################################################


```{r data analysis subsets}

### To make the comparative analysis subsets 
ED50_LD50_comp_data <- droplevels(bioassay_data_raw[!bioassay_data_raw$comp_rep != 'rep',])

ED50_LD50_comp_data_v2 <- droplevels(ED50_LD50_comp_data[!ED50_LD50_comp_data$species == 'Cupiennius_coccineus',])

ED50_LD50_comp_data_v2$ED50_mgkg_1hr <- as.numeric(ED50_LD50_comp_data_v2$ED50_mgkg_1hr)


### For cricket only version of dataset 
cricket_only_comp <- droplevels(ED50_LD50_comp_data[!ED50_LD50_comp_data$prey_species == 'Porcellio_scaber',])

cricket_only_comp_v2 <- droplevels(ED50_LD50_comp_data_v2[!ED50_LD50_comp_data_v2$prey_species == 'Porcellio_scaber',])

ED50_data_peaks_cricket <- droplevels(ED50_data_peaks[!ED50_data_peaks$prey_species == 'Porcellio_scaber',])

ED50_data_peaks_1hr_cricket <- droplevels(ED50_data_peaks_1hr[!ED50_data_peaks_1hr$prey_species == 'Porcellio_scaber',])


### For woodlice only version of dataset 
woodlice_only_comp <- droplevels(ED50_LD50_comp_data[!ED50_LD50_comp_data$prey_species == 'Acheta_domesticus',])

woodlice_only_comp_v2 <- droplevels(ED50_LD50_comp_data_v2[!ED50_LD50_comp_data_v2$prey_species == 'Acheta_domesticus',])

ED50_data_peaks_woodlice <- droplevels(ED50_data_peaks[!ED50_data_peaks$prey_species == 'Acheta_domesticus',])

ED50_data_peaks_1hr_woodlice <- droplevels(ED50_data_peaks_1hr[!ED50_data_peaks_1hr$prey_species == 'Acheta_domesticus',])
```


```{r Main GLM and Supplementary GLM S1: Relationship between ED50 and LD50}

#########################################
### Main Glm: LD50 vs ED50 (4 hr) ###
#########################################

main_glm <- glm((log10(ED50_mgkg)) ~ (log10(LD50_mgkg)*prey_species),
                      data = ED50_LD50_comp_data)

summary(main_glm)
# without size or silk *prey models #
#                                                Estimate  Std.Error t value Pr(>|t|)   
#  (Intercept)                                    -0.8228     0.2809  -2.930 0.008287 ** 
#  log10(LD50_mgkg)                                1.0094     0.2396   4.213 0.000428 ***
#  prey_speciesPorcellio_scaber                    1.0184     0.4993   2.040 0.054800 .  
#  log10(LD50_mgkg):prey_speciesPorcellio_scaber  -0.4788     0.3485  -1.374 0.184697   


#####################################
### GLM S1: LD50 vs ED50 (1 hr) ###
#####################################

glm_s1 <- glm((log10(ED50_mgkg_1hr)) ~ (log10(LD50_mgkg)*prey_species),
                      data = ED50_LD50_comp_data_v2)

summary(glm_s1)
#                                               Estimate Std. Error t value Pr(>|t|)    
#(Intercept)                                     -0.7346     0.2772  -2.650 0.016277 *  
#log10(LD50_mgkg)                                 0.9496     0.2339   4.060 0.000735 ***
#prey_speciesPorcellio_scaber                     1.2701     0.4940   2.571 0.019236 *  
#log10(LD50_mgkg):prey_speciesPorcellio_scaber   -0.5658     0.3409  -1.659 0.114337


```


#####################################################################################################
# For plotting main GLM 1, manually copy and paste the code below into the console, step by step. #
#####################################################################################################

# First, the "windows" code produces an extra tab for visualising the plot.

 windows(30, 30)

# After running the code below and it shows the first scatterplot, press Esc.
# Then use abline code to plot the line.

plot(log10(ED50_mgkg) ~ (log10(LD50_mgkg)*prey_species),
     data = ED50_LD50_comp_data,
     pch = 16,
     col = "purple",
     bty =  "n")
     
# The points #  

points(log10(ED50_LD50_comp_data[ED50_LD50_comp_data$prey_species == "Porcellio_scaber", "ED50_mgkg"])
       ~ log10(ED50_LD50_comp_data[ED50_LD50_comp_data$prey_species == "Porcellio_scaber", "LD50_mgkg"]),
       type = "p",
       pch = 16,
       col = "#79addc")

points(log10(ED50_LD50_comp_data[ED50_LD50_comp_data$prey_species == "Acheta_domesticus", "ED50_mgkg"])
       ~ log10(ED50_LD50_comp_data[ED50_LD50_comp_data$prey_species == "Acheta_domesticus", "LD50_mgkg"]),
       type = "p",
       pch = 16,
       col = "#f46200")
       
# The lines #       

# Cricket/main line #    
abline(main_glm$coefficients[1],
       main_glm$coefficients[2],
       col = "black",
       lwd = 3)

# The legend #

legend(1.5, -0.5, legend = c("Woodlice", "Crickets"),
       col = c("#79addc", "#f46200"),
       pch = 16,
       cex = 0.8,
       title = "Prey Model",
       text.font = 20, 
       bg = "white")   

       
##########################################################################################################################    
# For plotting Supplementary GLM S1 (ED50 1hr), manually copy and paste the code below into the console, step by step. #
##########################################################################################################################

# First, the "windows" code produces an extra tab for visualising the plot.

 windows(30, 30)

# After running the code below and it shows the first scatterplot, press Esc.
# Then use abline code to plot the line.

plot(log10(ED50_mgkg_1hr) ~ (log10(LD50_mgkg)*prey_species),
     data = ED50_LD50_comp_data_v2,
     pch = 16,
     col = "purple",
     bty =  "n")
     
# The points #  

points(log10(ED50_LD50_comp_data_v2[ED50_LD50_comp_data_v2$prey_species == "Porcellio_scaber", "ED50_mgkg_1hr"])
       ~ log10(ED50_LD50_comp_data_v2[ED50_LD50_comp_data_v2$prey_species == "Porcellio_scaber", "LD50_mgkg"]),
       type = "p",
       pch = 16,
       col = "#79addc")

points(log10(ED50_LD50_comp_data_v2[ED50_LD50_comp_data_v2$prey_species == "Acheta_domesticus", "ED50_mgkg_1hr"])
       ~ log10(ED50_LD50_comp_data_v2[ED50_LD50_comp_data_v2$prey_species == "Acheta_domesticus", "LD50_mgkg"]),
       type = "p",
       pch = 16,
       col = "#f46200")
       
# The lines #       
 
# Cricket/main line #     
abline(glm_s1$coefficients[1],
       glm_s1$coefficients[2],
       col = "black",
       lwd = 3)


# The legend #

legend(1.5, -0.5, legend = c("Woodlice", "Crickets"),
       col = c("#79addc", "#f46200"),
       pch = 16,
       cex = 0.8,
       title = "Prey Model",
       text.font = 20, 
       bg = "white")   
       
       
       
```{r Supplementary GLM S2: Main GLM with P. phalangioides excluded }

# To make El only dataset #
ED50_LD50_EL_only <- droplevels(ED50_LD50_comp_data[!ED50_LD50_comp_data$species == 'Pholcus_phalangioides',])

# model #
glm_s2 <- glm((log10(ED50_mgkg)) ~ (log10(LD50_mgkg)*prey_species),
                           data = ED50_LD50_EL_only)

summary(glm_s2)
#                                                Estimate Std. Error t value  Pr(>|t|)    
#  (Intercept)                                    -0.8065     0.3059  -2.637  0.016757 *  
#  log10(LD50_mgkg)                                1.0012     0.2548   3.929  0.000985 ***
#  prey_speciesPorcellio_scaber                    1.0570     0.5565   1.899  0.073667 .  
#  log10(LD50_mgkg):prey_speciesPorcellio_scaber  -0.4976     0.3773  -1.319  0.203794 


```
 
 
#######################################################################################################################
# For plotting Supplementary GLM S2 (EL only), manually copy and paste the code below into the console, step by step. #
#######################################################################################################################

# First, the "windows" code produces an extra tab for visualising the plot.

 windows(30, 30)

# After running the code below and it shows the first scatterplot, press Esc.
# Then use abline code to plot the line.

plot(log10(ED50_mgkg) ~ (log10(LD50_mgkg)*prey_species),
     data = ED50_LD50_EL_only,
     pch = 16,
     col = "purple",
     bty =  "n")
     
# The points #  

points(log10(ED50_LD50_EL_only[ED50_LD50_EL_only$prey_species == "Porcellio_scaber", "ED50_mgkg"])
       ~ log10(ED50_LD50_EL_only[ED50_LD50_EL_only$prey_species == "Porcellio_scaber", "LD50_mgkg"]),
       type = "p",
       pch = 16,
       col = "#79addc")

points(log10(ED50_LD50_EL_only[ED50_LD50_EL_only$prey_species == "Acheta_domesticus", "ED50_mgkg"])
       ~ log10(ED50_LD50_EL_only[ED50_LD50_EL_only$prey_species == "Acheta_domesticus", "LD50_mgkg"]),
       type = "p",
       pch = 16,
       col = "#f46200")
       
# The lines # 

# Cricket/main line #     
abline(glm_s2$coefficients[1],
       glm_s2$coefficients[2],
       col = "black",
       lwd = 3)


# The legend #

legend(1.5, -0.5, legend = c("Woodlice", "Crickets"),
       col = c("#79addc", "#f46200"),
       pch = 16,
       cex = 0.8,
       title = "Prey Model",
       text.font = 20, 
       bg = "white") 
 
 
    
```{r Main MCMCglmm: ED50 vs LD50 with ul/g lit data}

S2_bioassays <- read.csv("S2_ulg_lit_data.csv",
                             sep = ",",
                             header = T)

S2_bioassays <- S2_bioassays[,c("species",
                                "taxa",
                                "animal",
                                "species_matched", 
                                "prey_species", 
                                "prey_model_mass_g",
                                "model_class",
                                "Cohort_N" , 
                                "LD50_ulg", 
                                "ED50_ulg",
                                "extraction_method",
                                "injection_method")] 
                                 
S2_bioassays <- na.omit(S2_bioassays)


S2_bioassays$model_class <- factor(S2_bioassays$model_class,
                                       levels = c("Insecta", "Arachnida"))


### For cricket only version of dataset 
Insect_only_comp <- droplevels(S2_bioassays[!S2_bioassays$model_class == 'Arachnida',])

### For woodlice only version od dataset 
Arachnid_only_comp <- droplevels(S2_bioassays[!S2_bioassays$model_class == 'Insecta',])


####################
### r parameters ###
####################

### In order to include multiple measures of LD50 for each species in our analysis, 
### we need a random term for species (spec).
### The code for replacing genus with species (spec) ###

spec <- (gsub("_._","", S2_bioassays$species))
S2_bioassays <- data.frame(S2_bioassays, spec = spec)

### In order to include multiple measures of LD50 for each family in our analysis,
### we need a random term for family (fam).
### the code for adding family (fam) as a random term ###

fam <- (gsub("_","", S2_bioassays$taxa))
S2_bioassays <- data.frame(S2_bioassays, fam = fam)


##################################################
### Upload Phylogeny for species in S2 dataset ###
##################################################

lit_data_phylo <- read.tree("Lyons_et_al_TX_phylo.tre")


###########################
# r priors and parameters #
###########################

### Set up priors ###
### Will need a prior for incorporating multiple random terms ###

prior_mixed_x2_random <- list(G = list(G1 = list(nu=0.002, V=1), G2 = list(nu=0.002, V=1)),
                              R = list(nu=0.002, V=1))
prior_mixed_x2_random

### Parameters ###

nitt3 <- 2400000
burnin3 = 40000
thin3 = 100


########################################################
### Main MCMCglmm ED50 vs LD50 (ul venom per g prey) ###
########################################################

main_mcmcglmm <- MCMCglmm(log10(ED50_ulg) ~ log10(LD50_ulg),
                          data = S2_bioassays,
                          rcov = ~ units, 
                          random = ~ animal + spec,
                          family ="gaussian",
                          pedigree = lit_data_phylo,
                          nitt = nitt3,
                          burnin = burnin3,
                          thin = thin3,
                          prior = prior_mixed_x2_random,
                          verbose = FALSE)


summary(main_mcmcglmm)
#                post.mean l-95% CI u-95% CI eff.samp  pMCMC    
#(Intercept)       -0.5194  -0.9652  -0.1165    22259 0.0153 *  
#log10(LD50_ulg)    1.1037   0.8110   1.3923    23112 <4e-05 ***


####################################
### Data for Main MCMCglmm table ###
####################################

### Animal term (Phylogeny h2) ###

posterior.heritability_main_mcmcglmm_animal <-main_mcmcglmm$VCV[,"animal"]/
  +     + (main_mcmcglmm$VCV[,"animal"] + main_mcmcglmm$VCV[,"units"] 
        + main_mcmcglmm$VCV[,"spec"])

HPDinterval(posterior.heritability_main_mcmcglmm_animal,0.95)
# lower        upper
# 0.0002309618 0.3657627


posterior.mode(posterior.heritability_main_mcmcglmm_animal)
# var1 
# 0.002585699 


### species ###

posterior.heritability_main_mcmcglmm_spec <-main_mcmcglmm$VCV[,"spec"]/
  +     + (main_mcmcglmm$VCV[,"spec"] + main_mcmcglmm$VCV[,"animal"] 
        + main_mcmcglmm$VCV[,"units"])

HPDinterval(posterior.heritability_main_mcmcglmm_spec,0.95)
# lower         upper
# 0.0001510116 0.2575498

posterior.mode(posterior.heritability_main_mcmcglmm_spec)
# var1 
# 0.002246521 


### units (residuals) ###

posterior.heritability_main_mcmcglmm_units <-main_mcmcglmm$VCV[,"units"]/
  +     + (main_mcmcglmm$VCV[,"units"] + main_mcmcglmm$VCV[,"animal"] 
           + main_mcmcglmm$VCV[,"spec"])

HPDinterval(posterior.heritability_main_mcmcglmm_units,0.95)
# lower      upper
# 0.5621099 0.9987361

posterior.mode(posterior.heritability_main_mcmcglmm_units)
# var1 
# 0.991333


```

# For plotting the main MCMCglmm, manually copy and paste the code below into the console, step by step.

# First, the "windows" code produces an extra tab for visualising the plot.

 windows(30, 30)

# After running the code below and it shows the first scatterplot, press Esc.
# Then use abline code to plot the line.

 plot(log10(ED50_ulg) ~ log10(LD50_ulg),
     data = S2_bioassays,
     pch = 16,
     col = "#79addc",
     bty =  "n",
     xlim = c(-3.0, 2.0),
     ylim = c(-4.0, 2.0))


# The line #

# The abline requires the following:
# intercept + log10(median body length^2.6)*body length slope, EVO_DIST slope 
# log10(median body length) added on as body length acts as a z axis for this plot 
# NB! You will need to replace these numbers with the ones produced during your model run

 abline(-0.5194, 1.1037,
       lwd = 3,
       col = "#79addc")


####################################################################################################################################
####################################################################################################################################
````
