## Supplementary material for "Paralysis Efficiency (ED_50_) Scales Linearly with Lethality (LD_50_) in Spider Venoms": S4: S4_phylo_for_S2_TX.Rmd

````
----------------
          Title: "S4_phylo_for_Dataset_S2_(Literature data)"
         Author: "Keith Lyons"
Purpose of file: "to demonstrate how the phylogeny was created"
           Date: "16-09-2025"
         Output: html_document
----------------

```{r load packages}

library(ggplot2)
library(MASS)
library(MCMCglmm)
library(lme4)
library(invgamma)
library(mulTree)
library(caper)
library(phytools)
library(picante)
library(ape)
library(castor)
library(geiger)
library(corHMM)

### Installing the mulTree package if it doesn't work ###

# install.packages("devtools")
# library(devtools)
# install_github("TGuillerme/mulTree", ref = "release")
# library(mulTree)
```


```{r functions}

### This function is required to add species to the base phylogeny ###
### The base phylogeny was obtained from a published study (see later) ###

bind.tip<-function(tree,tip.label,edge.length=NULL,where=NULL){
  if(is.null(where)) where<-length(tree$tip)+1
  tip<-list(edge=matrix(c(2,1),1,2),
            tip.label=tip.label,
            edge.length=edge.length,
            Nnode=1)
  class(tip)<-"phylo"
  obj<-bind.tree(tree,tip,where=where)
  return(obj)}


```


Before running the code below, go to Session tab above -> Set Working Directory -> Choose Directory,
                                                                                   select location of data file S2
                                                                                   
If code doesn't run despite the correct directory being set, manually copy and paste the code below into the console.                                                                                   

```{r upload data and subsetting}

S2_bioassays <- read.csv("S2_ulg_lit_data.csv",
                             sep = ",",
                             header = T)

S2_bioassays <- S2_bioassays[,c("species",
                                "taxa",
                                "animal",
                                "species_matched",
                                "prey_species", 
                                "prey_model_mass_g",
                                "model_class",
                                "Cohort_N" , 
                                "LD50_ulg", 
                                "ED50_ulg",
                                "extraction_method",
                                "injection_method")] 
                                 


S2_bioassays$model_class <- factor(S2_bioassays$model_class,
                                       levels = c("Insecta", "Arachnida"))


### For cricket only version of dataset 
Insect_only_comp <- droplevels(S2_bioassays[!S2_bioassays$model_class == 'Arachnida',])

### For woodlice only version od dataset 
Arachnid_only_comp <- droplevels(S2_bioassays[!S2_bioassays$model_class == 'Insecta',])

```


```{r random terms}

### In order to include multiple measures of LD50 for each species in our analysis, 
### we need a random term for species (spec).
### The code for replacing genus with species (spec) ###

spec <- (gsub("_._","", S2_bioassays$species))
S2_bioassays <- data.frame(S2_bioassays, spec = spec)

### In order to include multiple measures of LD50 for each family in our analysis,
### we need a random term for family (fam).
### the code for adding family (fam) as a random term ###

fam <- (gsub("_","", S2_bioassays$taxa))
S2_bioassays <- data.frame(S2_bioassays, fam = fam)

```


 Backbone of phylogenetic tree is adapted from Wolff et al. 2022., 
 Stabilized morphological evolution of spiders despite mosaic changes in foraging ecology. Syst Biol.

```{r phylo}

### The phylo file from Wolff et al is included in the S4 folder
### Base phylo name = "consensus_rooted7_12dated.tre"
phylo_Wolff <- read.tree("consensus_rooted7_12dated.tre")

dummy_data <- data.frame(species = unique(S2_bioassays$animal), 
                         animal = unique(S2_bioassays$animal))


comp_data <- comparative.data(phy = phylo_Wolff, 
                              data = dummy_data, 
                              names.col = animal, 
                              vcv=TRUE)

spider_dataset_matched <- S2_bioassays[S2_bioassays$animal %in% S2_bioassays$species_matched,]

spider_phylo <- comp_data$phy

# windows(30, 30)
plot(spider_phylo,
     cex = 0.5,)
```

# Salticidae

Add Salticidae_Phidippus_octopunctatus, Salticidae_Phidippus_audax, Salticidae_Phidippus_johnsoni 
and 
Salticidae_Phidippus_ardens as a sister group to Salticidae_Plexippus_paykulli 
and 
then remove Salticidae_Plexippus_paykulli 
based on evidence from: 
 Guerrero-Fuentes, D.R., Francke-Ballvé, O.F. and Ceccarelli, F.S., 2024. Molecular phylogeny of the large-bodied dendryphantines of North America: Paraphidippus, Phidippus, Pelegrina, and their relatives (Salticidae: Dendryphantina). Insect Systematics and Diversity, 8(6), p.9.
AND
  Edwards, G.B., 2004. Revision of the jumping spiders of the genus Phidippus. Occasional Papers of the Florida State Collection of Arthropods, 11, pp.1-156.
AND
  Maddison, W.P. and Hedin, M.C., 2003. Jumping spider phylogeny (Araneae: Salticidae). Invertebrate systematics, 17(4), pp.529-549.

Also match Eutichuridae_Cheiracanthium_mildei as a basel node to Salticidae based on: 
  Azevedo, Guilherme HF, Charles E. Griswold, and Adalberto J. Santos."Systematics and evolution of ground spiders revisited (Araneae, Dionycha, Gnaphosidae)." Cladistics 34, no. 6 (2018): 579-626.    


```{r Salticidae}

Salticidae <- read.tree(text = "(((((((((Salticidae_Phidippus_ardens:35,Salticidae_Phidippus_johnsoni:35):4, Salticidae_Phidippus_audax:40):2):2, Salticidae_Phidippus_octopunctatus:45):2):2, Salticidae_Plexippus_paykulli:50):20,Philodromidae_Tibellus_oblongus:70):5):5,Eutichuridae_Cheiracanthium_mildei:80):5;")

Salticidae$root.edge <- 35

plot(Salticidae, cex = 0.5)

spider_phylo$tip.label[spider_phylo$tip.label == "Salticidae_Plexippus_paykulli"] <- "Salticidae_Plexippus_rm"
plot(spider_phylo, cex = 0.5)

spider_phylo$tip.label[spider_phylo$tip.label == "Philodromidae_Tibellus_oblongus"] <- "Philodromidae_Tibellus_oblongus_rm"
plot(spider_phylo, cex = 0.5)

spider_phylo$tip.label[spider_phylo$tip.label == "Eutichuridae_Cheiracanthium_mildei"] <- "Eutichuridae_Cheiracanthium_mildei_rm"
plot(spider_phylo, cex = 0.5)


Salti_add_tree <- bind.tree(spider_phylo, Salticidae, where = 56)
# 55 is the node label
plot(Salti_add_tree, cex = 0.5)

# Plexippus_rm can now be removed
Salti_add_treeF <- drop.tip(Salti_add_tree, c("Salticidae_Plexippus_rm"))
plot(Salti_add_treeF, cex = 0.5)

# T. oblongus was added to salti tree so this tip can be removed from previous position
Salti_add_treeF1 <- drop.tip(Salti_add_treeF, c("Philodromidae_Tibellus_oblongus_rm"))
plot(Salti_add_treeF1, cex = 0.5)

# C. mildei was added to salti tree so this tip can be removed from previous position
Salti_add_treeF1 <- drop.tip(Salti_add_treeF1, c("Eutichuridae_Cheiracanthium_mildei_rm"))
# windows(30, 30)
plot(Salti_add_treeF1, cex = 0.5)


```


Add Lycosidae_Tigrosa_helluo as a sister group to Lycosidae_Schizocosa_ocreata (inference for Schizocosa_mccooki), Lycosidae_Hogna frondicola (inference for Hogna carolinensis) and Lycosidae_Trochosa_ruricola (inference for Trochosa sp.)
based on:
  Piacentini, L.N. and Ramírez, M.J., 2019. Hunting the wolf: a molecular phylogeny of the wolf spiders (Araneae, Lycosidae). Molecular Phylogenetics and Evolution, 136, pp.227-240.


```{r Lycosidae}

Lycosidae <- read.tree(text = "(((((Lycosidae_Schizocosa_ocreata:1,Lycosidae_Tigrosa_helluo:1),Lycosidae_Hogna_frondicola:1):1,Lycosidae_Trochosa_ruricola:1):1):1);")

Lycosidae <- read.tree(text = "(((((Lycosidae_Schizocosa_ocreata:1,Lycosidae_Tigrosa_helluo:1),Lycosidae_Trochosa_ruricola:1):1,Lycosidae_Hogna_frondicola:1):1):1);")


plot(Lycosidae, cex = 0.5)

Lycosidae_fix <- compute.brlen(Lycosidae)
# not sure how this edge length is determined so added nearby edge length 
Lycosidae_fix$edge.length <- Lycosidae_fix$edge.length*51.172660
Lycosidae_fix$root.edge <- 10

plot(Lycosidae_fix, cex = 0.5)
Lycosidae_fix$edge.length

# windows(30, 30)
plot(Salti_add_treeF1, cex = 0.5)
Salti_add_treeF1$tip.label[Salti_add_treeF1$tip.label == "Lycosidae_Hogna_frondicola"] <- "Lycosidae_Hogna_frondicola_rm"
plot(Salti_add_treeF1, cex = 0.5)

Salti_add_treeF1$tip.label[Salti_add_treeF1$tip.label == "Lycosidae_Schizocosa_ocreata"] <- "Lycosidae_Schizocosa_ocreata_rm"
plot(Salti_add_treeF1, cex = 0.5)

Salti_add_treeF1$tip.label[Salti_add_treeF1$tip.label == "Lycosidae_Trochosa_ruricola"] <- "Lycosidae_Trochosa_rm"
plot(Salti_add_treeF1, cex = 0.5)

Lyco_add_tree <- bind.tree(Salti_add_treeF1, Lycosidae_fix, where = 56)
plot(Lyco_add_tree, cex = 0.5)

Lyco_add_treeF <- drop.tip(Lyco_add_tree, c("Lycosidae_Hogna_frondicola_rm"))
Lyco_add_treeF <- drop.tip(Lyco_add_treeF, c("Lycosidae_Schizocosa_ocreata_rm"))
Lyco_add_treeF <- drop.tip(Lyco_add_treeF, c("Lycosidae_Trochosa_rm"))
plot(Lyco_add_treeF, cex = 0.5)

Lyco_add_treeF$node.label <- seq(length(Lyco_add_treeF$node.label))
plot(Lyco_add_treeF, cex = 0.5)

```

Add Agelenidae_Calilena_sp. as a sister group to Agelenidae_Hololena_curta
based on:
  Bolzern, A., Burckhardt, D. and Hänggi, A., 2013. Phylogeny and taxonomy of European funnel-web spiders of the Tegenaria–Malthonica complex (Araneae: Agelenidae) based upon morphological and molecular data. Zoological Journal of the Linnean Society, 168(4), pp.723-848.

```{r Agelenidae}

Agelenidae_fix <- read.tree(text = "(Agelenidae_Hololena_curta:11,Agelenidae_Calilena_sp.:11):1;")
Agelenidae_fix$root.edge <- 30
plot(Agelenidae_fix, cex = 0.5)


Lyco_add_treeF$tip.label[Lyco_add_treeF$tip.label == "Agelenidae_Hololena_curta"] <- "Agelenidae_Hololena_rm"
plot(Lyco_add_treeF, cex = 0.5)

Agel_add_tree <- bind.tree(Lyco_add_treeF, Agelenidae_fix, where = 53)
plot(Agel_add_tree, cex = 0.5)

Agel_add_treeF <- drop.tip(Agel_add_tree, c("Agelenidae_Hololena_rm"))
plot(Agel_add_treeF, cex = 0.5)

```


Add Tliltocatl_albopilosus, Avicularia_avicularia and Haplopus_limensis as a sister group to Theraphosidae_Aphonopelma_vorhiesi 
based on:
  Ortiz, D., Francke, O.F. and Bond, J.E., 2018. 
A tangle of forms and phylogeny: extensive morphological homoplasy and molecular 
clock heterogeneity in Bonnetina and related tarantulas. Molecular Phylogenetics 
and Evolution, 127, pp.55-73.

Aphonopelma chalcodes has inference species in tree (Aphonopelma vorhiesi)

```{r Theraphosidae_fix}

Theraphosidae_fix <- read.tree(text = "((Theraphosidae_Aphonopelma_vorhiesi:10,Theraphosidae_Aphonopelma_chalcodes:10):40,(Theraphosidae_Tliltocatl_albopilosus:30,Theraphosidae_Avicularia_avicularia:30, Theraphosidae_Hapalopus_limensis:30):20):20;")

Theraphosidae_fix$root.edge <- 45
plot(Theraphosidae_fix, cex = 0.5)


Agel_add_treeF$tip.label[Agel_add_treeF$tip.label == "Theraphosidae_Aphonopelma_vorhiesi"] <- "Theraphosidae_Aphonopelma_vorhiesi_rm"
plot(Agel_add_treeF, cex = 0.5)

Theraphosidae_add_tree <- bind.tree(Agel_add_treeF, Theraphosidae_fix, where = 37)
plot(Theraphosidae_add_tree, cex = 0.5)

Theraphosidae_add_treeF <- drop.tip(Theraphosidae_add_tree, c("Theraphosidae_Aphonopelma_vorhiesi_rm"))
# windows(30, 30)
plot(Theraphosidae_add_treeF, cex = 0.5)

```


Add Gnaphosidae = Haplodrassus_sp., Herpyllus_propinquous and Orodrassuss_sp. as a sister group to Gnaphosidae_Gnaphosa_lucifuga
Based on:
  Azevedo, Guilherme HF, Charles E. Griswold, and Adalberto J. Santos.
"Systematics and evolution of ground spiders revisited (Araneae, Dionycha, 
Gnaphosidae)." Cladistics 34, no. 6 (2018): 579-626.            
              
    ```{r Gnaphosidae}

Gnaphosidae_fix <- read.tree(text = "(((Gnaphosidae_Gnaphosa_lucifuga:9,Gnaphosidae_Herpyllus_propinquous:9):10,Gnaphosidae_Orodrassus_sp.:19,Gnaphosidae_Haplodrassus_sp.:19):1);")

Gnaphosidae_fix$root.edge <- 10
plot(Gnaphosidae_fix, cex = 0.5)


Theraphosidae_add_treeF$tip.label[Theraphosidae_add_treeF$tip.label == "Gnaphosidae_Gnaphosa_lucifuga"] <- "Gnaphosidae_Gnaphosa_lucifuga_rm"
plot(Theraphosidae_add_treeF, cex = 0.5)

Gnaphosidae_add_tree <- bind.tree(Theraphosidae_add_treeF, Gnaphosidae_fix, where = 76)
plot(Gnaphosidae_add_tree, cex = 0.5)

Gnaphosidae_add_treeF <- drop.tip(Gnaphosidae_add_tree, c("Gnaphosidae_Gnaphosa_lucifuga_rm"))
plot(Gnaphosidae_add_treeF, cex = 0.5)

```


Make ultranmetric

```{r ultra}

final_tree <- force.ultrametric(Gnaphosidae_add_treeF)
final_tree$node.label <- c()


dummy_data <- data.frame(species = unique(S2_bioassays$animal), 
                         animal = unique(S2_bioassays$animal))


comp_data_fin <- comparative.data(phy = final_tree, 
                                  data = dummy_data, 
                                  names.col = animal, 
                                  vcv=TRUE)

write.tree(file = "Lyons_et_al_TX_phylo.tre", comp_data_fin$phy)

lit_data_phylo <- read.tree("Lyons_et_al_TX_phylo.tre")
# windows(30, 30)
plot(lit_data_phylo, cex = 0.5)

```
````
