## Supplementary material for "Paralysis Efficiency (ED_50_) Scales Linearly with Lethality (LD_50_) in Spider Venoms": S5

*Corresponding Author

[d]Midlands Bug and Reptile Zoo, Longford, Ireland.

**ORCID ID**: **K.L.** 0000-0002-3572-0847, **D.L.** 0000-0002-2943-748X, **L.McS.** 0000-0002-8823-1828, **M.M.** 0009-0005-4235-9059, **B.L.C.** 0000-0002-0709-9678, **A.V.** 0000-0001-9148-0916, **J.P.D.** 0000-0002-6645-0472, **M.M.D.** 0000-0002-8567-819X, **K.H.** 0000-0002-3548-6253.

#### Supplementary S5: Supplementary Tables and Model Outputs

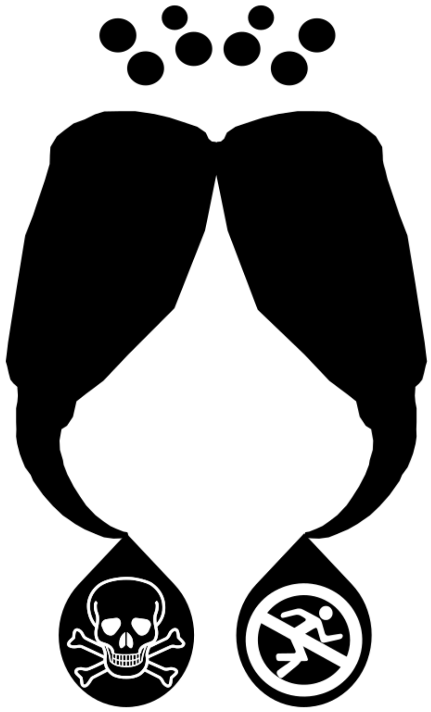

### Supplementary S5: Supplementary Tables and Model Outputs

**Table S1: Body length and venom yield data for the 12 spider species tested in the lab.** The data presented includes the number of specimens used for venom extraction, full spider body length (mm), the total liquid venom yield (µl) extracted, the total venom yield extracted expressed in milligram of lyophilised protein content per microliter of venom (mg/µl) and the venom yield expressed in milligram of dried venom extracted per specimen (mg/spider).

| Spider Species | Number of Specimens Extracted | Mean Full Body Length (mm) | Total Venom Yield (µl) | Total Venom Yield (mg/µl) (100% Concentration) | Venom Yield (mg/spider) |
|---|---|---|---|---|---|
| *Amaurobius similis* | 117 | 9.46 | 13.91 | 0.07908 | 0.0094 |
| *Cupiennius coccineus* | 5 (2 twice) | 16.3 | 4.73 | 0.05920 | 0.04 |
| *Cupiennius salei* | 6 (4 twice) | 15.32 | 8.05 | 0.04720 | 0.038 |
| *Eratigena atrica* | 99 | 12.01 | 42.33 | 0.15120 | 0.065 |
| *Heteropoda venatoria* | 10 (4 twice) | 18.32 | 44.55 | 0.10325 | 0.33 |
| *Larinioides sclopetarius* | 81 | 7.25 | 7.17 | 0.23152 | 0.0205 |
| *Meta menardi* | 63 | 11.6 | 15.75 | 0.07619 | 0.019 |
| *Monocentropus balfouri* | 4 | 46.43 | 19.9 | 0.11055 | 0.55 |
| *Pholcus phalangioides* | 140 | 6.9 | 7.9 | 0.08228 | 0.0046 |
| *Phormictopus cancerides* | 1 | 48.9 | 9.93 | 0.18429 | 1.83 |
| *Piloctenus haematostoma* | 4 | 26.48 | 30.29 | 0.00990 | 0.075 |
| *Steatoda nobilis* | 117 | 9.83 | 14.17 | 0.09880 | 0.012 |

**Table S2: Shows the venom concentrations that were tested for 12 spider species venoms during the bioassay experiments in both prey models**, House cricket *Acheta domesticus* and Common Rough woodlouse *Porcellio scaber* (n = 20 per cohort). Also presented are the number of spiders extracted to obtain each species venom sample, the total crude venom extracted (µl) and the 100% venom yield concentration obtained, expressed in milligram of lyophilised (dried) venom per microliter of venom (mg/µl).

| Spider Species | Number of Specimens Extracted | Total Venom Yield (µl) | Total Venom Yield (mg/µl) (100% Concentration) | Venom Concentration Injected (mg/µl) | Venom Concentration Injected (% Concentration) |
|---|---|---|---|---|---|
| *Amaurobius similis* | 117 | 13.91 | 0.07908 | 0.003164 | 4 |
| *Amaurobius similis* | 117 | 13.91 | 0.07908 | 0.000791 | 1 |
| *Amaurobius similis* | 117 | 13.91 | 0.07908 | 0.000158 | 0.2 |
| *Amaurobius similis* | 117 | 13.91 | 0.07908 | 0.000079 | 0.1 |
| *Amaurobius similis* | 117 | 13.91 | 0.07908 | 0.000039 | 0.05 |
| *Amaurobius similis* | 117 | 13.91 | 0.07908 | 0.000008 | 0.01 |
| *Cupiennius coccineus* | 5 (2 twice) | 4.73 | 0.05920 | 0.001184 | 2 |
| *Cupiennius coccineus* | 5 (2 twice) | 4.73 | 0.05920 | 0.000118 | 0.2 |
| *Cupiennius coccineus* | 5 (2 twice) | 4.73 | 0.05920 | 0.000012 | 0.02 |
| *Cupiennius salei* | 6 (4 twice) | 8.05 | 0.04720 | 0.001416 | 3 |

### Supplementary S5: Supplementary Tables and Model Outputs

| | | | | | |
|---|---|---|---|---|---|
| *Cupiennius salei* | 6 (4 twice) | 8.05 | 0.04720 | 0.000236 | 0.5 |
| *Cupiennius salei* | 6 (4 twice) | 8.05 | 0.04720 | 0.000047 | 0.1 |
| *Cupiennius salei* | 6 (4 twice) | 8.05 | 0.04720 | 0.000005 | 0.01 |
| *Eratigena atrica* | 99 | 42.33 | 0.15120 | 0.030239 | 20 |
| *Eratigena atrica* | 99 | 42.33 | 0.15120 | 0.01512 | 10 |
| *Eratigena atrica* | 99 | 42.33 | 0.15120 | 0.006048 | 4 |
| *Eratigena atrica* | 99 | 42.33 | 0.15120 | 0.003024 | 2 |
| *Eratigena atrica* | 99 | 42.33 | 0.15120 | 0.001512 | 1 |
| *Eratigena atrica* | 99 | 42.33 | 0.15120 | 0.000756 | 0.5 |
| *Eratigena atrica* | 99 | 42.33 | 0.15120 | 0.000151 | 0.1 |
| *Heteropoda venatoria* | 10 (4 twice) | 44.55 | 0.10325 | 0.008264 | 8 |
| *Heteropoda venatoria* | 10 (4 twice) | 44.55 | 0.10325 | 0.004132 | 4 |
| *Heteropoda venatoria* | 10 (4 twice) | 44.55 | 0.10325 | 0.001033 | 1 |
| *Heteropoda venatoria* | 10 (4 twice) | 44.55 | 0.10325 | 0.0005165 | 0.5 |
| *Heteropoda venatoria* | 10 (4 twice) | 44.55 | 0.10325 | 0.0001033 | 0.1 |
| *Heteropoda venatoria* | 10 (4 twice) | 44.55 | 0.10325 | 0.00001033 | 0.01 |
| *Larinioides sclopetarius* | 81 | 7.17 | 0.23152 | 0.002315 | 1 |
| *Larinioides sclopetarius* | 81 | 7.17 | 0.23152 | 0.0011575 | 0.5 |
| *Larinioides sclopetarius* | 81 | 7.17 | 0.23152 | 0.0002315 | 0.1 |
| *Meta menardi* | 63 | 15.75 | 0.07619 | 0.003048 | 4 |
| *Meta menardi* | 63 | 15.75 | 0.07619 | 0.000762 | 1 |
| *Meta menardi* | 63 | 15.75 | 0.07619 | 0.000571 | 0.75 |
| *Meta menardi* | 63 | 15.75 | 0.07619 | 0.000381 | 0.5 |
| *Meta menardi* | 63 | 15.75 | 0.07619 | 0.000076 | 0.1 |
| *Meta menardi* | 63 | 15.75 | 0.07619 | 0.000008 | 0.01 |
| *Monocentropus balfouri* | 4 | 19.9 | 0.11055 | 0.006633 | 6 |
| *Monocentropus balfouri* | 4 | 19.9 | 0.11055 | 0.002211 | 2 |
| *Monocentropus balfouri* | 4 | 19.9 | 0.11055 | 0.001105 | 1 |
| *Monocentropus balfouri* | 4 | 19.9 | 0.11055 | 0.000553 | 0.5 |
| *Monocentropus balfouri* | 4 | 19.9 | 0.11055 | 0.000111 | 0.1 |
| *Monocentropus balfouri* | 4 | 19.9 | 0.11055 | 0.000011 | 0.01 |
| *Pholcus phalangioides* | 140 | 7.9 | 0.08228 | 0.001646 | 2 |
| *Pholcus phalangioides* | 140 | 7.9 | 0.08228 | 0.000823 | 1 |
| *Pholcus phalangioides* | 140 | 7.9 | 0.08228 | 0.000411 | 0.5 |
| *Pholcus phalangioides* | 140 | 7.9 | 0.08228 | 0.000041 | 0.05 |
| *Phormictopus cancerides* | 1 | 9.93 | 0.18429 | 0.005529 | 3 |
| *Phormictopus cancerides* | 1 | 9.93 | 0.18429 | 0.0027645 | 1.5 |
| *Phormictopus cancerides* | 1 | 9.93 | 0.18429 | 0.001843 | 1 |
| *Phormictopus cancerides* | 1 | 9.93 | 0.18429 | 0.0009215 | 0.5 |
| *Phormictopus cancerides* | 1 | 9.93 | 0.18429 | 0.0001843 | 0.1 |
| *Piloctenus haematostoma* | 4 | 30.29 | 0.00990 | 0.00099 | 10 |
| *Piloctenus haematostoma* | 4 | 30.29 | 0.00990 | 0.000198 | 2 |
| *Piloctenus haematostoma* | 4 | 30.29 | 0.00990 | 0.000099 | 1 |
| *Piloctenus haematostoma* | 4 | 30.29 | 0.00990 | 0.0000099 | 0.1 |
| *Steatoda nobilis* | 117 | 14.17 | 0.09880 | 0.002964 | 3 |
| *Steatoda nobilis* | 117 | 14.17 | 0.09880 | 0.000988 | 1 |
| *Steatoda nobilis* | 117 | 14.17 | 0.09880 | 0.000494 | 0.5 |
| *Steatoda nobilis* | 117 | 14.17 | 0.09880 | 0.000099 | 0.1 |
| *Steatoda nobilis* | 117 | 14.17 | 0.09880 | 0.000049 | 0.05 |
| *Steatoda nobilis* | 117 | 14.17 | 0.09880 | 0.000025 | 0.025 |

| | | | | | |
|---|---|---|---|---|---|
| *Steatoda nobilis* | 117 | 14.17 | 0.09880 | 0.000012 | 0.0125 |
| *Steatoda nobilis* | 117 | 14.17 | 0.09880 | 0.000006 | 0.00625 |

**Table S3: Calculated potency measures for 12 spider species venoms** tested on house cricket *A. domesticus* and woodlouse *P. scaber*. Presented are the **$ED_{50}$ (mg/kg) (1 hr endpoint)** and **$LD_{50}$ (mg/kg) (24 hr endpoint)** values for the venom of each species, in both prey models, with standard errors, calculated using the dried venom protein mass (mg) and median prey model mass for crickets (0.0003 kg) or woodlice (0.00009 kg). Note *C. coccineus* venom failed to achieve an $ED_{50}$ in woodlice using the 1 hr endpoint. *A. domesticus* data for *Eratigena atrica* was produced as part of Lyons et al. (2023). $ED_{50}$ (1 hr) values that are different to their corresponding $ED_{50}$ (4 hr) values (see Table 2 in main manuscript) are shown in **bold**.

| | *Acheta domesticus* (cricket) | | | | *Porcellio scaber* (woodlouse) | | | |
|---|---|---|---|---|---|---|---|---|
| Spider Species | $ED_{50}$ mg/kg | Standard Error (-/+) | $LD_{50}$ mg/kg | Standard Error (-/+) | $ED_{50}$ mg/kg | Standard Error (-/+) | $LD_{50}$ mg/kg | Standard Error (-/+) |
| *Amaurobius similis* | 1.92 | 0.61 (1.31 - 2.53) | 9.04 | 2.35 (6.69 - 11.39) | **14.5** | **6.99 (7.51 – 21.49)** | 33.49 | 17.67 (15.82 - 51.16) |
| *Cupiennius coccineus* | 4.8 | 1.24 (3.56 - 6.04) | 10.52 | 6.71 (3.81 - 17.23) | **na** | **na (na - na)** | 26.38 | 10.86 (15.52 - 37.24) |
| *Cupiennius salei* | 2.45 | 0.66 (1.79 - 3.11) | 6.45 | 1.31 (5.14 - 7.76) | **22.66** | **8.26 (14.4 – 30.92)** | 37.2 | 19.93 (17.27 - 57.13) |
| *Eratigena atrica* | **7.66** | **2.53 (5.13 – 10.19)** | 54.63 | 10.66 (43.99 - 65.31) | **21.5** | **25.87 (-4.37 – 47.37)** | 300.54 | 50.23 (250.31 - 350.77) |
| *Heteropoda venatoria* | 1.00 | 0.14 (0.86 - 1.14) | 37.64 | 6.68 (30.96 - 44.32) | **21.58** | **8.32 (13.26 – 29.9)** | 131.53 | 57.16 (74.37 - 188.69) |
| *Larinioides sclopetarius* | 2.78 | 0.46 (2.32 - 3.24) | 15.12 | 6.87 (8.25 - 21.99) | **28.03** | **3.5 (24.53 – 31.53)** | 37.3 | 15.18 (22.12 - 52.48) |
| *Meta menardi* | **0.81** | **0.20 (0.61 – 1.01)** | 12.47 | 2.74 (9.73 - 15.21) | **3.13** | **3.46 (-0.33 – 6.59)** | 75.01 | 58.5 (16.51 - 133.51) |
| *Monocentropus balfouri* | 2.98 | 0.92 (2.06 - 3.9) | 15.18 | 1.66 (13.52 - 16.84) | **12.03** | **5.59 (6.44 – 17.62)** | 8.47 | 6.51 (1.96 - 14.98) |
| *Pholcus phalangioides* | **2.14** | **0.44 (1.7 – 2.58)** | 6.31 | 2.25 (4.06 - 8.56) | **8.59** | **1.28 (7.31 – 9.87)** | 12.62 | 1.56 (11.06 - 14.18) |
| *Phormictopus cancerides* | 16.71 | 3.56 (13.15 - 20.27) | 58.85 | 53.94 (4.91 - 112.79) | **53.0** | **8.39 (44.61 – 61.39)** | 100.34 | 98.5 (1.84 - 198.84) |
| *Piloctenus haematostoma* | 2.66 | 0.57 (2.09 - 3.23) | 5.88 | 1.88 (4 - 7.76) | **25.37** | **18.82 (6.55 – 44.19)** | 19.21 | 9.47 (9.74 - 28.68) |
| *Steatoda nobilis* | **0.05** | **0.028 (0.022 - 0.078)** | 0.72 | 0.20 (0.52 - 0.92) | **1.87** | **1.39 (0.48 – 3.26)** | 7.47 | 6.7 (0.77 – 14.17) |

### Supplementary S5: Supplementary Tables and Model Outputs

**Table M1: Main GLM testing the relationship between $\log_{10}$ of** $LD_{50}$ **(mg/kg) and $\log_{10}$ of** $ED_{50}$ **(mg/kg)** for 24 measures of $LD_{50}$ and 24 measures of $ED_{50}$ (4 hr endpoint) across 12 species in two prey models, *Acheta domesticus* and *Porcellio scaber*. The modes (β) and standard error range are provided for both fixed and random terms, with $\log_{10} ED_{50}$ as the response variable and $\log_{10} LD_{50}$ as the fixed term. Results are deemed significant when p-value < 0.05 and are highlighted in **bold**.

| | β | Standard error | t value | P value |
|---|---|---|---|---|
| **Fixed Terms** | | | | |
| Intercept $_{\log10}$ $ED_{50}$ $_{(mg/kg)}$ | **-0.82** | **0.28** | **-2.93** | **0.01** |
| $Log_{10}$ $LD_{50}$ (mg/kg) | **1.01** | **0.24** | **4.21** | **0.0004** |
| Prey model *Porcellio scaber* | 1.02 | 0.5 | 2.04 | 0.055 |
| $Log_{10}$ $LD_{50}$ (mg/kg): Prey model *Porcellio scaber* | -0.48 | 0.35 | -1.37 | 0.18 |

**Table M2: Main MCMCglmm testing the relationship between $\log_{10}$ of $LD_{50}$ (µl/g) and $\log_{10}$ of $ED_{50}$ (µl/g)** for 55 measures of $LD_{50}$ and 55 measures of $ED_{50}$ across 40 species in eight prey models (seven insects and one arachnid) using data from Friedel and Nentwig 1989, Nentwig *et al.* 1992, Quistad *et al.* 1992, Michálek *et al.* 2019 and Michálek *et al.* 2022 (full citations in main manuscript and S2 dataset). The modes (β) and standard error range are provided for both fixed and random terms, with $\log_{10} ED_{50}$ as the response variable and $\log_{10} LD_{50}$ as the fixed term. The random terms associated with phylogenetic relatedness (phylogeny ($h^2$)), intraspecific variation (species) and residual variation (residual) are also presented. Significant values, which are highlighted in **bold**, are those with 95% of the posterior estimate above or below zero.

| | β | 95% Lower CI | 95% Upper CI |
|---|---|---|---|
| **Fixed Terms** | | | |
| Intercept $_{Log10}$ $ED_{50}$ $_{(µl/g)}$ | **-0.52** | **-0.97** | **-0.12** |
| $Log_{10}$ $LD_{50}$ (µl/g) | **1.10** | **0.81** | **1.39** |
| **Random Terms** | | | |
| Phylogeny ($h^2$) | 0.003 | 0.0002 | 0.37 |
| Species | 0.002 | 0.0002 | 0.26 |
| Residuals | 0.56 | 1.00 | 0.99 |

### Supplementary S5: Supplementary Tables and Model Outputs

**Table S4: Supplementary GLM 1** testing the relationship between $log_{10}$ of $LD_{50}$ (mg/kg) and $log_{10}$ of $ED_{50}$ (mg/kg) for 23 measures of $LD_{50}$ and 23 measures of $ED_{50}$ (1 hr endpoint) across 11 species (*Cupiennius coccineus* excluded), in two prey models, *Acheta domesticus* and *Porcellio scaber*. The modes (β) and standard error range are provided for both fixed and random terms, with $log_{10}$ $ED_{50}$ as the response variable and $log_{10}$ $LD_{50}$ as the fixed term. Results are deemed significant when $p \leq 0.05$ and are highlighted in **bold**.

| | β | Standard error | t value | P value |
|---|---|---|---|---|
| **Fixed Terms** | | | | |
| Intercept $_{log10}$ $ED_{50\ (mg/kg)}$ | **-0.73** | **0.28** | **-2.65** | **0.02** |
| $Log_{10}$ $LD_{50}$ (mg/kg) | **0.95** | **0.23** | **4.06** | **0.0007** |
| Prey model *Porcellio scaber* | **1.27** | **0.49** | **2.57** | **0.02** |
| $Log_{10}$ $LD_{50}$ (mg/kg): Prey model *Porcellio scaber* | -0.57 | 0.34 | -1.66 | 0.11 |

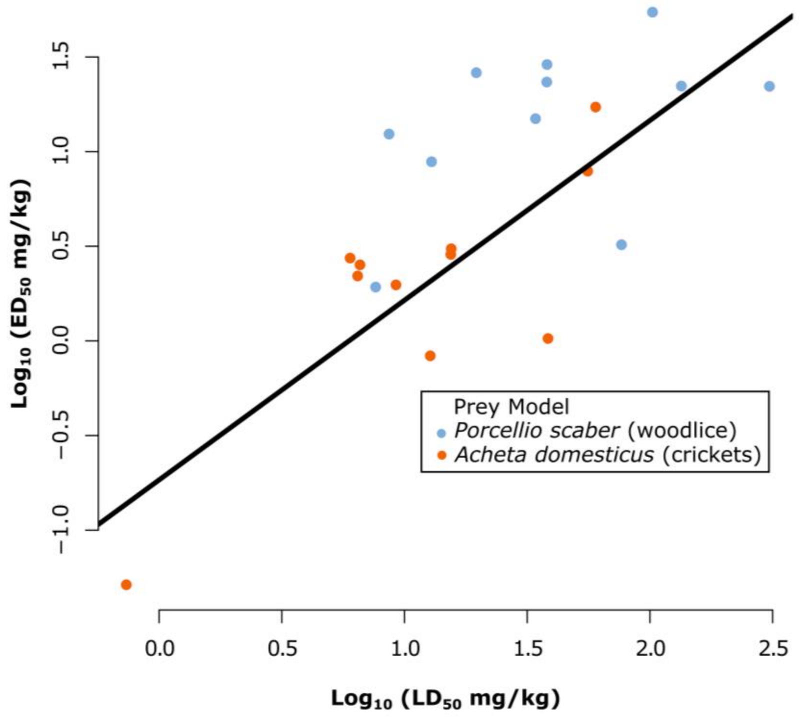


**Figure S1:** Relationship between $log_{10}$ of $LD_{50}$ (mg/kg) and $log_{10}$ of $ED_{50}$ (mg/kg) for 23 measures of $LD_{50}$ and 23 measures of $ED_{50}$ (1 hr endpoint) across 11 species in two prey models, *A. domesticus* (orange) and *P. scaber* (blue). The fitted orange line highlights the significant positive, isometric relationship between $log_{10}$ $LD_{50}$ (mg/kg) and $log_{10}$ $ED_{50}$ (mg/kg)(1 hr) in A. domesticus (β = 0.94, SE = 0.23, p value = 0.0008: Table S3).

#### Supplementary S5: Supplementary Tables and Model Outputs

**Table S5: Supplementary GLM 2** testing the relationship between $log_{10}$ of $LD_{50}$ (mg/kg) and $log_{10}$ of $ED_{50}$ (mg/kg) for 23 measures of $LD_{50}$ and 23 measures of $ED_{50}$ (4 hr endpoint) across 11 species (*Pholcus phalangioides* excluded as their venom was extracted via venom gland extraction), in two prey models, *Acheta domesticus* and *Porcellio scaber*. The modes (β) and standard error range are provided for both fixed and random terms, with $log_{10}$ $ED_{50}$ as the response variable and $log_{10}$ $LD_{50}$ as the fixed term. Results are deemed significant when p ≤ 0.05 and are highlighted in **bold**.

| | **β** | **Standard error** | **t value** | **P value** |
|---|---|---|---|---|
| **Fixed Terms** | | | | |
| Intercept $_{log10\ ED50\ (mg/kg)}$ | **-0.81** | **0.31** | **-2.64** | **0.02** |
| $Log_{10}$ $LD_{50}$ (mg/kg) | **1.00** | **0.25** | **3.93** | **0.001** |
| Prey model *Porcellio scaber* | 1.06 | 0.56 | 1.90 | 0.07 |
| $Log_{10}$ $LD_{50}$ (mg/kg): Prey model *Porcellio scaber* | -0.55 | 0.34 | -1.63 | 0.12 |

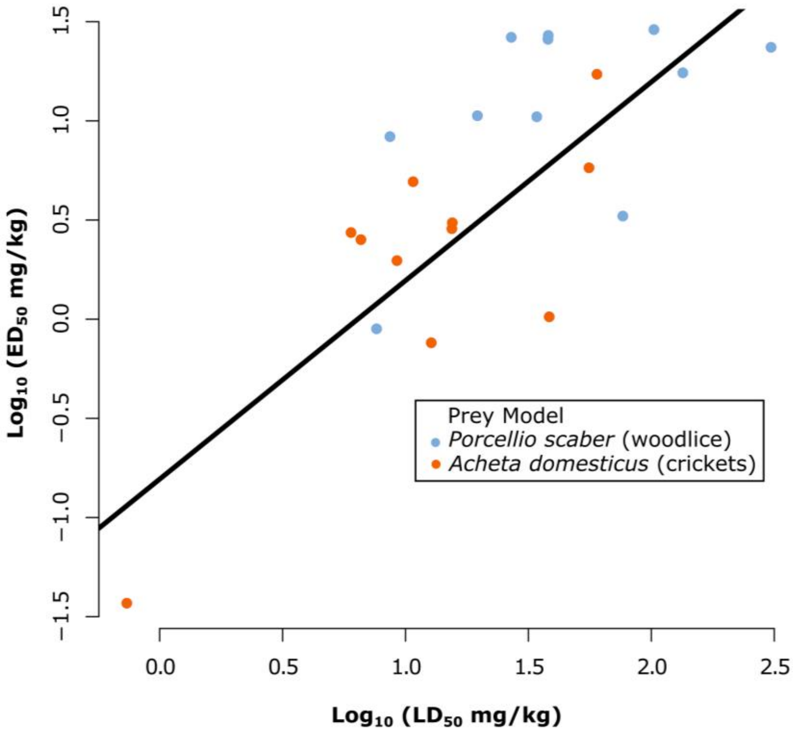


**Figure S2: Relationship between $log_{10}$ of** $LD_{50}$ **(mg/kg) and $log_{10}$ of** $ED_{50}$ **(mg/kg)** for 23 measures of $LD_{50}$ and 23 measures of $ED_{50}$ (4 hr endpoint) across 11 species (*P. phalangioides* excluded) in two prey models, *A. domesticus* (orange) and *P. scaber* (blue). The fitted orange line highlights the significant positive, isometric relationship between $log_{10}$ $LD_{50}$ (mg/kg) and $log_{10}$ $ED_{50}$ (mg/kg)(1 hr) in *A. domesticus* (β = 1.0, SE = 0.25, p value = 0.001: Table S4).

### Supplementary S5: Supplementary Tables and Model Outputs

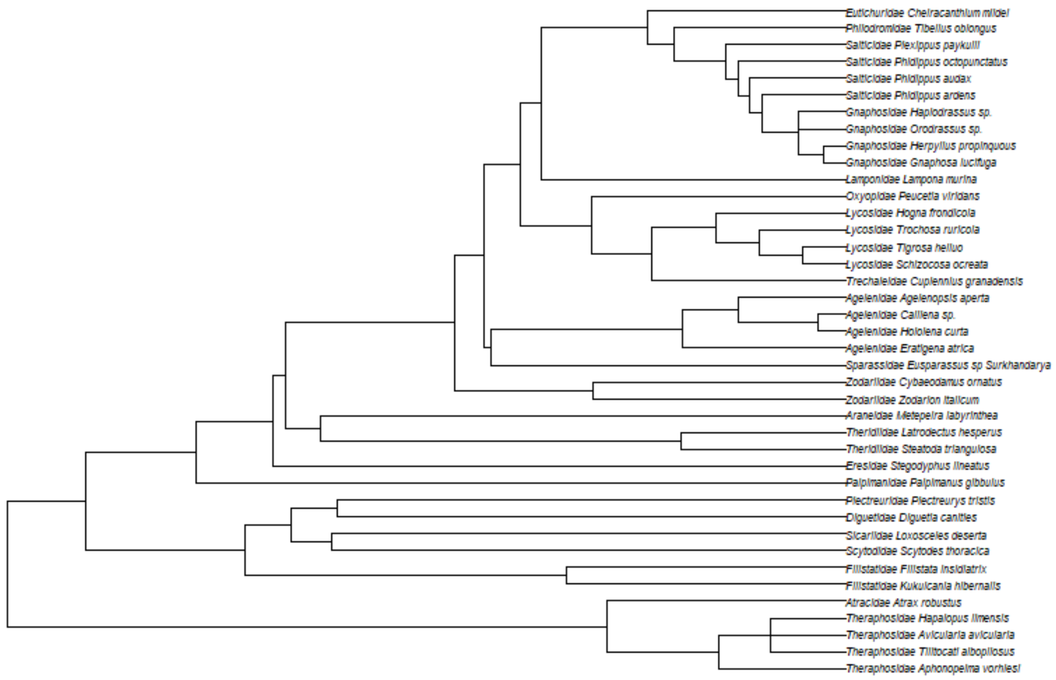


**Figure S3:** Phylogenetic tree produced using Supplementary S4 R studio code for 40 spider species in S2 literature based dataset.
